# Redox Activated Substrates for Enhancing Activatable Cyclopropene Bioorthogonal Reactions

**DOI:** 10.1101/2023.10.13.562277

**Authors:** Wei-Siang Kao, Wei Huang, Yunlei Zhang, Andrea Meyer, Scott T. Laughlin

**Affiliations:** Department of Chemistry, Stony Brook University, Stony Brook, NY 11790, United States

**Keywords:** cyclopropene, bioorthogonal chemistry, click chemistry, cycloaddition

## Abstract

Bioorthogonal chemistry has become a mainstay in chemical biology and is making inroads in the clinic with recent advances in protein targeting and drug release. Since the field’s beginning, a major focus has been on designing bioorthogonal reagents with good selectivity, reactivity, and stability in complex biological environments. More recently, chemists have imbued reagents with new functionalities like click-and-release or light/enzyme-controllable reactivity. We have previously developed a controllable cyclopropene-based bioorthogonal ligation, which has excellent stability in physiological conditions and can be triggered to react with tetrazines by exposure to enzymes, biologically significant small molecules, or light spanning the visual spectrum. Here, to improve reactivity and gain a better understanding of this system, we screened diene reaction partners for the cyclopropene. We found that a cyclopropene–quinone pair is 26 times faster than reactions with 1,2,4,5-tetrazines. Additionally, we showed that the reaction of the cyclopropene–quinone pair can be activated by two orthogonal mechanisms, caging group removal on the cyclopropene and oxidation/reduction of the quinone. Finally, we demonstrated that this caged cyclopropene–quinone can be used as a bioorthogonal imaging tool to label the membranes of cultured cells.

## INTRODUCTION

Since its development in the late 1990s, bioorthogonal chemistry has empowered scientists to conjugate useful chemical groups like fluorophores, biosensors, or drugs to biomolecules like proteins, lipids, and nucleic acids in cells and living organisms^1^. Many new bioorthogonal reactions have been developed, and much of the focus of bioorthogonal reagent design has been on improving reaction rates and reagent stability in the complex biological milieu of living cells^2^. More recently, chemists have designed reactions with an eye towards expanded capabilities, including click-and-release for on-demand drug delivery^3^ and the ability to be “turned on” by exposure to an external stimulus (*e*.*g*., light, enzyme, or metabolites)^4–6^. These activatable bioorthogonal reactions provide the means to control when and where the reaction occurs, promising the ability to limit bioorthogonal reactions to specific cell types or subcellular locations, an incredibly useful feature that has been available to protein-based tools for decades. For example, Fox and coworkers have created a peroxidase- or light and photocatalyst-triggered oxidation reaction that converts dihydrotetrazine into the trans-cyclooctene-reactive s-tetrazine for reaction in hydrogels^7^ and living cells^8^. While this approach takes advantage of the impressive reaction rates for trans-cyclooctene-tetrazine ligations, its use with non-light-based activation modalities has not yet been demonstrated, and the dihydrotetrazines do not quench pendant fluorophores like tetrazines, precluding the use of turn on tetrazine-fluorophores. Devaraj and coworkers have employed a similar strategy to trap the dihydrotetrazine with light labile protecting groups^9^. Other examples include Lin and coworkers work with tetrazoles as light-activated substrates for bioorthogonal ligation via 1,3-dipolar cycloaddition,^10^ and Carrico and coworkers light-triggered Staudinger-Bertozzi ligation.^11^

Our work in this area has focused on the establishment of a modular enzyme-, light-, or metabolite-triggered inverse electron demand Diels– Alder (iEDDA) reaction based on the cyclopropene-tetrazine ligation **(Figure 1A)**.^12–14^ In this approach, we append a caging group to a nitrogen positioned at the C3 carbon of the cyclopropene to prevent cycloaddition with s-tetrazines due to steric hindrance and modulation of the nitrogen’s electron density. An advantage of this work is that the labile caging motif on the cyclopropene scaffold can be easily switched to another caging group within two steps (protecting group removal followed by caging group installation)^12^. We have confirmed that the caged cyclopropene is stable at physiological conditions and in the presence of tetrazines for months without decomposing or reacting with the tetrazine. To begin the reaction, the caging group on the cyclopropene can be removed by exposure to the appropriate stimulus, such as light of the appropriate wavelength or an uncaging enzyme. Once activated, the reactions between uncaged cyclopropene and tetrazines proceed with 2^nd^ order rate constants of 1x10^-1^ – 6x10^-4^ M^-1^s^-1^.

**Figure 1.**
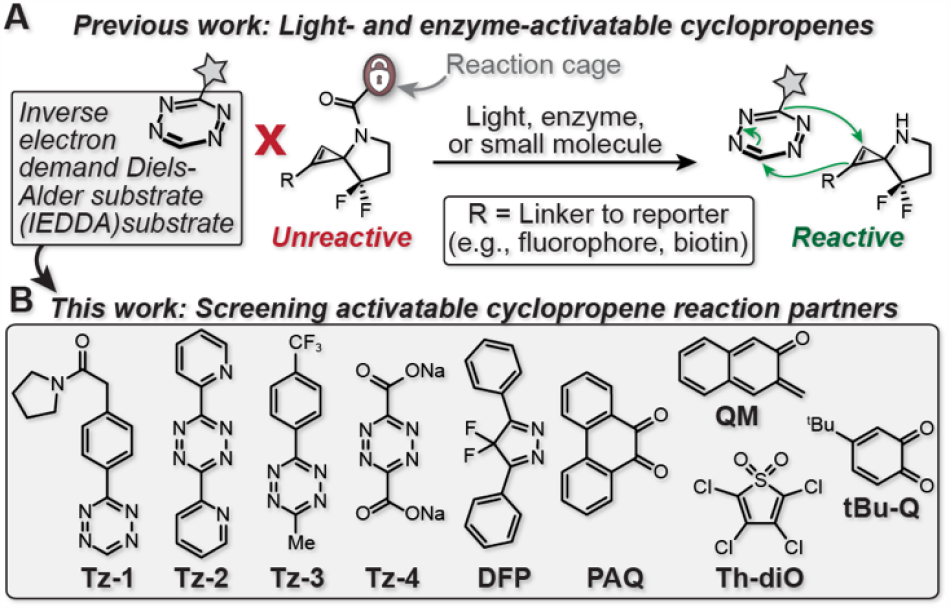
Potential diene substrates and strategies for reaction partner screening in activatable cyclopropene–tetrazine bioorthogonal ligation. **(A)** Previously developed activatable cyclopropene–tetrazine bioorthogonal ligation. The removal of the caging group permits the cycloaddition between cyclopropene and diene substrates like the tetrazine shown. **(B)** Diene substrates evaluated for use with activatable cyclopropenes in this work.

While we have shown that the activatable cyclopropene-tetrazine ligation can be used for bioconjugation in cells^12^, optimizing the reaction kinetics to achieve parity with best-in-class bioorthogonal reagents will further improve its utility. Such optimizations can be accomplished through modifications to the cyclopropene component of the reaction, which we have explored with the creation of a ketone-containing cyclopropene^15^ and continues to be of interest to our lab. Here, however, we take a different approach. We evaluate alternative iEDDA reaction partners in an effort to optimize the reaction rates of this class of activatable reaction, and, potentially, to evaluate additional activation strategies.

Essentially, we sought diene substrates that have been previously reported for other iEDDA reactions to examine whether they react with the activatable cyclopropene shown in **Figure 1A**. We began our search with modified s-tetrazines, as they are popular dienes for iEDDA reactions due to their rapid reaction rates, which can exceed 10^4^ M^-1^s^-1^ with strained alkenes^16,17^ and alkynes.^18,19^ To select compounds, we drew inspiration from recent work by Fox and coworkers exploring the commonly used preparation through a silver-mediated Liebeskind–Srogl cross-coupling.^20^ This report showed that tetrazines decorated with functional groups to improve iEDDA reactivity and/or tetrazine stability could be synthesized in fewer steps than the traditional methods for tetrazine preparation.^19,21,22^ In an effort to enhance the reactivity toward the dienophile, the activatable cyclopropene in our case, we selected tetrazines with strong electron withdrawing groups (**Tz-3** and **Tz-4, Figure 1B**) to evaluate whether the cycloaddition reactivity can be enhanced relative to the commonly-used tetrazines in our group and elsewhere, **Tz-1** and **Tz-2 (Figure 1B**).

In addition to tetrazines, we sought other diene substrates to evaluate as candidates for improved reactivity, each of which is shown in **Figure 1B**. These additional substrates included a 4,4-difluoro-3,5-diphenyl-4*H*-pyrazole (**DFP, Figure 1B**) recently reported by Raines and coworkers^23^, with a 5.2 M^-1^s^-1^ reaction rate with bicyclononyne (**BCN**), almost two times faster than tetrazine/BCN. A 9,10-phenanthrenequinone (**PAQ, Figure 1B**), developed by Zhang and coworkers for visible light-triggered bioorthogonal photoclick cycloadditions. Here, 405 nm light or two-photon excitation triggers the ligation with an alkene within one minute, with rate constants up to 2.76 M^-1^s^-1^.^24^ An *o*-naphthoquinone methide developed by Popik and coworkers (**QM, Figure 1B**),^25,26^ which can be generated *in situ* by treating 3-(hydroxymethyl)-2-naphthol (**NQMP**) with 300 nm light (**Figure S4A**), suggesting the possibility of dual methods for controlling the bioorthogonal ligation with activatable cyclopropenes. A thiophene dioxide (**Th-diO, Figure 1B**) developed by Wang and coworkers as a prodrug to release sulfur dioxide (SO_2_) in cells after undergoing an iEDDA reaction with cyclooctyne with rates approaching 0.26 M^-1^s^-1^ in methanol at room temperature.^27^ And, finally, a tert butyl quinone (**tBu-Q, Figure 1B**), brought to our attention by reports from Hest and coworkers.^28,29^ **tBu-Q**-based substrates are particularly interesting due to the potential for quinone generation *in situ* through oxidization of the corresponding catechol, raising the potential for activating the reaction by exposure to an oxidase enzyme, oxidizing agents, or light.^30^

In the following report, we describe the synthesis and evaluation of the above candidate IEDDA reaction partners for the activatable cyclopropenes. First, we determine their reactivity, stability, and synthetic accessibility. We examine the reactivity with both the caged and uncaged forms of an activatable cyclopropene (**H-CP-FF-Ben, Figure 2A**) as well as the stability of each diene molecule in physiological conditions. We identify one promising new reaction partner, the quinone (**tBu-Q, Figure 2A**) and then evaluate its reaction with activatable cyclopropenes in the presence or absence of an oxidant and cyclopropene activation stimulus by confocal microscopy on the membranes of cultured cells.

**Figure 2.**
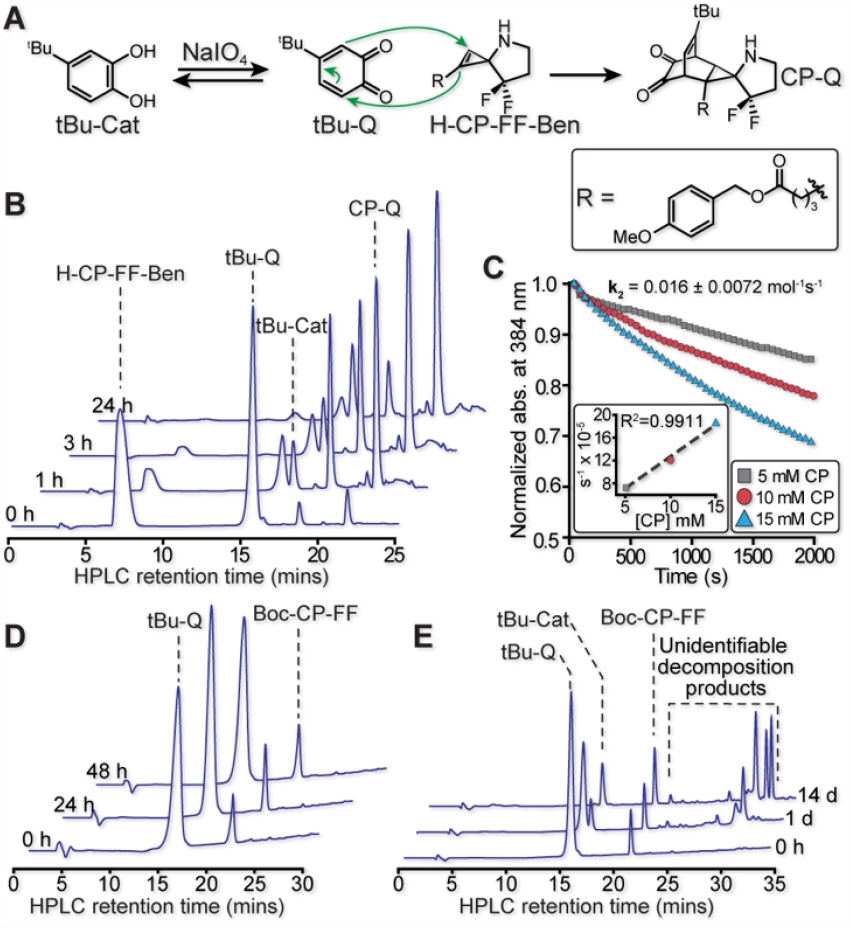
General bioorthogonal pairs screening process for caged- /uncaged-cyclopropene and dienes. **(A)** The cycloaddition reaction scheme of uncaged difluoro cyclopropene **H-CP-FF-Ben** and tertbutyl quinone **tBu-Q (B)** HPLC analyses of the reactivity between cyclopropene **H-CP-FF-Ben** and quinone **tBu-Q (C)** Kinetic measurement of cycloaddition rate **(D)** HPLC analyses of no reaction between caged difluoro cyclopropene **Boc-CP-FF** and tert-butyl quinone **tBu-Q** in acetonitrile **(E)** HPLC analyses of no reaction between caged cyclopropene **Boc-CP-FF** and Quinone **tBu-Q** in physiological conditions.

## RESULTS AND DISCUSSION

To investigate the reactivity of an activatable cyclopropene with the candidate dienes from **Figure 1B** we used HPLC to monitor the reaction progression. First, we evaluated the modified tetrazines **Tz-3** and **Tz-4**. 1,2,4,5-Tetrazine derivatives are widely used in bioorthogonal reactions due to their fast reaction rates with cyclopropenes and trans-cyclooctenes and good selectivity at low concentrations. In previous studies, the mono-substituted tetrazine **Tz-1** and di-substituted tetrazine **Tz-2** had a second-order reaction rate of approximately 0.0006 M^-1^s^-1^ and 0.00017 M^-1^s^-1^ with difluoro-cyclopropene **H-CP-FF**.^13,14^ Consistent with literature reports, we found that mono-substituted tetrazine was more reactive but less stable than di-substituted tetrazine. We predicted that introducing an electron-withdrawn group on the tetrazine would increase the reactivity towards the cyclopropene dienophile, since previous reports have suggested that the reactivity toward dienophiles may be enhanced with more electron deficient substitutions to tetrazine.^31–34^. Accordingly, we chose **Tz-3** and **Tz-4**, which have been previously reported,^35–37^ to test electron deficient tetrazines in the context of the activatable cyclopropene system.

To begin evaluating these tetrazines, we mixed **Tz-3** and **H-CP-FF-Ben** at 5 mM in 1:1 MeCN:PBS and monitored the reaction for reagent stability and product formation by HPLC. After 2 hours, we observed no change in the HPLC trace, indicating no, or very slow, reactivity with the reagent pair (**Figure S1**). Extending the reaction time to 3 days showed a small decrease in HPLC peak volume for both reagents and the formation of two small broad peaks whose identity we were unable to determine by mass spectrometry (**Figure S1**).

Moving on to **Tz-4**, early studies by Bradley and coworkers found that the Diels-Alder reaction between **Tz-4** and allyl glycidyl ether in DMSO reaches 100% conversion in ∼1.5 hours.^38^ Later, Koert and coworkers showed that **Tz-4** reacts with cyclooctyne in room temperature CH_2_Cl_2_, reaching completion in ∼40 seconds.^39^ However, we observed no product formation in our HPLC assay with the activatable cyclopropene **H-CP-FF-Ben**. When we tested the tetrazine **Tz-4** with cyclopropene, we found that the **Tz-4** had poor stability and decomposed quickly in aqueous buffer, producing no identifiable products according to ^1^H NMR analysis of the reaction mixture.^34,40^

We continued our exploration of other candidate diene reaction partners for and **H-CP-FF-Ben** with **DFP** (**Figure 1B**). Previous work by Raines and coworkers found a reaction rate between **DFP** and **BCN** of 5.2 M^-1^s^-1^, two times faster than **BCN** with diphenyl tetrazine in physiological conditions.^23^ To test **DFP**’s reactivity toward the cyclopropene **H-CP-FF-Ben**, we performed an HPLC analysis in MeCN: PBS=1:1, monitoring reaction times as long at 24 h. Unfortunately, under these conditions we did not observe either the formation of products or decomposition of starting materials **(Figure S2)**.

Next, we moved on to evaluate **PAQ** and **QM** (**Figure 1B**) as potential reaction partners with **H-CP-FF-Ben**. Previous reports have shown that the cycloaddition between alkene and **PAQ**, or **QM**, can be initiated by light in a wavelength range of 250 to 350 nm.^24,41–43^ Intrigued by the possibility of dual activation modalities (*i*.*e*., light-based formation of the **PAQ** or **QM** and light- or enzyme-based activation of the cyclopropene), we investigated their reaction with **H-CP-FF-Ben**. We first examined the reaction between **H-CP-FF-Ben** and **PAQ** in the dark for 1 h. As expected for this duration in the dark, we observed no decomposition or product formation by HPLC (**Figure S3**). Unfortunately, when the mixture of **H-CP-FF-Ben** and **PAQ** was irradiated with 365 nm or 405 nm light for 1 minute to activate PAQ, we observed decomposition of cyclopropene **H-CP-FF-Ben (Figure S3)**. We hypothesize that light activated PAQ, which forms a radical intermediate upon light exposure^24^, reacted with the cyclopropene to form a mixture of unidentifiable side products. We then evaluated **QM**’s reactivity with **H-CP-FF-Ben**. In these studies, we found, consistent with literature reports^25^, that the quinone methide intermediate has a short halflife (τ ≈ 7 ms) in aqueous conditions, reducing back to the starting material, 3-(hydroxymethyl)-2-naphthol (**NQMP**), thus precluding its reaction with **H-CP-FF-Ben (Figure S4)**.

We then moved our attention to **Th-diO** (**Figure 1B**), which has been shown in previous reports to be a versatile substrate that can be used as an iEDDA diene.^44^ Previous research has demonstrated that thiophene dioxide derivatives can undergo Diels–Alder reactions with alkyne or alkene in organic solvent at various temperature conditions, followed by the release of sulfur dioxide to form the aromatic ring.^45,46^ Thiophene dioxide derivatives share a similar two-step mechanism with tetrazine derivatives, where the cheletropic reaction occurs after cycloaddition between diene and dienophile. In the first step, the cycloaddition rate depends on the energy gap differences between the diene’s LUMO and the dienophile’s HOMO. The conformation of the product and the tendency to form the aromatic ring are the two major driving forces in the second step. To improve the reactivity and solubility in the reaction media (acetonitrile: PBS in 1:1), we selected tetrachlorothiophene dioxide **Th-diO** as the diene to analyze the ligation rate with cyclopropene. Our HPLC analyses followed a similar method to that used for the other candidate dienes described above, unfortunately, once again showing no product formation **(Figure S5)**.

Finally, we concluded our search for new diene reaction partners for **H-CP-FF-Ben** by exploring their reaction with 1,2-quinones. 1,2-quinones have been reported to undergo a concerted [4+2] cycloaddition reaction with cyclooctynes in organic solvents.^28^ The 1,2-quinone can be generated by oxidizing the corresponding catechol with sodium periodate or tyrosinase to form the 1,2-quinone from phenol in the presence of oxygen (**Figure 2A**).^47^ Due to their fast reaction rate with cyclooctyne, 1,2-quinone substrates have been considered as potential bioorthogonal diene substrates with reaction rates up to 500 M^-1^s^-1^.^28^ Here, we investigated the reactivity of cyclopropene **H-CP-FF-Ben** with tert-butyl quinone **tBu-Q** in a 1-to1 ratio in 50% acetonitrile/PBS at room temperature **(Figure 2B)**. Our HPLC analysis showed that the starting material cyclopropene **H-CP-FF-Ben** and tert-butyl quinone **tBu-Q**, gradually decreased in intensity, while new peaks appeared corresponding to tert-butyl catechol **tBu-Cat**, an unknown product, potentially an intermediate, that initially increases and then decreases over time, and the cycloaddition product **CP-Q**, with retention times (*t*_*R*_) = 16.4 min, 18.8 min, and 21.9 min, respectively **(Figure 2B)**. We collected the unknown peak and attempted to characterize it as an intermediate or decomposition product by mass spectrometry and NMR, but we were unable to identify it due, in part, to its short lifetime (It degraded on the order of 24 hours into compounds we were unable to identify by mass spectrometry). After a day, **H-CP-FF-Ben** was consumed and only trace **tBu-Q** remained, with the majority of the reaction mixure being composed of the expected ligation product **CP-Q** and reduced **tBu-Q** in the form of **tBu-Cat**. We evaluated the reaction rate of this cyclopropene–quinone reaction by monitoring the disappearance of tert-butyl quinone **tBu-Q** using its characteristic absorbance at 384 nm and found a second-order rate constant of 0.016 +/- 0.0072 M^-1^s^-1^, which is 26 times faster than **H-CP-FF-Ben**’s reaction with **Tz-1 (Figure 2C)**.

Having identified a new reaction partner for this activatable cyclopropene, we next sought to confirm the cyclopropene’s ability to be caged by N-modification and activated by N-modification removal. Previous studies have shown these activatable cyclopropenes can have their reactivity caged through N-modification with a diverse array of carbamate-linked removable protecting groups.^48^ To evaluate reactivity caging with **tBu-Q**, we tested the reaction between a Boc-caged difluoro cyclopropene **Boc-CP-FF** and **tBu-Q** by HPLC. In these experiments, we found no reaction between these two reagents in either organic solvent **(Figure 2D)** or 50% acetonitrile/PBS buffer for over 2 weeks **(Figure 2E)**. However, we did observe **tBu-Q**’s reduction to the catechol in the 50% acetonitrile/PBS buffer conditions over 1–2 days, consistent with our results in **Figure 2B** and literature reports of quinone reduction in aqueous conditions. ^49,50^ We further confirmed that the new peaks are from the decomposition of tert-butyl quinone **tBu-Q** in a control experiment with only **tBu-Q** under the same 50% aqueous/organic conditions, indeed finding that the quinone is reduced to the corresponding catechol **(Figure S6)**. We did not observe this **tBu-Q** reduction in organic solvents **(Figure 2D)**.

These findings show that the reaction of quinone **tBu-Q** with uncaged activatable cyclopropene **H-CP-FF-Ben** is 26-times faster than that of the previous faster partner **Tz-1**. However, the spontaneous aqueous reduction of **tBu-Q** to the catechol presents an intriguing opportunity to create a dual activatable reagent, restricting the quinonecyclopropene reaction to cells or organelles with oxidizing environments (to generate **tBu-Q**) and further controlling reaction timing or location by light- or enzyme-based activation of a caged **H-CP-FF-Ben**.

To begin exploring the potential of this dual oxidant- and light-based reaction activation, we evaluated the activatable cyclopropene and quinone for the labeling of cellular lipids. Bioorthogonal lipid labeling has provided valuable information on lipid metabolism, localization, and lipid post-translational modifications through the incorporation of either metabolic clickable precursors or chemically synthesized lipids with clickable handles.^6,21,51,52^ Nevertheless, there have been few reports for controllable bioorthogonal lipid labeling, which limits the use of bioorthogonal lipid labeling for studying lipid function in specific cell types or tissues. To achieve cell-selective lipid labeling, we linked the activatable difluoro cyclopropene scaffold with a sulforhodamine B fluorophore and caged it using the light-labile group Nvoc (**Nvoc-CP-Rho**) to prevent the ligation reaction from occurring prior to light exposure (**Figure 3A**). Additionally, we conjugated a catechol substrate (**tBu-Q** precursor) to a DOPE phospholipid **(C-DOPE) (Figure 3A**, see **SI** and **SI Scheme 7** for synthetic details and compound characterization). As one of the most abundant lipids in cell membranes,^53^ DOPE has been modified with fluorophores and successfully used in labeling both the plasma membrane^54^ and membranes of intracellular organelles, including the ER, mitochondria, Golgi, and late endosomes.^51^

**Figure 3.**
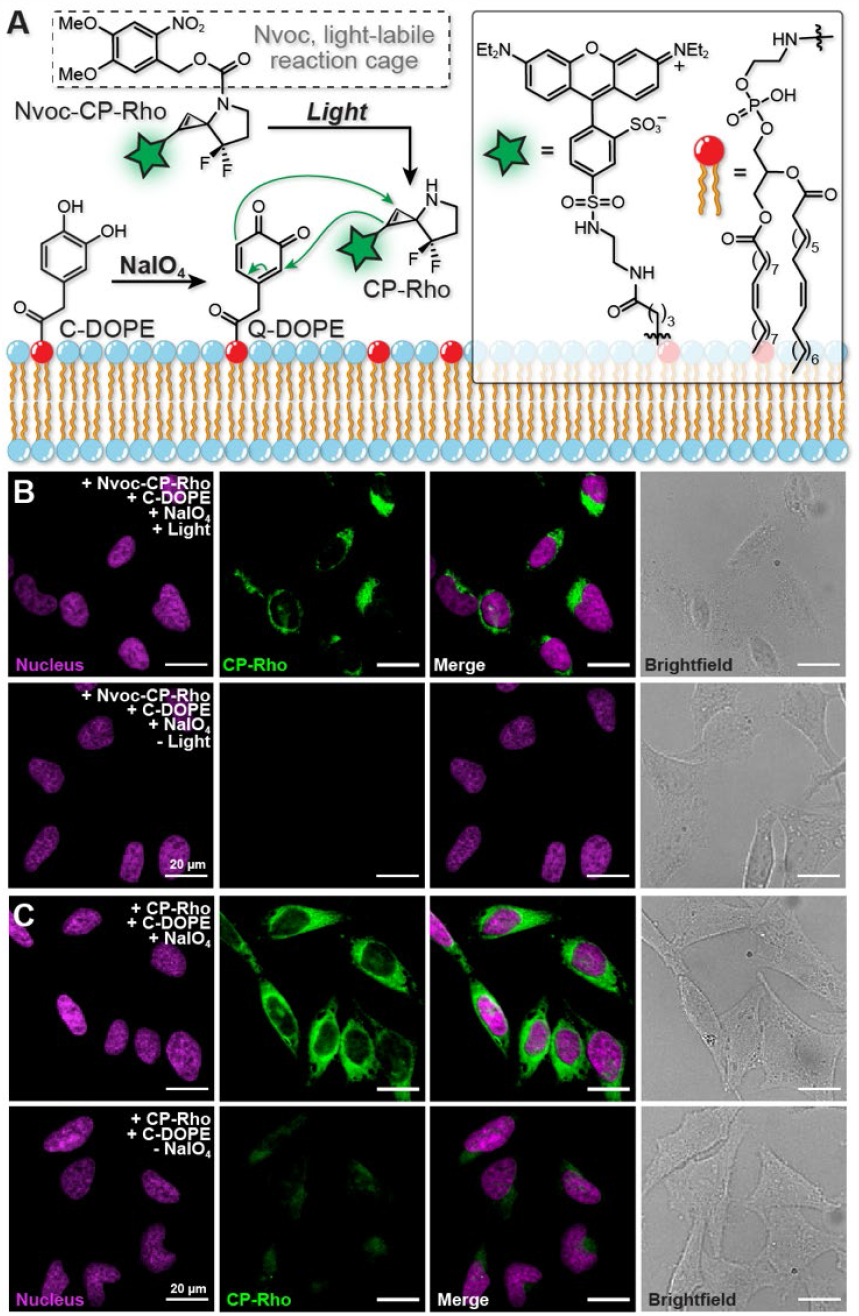
The light- and oxidation-activatable cyclopropene-quinone reaction for cell imaging. **(A)** Schematic description of the activatable cyclopropene-quinone system for imaging labeled lipids in cells. A catechol-labeled phospholipid (**C-DOPE**) was incubated with cells for incorporation into cellular membranes. An activatable cyclopropene caged by the light-labile Nvoc group and labeled with sulforhodamine B (**Nvoc-CP-Rho**) was then delivered to cells. Exposure to light removes Nvoc to activate the cyclopropene (**CP-Rho**) and addition of NaIO_4_ converts C-DOPE into the reactive quinone-conjugated phospholipid (**Q-DOPE**), allow the ligation reaction to proceed. **(B)** Light- and oxidation-activated ligation for cellular imaging. Top row, the ligation reaction was initiated after 365 nm light irradiation for 1 minute. Prominent CP-Rho labeling can be observed (Green). Bottom row, without 365 nm light exposure the caged Nvoc-CP-Rho remains intact, producing no observable CP-Rho labeling. **(C)** Evaluating the requirement for oxidant exposure. Top row, with the addition of NaIO_4_, the provided CP-Rho produced prominent flruoescent labeling of cell membranes (Green). Bottom row, without the addition of NaIO4, only minimal labeling with CP-Rho is observed. Magenta nucleus fluorescence represents Hoescht 33342 labeled nuclei. Images are displayed as maximum intensity projections of a confocal microscopy z stack. All scale bars: 20 µm.

We incubated Hela cells with **C-DOPE** for 1 hour to allow it to incorporate into the cell membranes. The now **C-DOPE**-labeled cells were washed three times with PBS (1 mL) and fixed with 4% PFA in PBS (1 mL) for 20 minutes at room temperature. Next, we incubated the **C-DOPE**-labeled cells with **Nvoc-CP-Rho** (10 µM in PBS) and the oxidant sodium periodate (NaIO^4^, 1 mM in H_2_O) to convert the catechols to quinones *in situ*. To initiate the ligation reaction, we irradiated the samples at 365 nm (1 min, 0.3W) to remove the caging group on the cyclopropene, allowing the liberated **CP-Rho** to react with **Q-DOPE** integrated into the Hela cell’s membranes, or kept cells in the dark as control **(Figure 3A)**.

We then evaluated these cells by confocal microscopy. The results showed no observable fluorescence labeling on the Hela cell membranes when the cyclopropene was caged in the presence of sodium periodate (**Figure 3B**, bottom panels). Once the sample was exposed to 365 nm light to release **CP-Rho**, the ligation occurred, producing fluorophore labeling of the membranes in HeLa cells **(Figure 3B**, top panels). We further examined whether adding an oxidant or light exposure would affect the ligation outcomes. Without the oxidant, the reaction produced only very modest cell membrane labeling **(Figure 3C**, bottom panels), consistent with there being a small concentration of quinone produced in the absence of an added oxidizing agent. Adding NaIO_4_ produced the expected intense fluorescent labeling of Hela cell membranes (**Figure 3C**, top panels). We also performed additional control experiments to verify that this system can only be activated by applying both light and oxidant **(Figure S7)**. These results suggest that this catechol/quinone and activatable cyclopropene system can serve as a dual-activatable reaction for imaging cells based on the presence of an oxidizing environment, which could report on cells or organelles with oxidizing environments with the added spatiotemporal control provided by light- or enzyme-based activation of the activatable cyclopropene.

In conclusion, we have screened several classes of diene candidates in an effort to identify an activatable cyclopropene reaction partner with improved reaction rates. We found that the activatable cyclopropene **H-CP-FF-Ben** had a faster reaction rate with quinone substrates than the best-in-class tetrazines, but the quinones have poor stability in aqueous environments due to their tendency to reduce to the catechol. The addition of an oxidant served to preserve a sufficient pool of reactive quinone. The requirement of an oxidizing environment for quinine stability and efficient reactions with the activatable cyclopropene have the potential to provide a dual, oxidant and light, method for reaction activation. Finally, we used this oxidant and light activatable reaction to image the presence of modified lipids embedded in the cell membranes of Hela cells using confocal fluorescence microscopy.

## ACKNOWLEDGMENT

We thank Nan Wang and the Center for Advanced Study of Drug Action (CASDA) at Stony Brook University for mass spectrometry resources and analysis, Dr. Francis Picart and Dr. Fang Liu for NMR instrumental support and analysis.

## Supporting Information

## List of abbreviations

Calcd: Calculated
DCM: Dichloromethane
DIPEA: *N,N*-diisopropylethylamine
DMF: Dimethylformamide
DOPE: Dioleoylphosphatidylethanolamine
ESI: Electrospray ionization
EtOAc: Ethyl acetate
EtOH: Ethanol
HATU: 1-[Bis(dimethylamino)methylene]-1H-1,2,3-triazolo[4,5-b]pyridinium 3-oxid hexafluorophosphate
HBSS: Hank’s balanced salt solution
HMPA: Hexamethylphosphoramide
HOBt: Hydroxybenzotriazole
HPLC: High performance liquid chromatography
HRMS: High resolution mass spectrometry
LAH: Lithium aluminum hydride
LiHMDS: Lithium bis(trimethylsilyl)amide
MeCN: Acetonitrile
MeOH: Methanol
NEM: *N*-Ethylmaleimide
MHz: Megahertz
NPPOC: 3’-nitrophenylpropyloxycarbonyl
PBS: Phosphate-buffered saline
PFA: Paraformaldehyde
R_f_: Retention factor
R_t_: Retention time
THF: Tetrahydrofuran
TLC: Thin layer chromatography
UV: Ultraviolet

## General materials and methods

All chemical reagents were of analytical grade, obtained from commercial suppliers, and used without further purification unless otherwise specified. Reactions were monitored by thin-layer chromatography on pre-coated glass TLC plates (Analtech UNIPLATE^TM^ silica gel HTF w/ organic binder, 250 μm thickness, with UV 254 indicator) or by LC/MS (Agilent 6110 LC-MSD, direct-injection mode, 1–20 μL, ESI). TLC plates were visualized by UV illumination or developed with either potassium permanganate stain (KMnO_4_ stain: 1.5 g KMnO_4_, 10 g K_2_CO_3_, and 1.25 mL of 10% NaOH dissolved in 200 mL H_2_O) or ninhydrin stain (1.5 g ninhydrin dissolved in 100 mL of 1-butanol and 3 mL of conc. AcOH). Flash chromatography was carried out using Sorbtech, 60 Å, 40–63 µm or Millipore 60 Å, 40–63 µm silica gel according to the procedure described by Still.^1^ HPLC was performed using a Shimadzu HPLC (FCV-200AL) equipped with an Agilent reversed phase Zorbax Sb-Aq C18 column (4.6 × 250 mm or 21.2 × 250 mm) fitted with an Agilent stand-alone prep guard column. NMR spectra (^1^H and ^13^C) were obtained using a 300, 400, 500, or 700 MHz Bruker spectrometer and analyzed using Mestrenova 14.0. ^1^H and ^13^C chemical shifts (δ) were referenced to residual solvent peaks. Following residual solvent peaks were chosen: (for ^1^H NMR) CDCl_3_, 7.2600 ppm; CD_3_OD, 3.3100 ppm; (CD_3_)_2_SO, 2.5000 ppm (for ^13^C NMR) CDCl_3_, 77.1600 ppm; CD_3_OD, 49.0000 ppm; (CD_3_)_2_SO, 39.5200 ppm; D_2_O, 4.7900 ppm. Further, the following abbreviations were used to define ^1^H NMR peaks: s, singlet; d, doublet; t, triplet; q, quartet; m, multiplet. Low-resolution electrospray ionization (ESI) and High-resolution electrospray ionization (ESI) mass spectra were obtained at the Stony Brook University Center for Advanced Study of Drug Action (CASDA) Mass Spectrometry Facility with an Agilent 6110 Single Quad LC/MSD and Bruker Impact II LC-UV-TOF spectrometer respectively. All the *N*-Boc protected (**Boc-CP-FF** and **Boc-CP-Ket**), *N*-Boc deprotected cyclopropene (**CP-FF** and **CP-Ket**), C1 p-methoxybenzyl cyclopropene (**H-CP-FF-Ben**), and tetrazine **Tz1** were synthesized as described recently by us.^2–4^ The synthesis of tetrachlorothiophene dioxide (**Th-diO**) was synthesized according to work described by Wang et al.^5^ 35mm glass bottom microwell dish (20 mm glass diameter) were purchased from MatTek Corporation (Ashland, MA). The Hela cell line was a gift from Peter Tonge (ATCC CCL-2^TM^). Hoechst 33342 was purchased from Thermo Fisher Scientific (Boston, MA). All cell images were acquired using a confocal microscope (Zeiss Axio Examiner. D1 modified with an Andor Differential Scanning Disk confocal unit) using a Zeiss water immersion 40X NA 1 Plan-APOCHROMAT or 20X NA 0.50 N-ACHROPLAN objective (Zeiss) and analyzed using ImageJ to produce maximum intensity Z projections.

## HeLa cell culture

HeLa cells were maintained in high-glucose Dulbecco’s Modified Eagle Medium (DMEM) supplemented with 10% FBS and 1% Gibco^TM^ Antibiotic-Antimycotic at 37 °C in air supplemented with 5% CO_2_. Cells were passaged on alternating days when they reached approximately 90% confluency.

## HeLa cell imaging

Cells were seeded at 100,000 cells per 35 mm glass-bottom culture dish (MatTek) and grown for 48 h to 60–80% confluency. To begin labeling, the cells were washed twice with 37 °C PBS (1 mL), incubated with 100 µM catechol-DOPE **(C-DOPE)** in HBSS at 37 °C for 1 h, washed three times with PBS (1 mL), and fixed with 4% PFA in PBS (1 mL) for 20 minutes at room temperature. After fixation, the cells were washed three times with PBS (1 mL) and then treated with 40 mM *N*-Ethylmaleimide (NEM) in PBS (1.5 mL) at 37 °C for 24h to prevent cross-reaction between thiols and quinones. After 24 h, the NEM solution was aspirated, and the cells were washed with PBS (1 mL) prior to treatment with 1 mM sodium periodate in water (1 mL), or water (1 mL) without oxidant, at room temperature for 1 h. After the 1 h incubation, the cell bathing media was aspirated, and the cells were incubated with either 10 µM **H-CP-FF-Rho** or 10 µM of **Nvoc-CP-FF-Rho** in PBS (1 mL, 37 °C) for 14 h. The cells were washed three times with PBS (1 mL) and then submitted to a final wash of 48 hours in PBS (2 mL) at room temperature to diminish the nonspecific labeling. After 48 h, the PBS was aspirated, and the cell nuclei were labeled with 1 µg/mL Hoescht 33342 in PBS (1 mL) for 20 minutes, followed by one PBS wash (1 mL). Then PBS (2 mL) was added, and the cells were imaged by confocal microscopy. For light-controlled labeling with **Nvoc-CP-FF-Rho**, the cells were illuminated (or left in the dark for the control) with a 365 nm mounted LED (Thorlabs, 0.36W) for 1 min with a gentle shaking of the dish after the addition of **Nvoc-CP-FF-Rho**.

**Figure S1:**
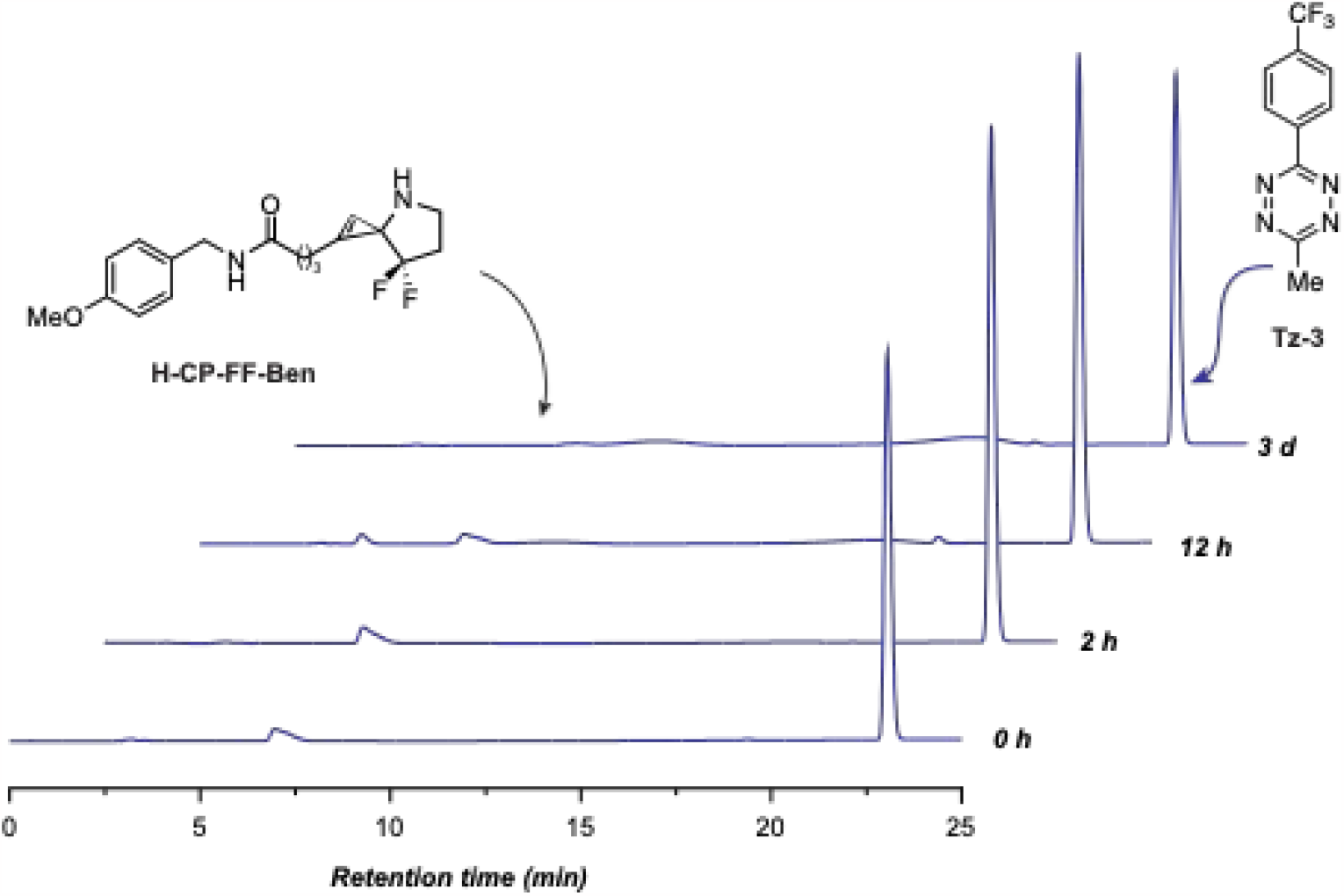
HPLC analyses of no reaction between uncaged cyclopropene H-CP-FF-Ben and 3-methyl-6-(4-(trifluoromethyl)phenyl)-1,2,4,5-tetrazine Tz-3. A reaction mixture consisting of uncaged cyclopropene **H-CP-FF-Ben** (5.0 mM) and 3-methyl-6-(4-(trifluoromethyl)phenyl)-1,2,4,5-tetrazine **Tz-3** (5.0 mM) in 200 µL of 1:1 MeCN/PBS (pH = 7.4, ThermoFisher Scientific# 10010023) was incubated in the dark at rt. At each designated time point (0 h, 2 h, 12 h, 3 d) a 20 µL aliquot was withdrawn from the reaction mixture, diluted to 200 µL using 1:1 MeCN/H_2_O and subjected to HPLC (30–100% MeCN + 0.1% TFA over 25 mins, flow rate = 1 mL/min). Each HPLC peak was confirmed to be the expected, unreacted reagent by MS analyses.

**Figure S2:**
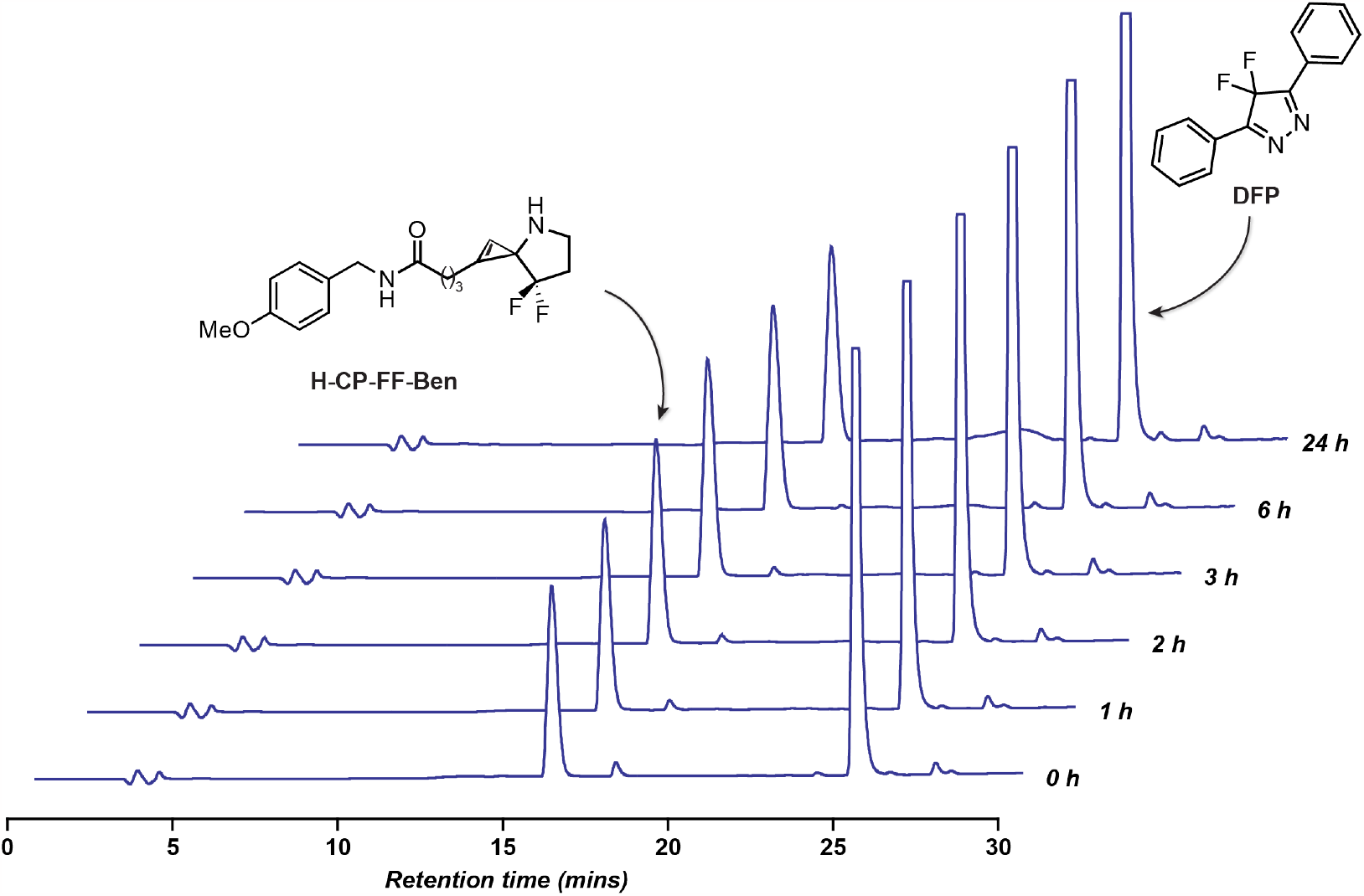
HPLC analyses of no reaction between uncaged cyclopropene H-CP-FF-Ben and 4,4-difluoro-3,5-diphenyl-4*H*-pyrazole (DFP). A reaction mixture consisting of uncaged cyclopropene **H-CP-FF-Ben** (5.0 mM) and 4,4-difluoro-3,5-diphenyl-4H-pyrazole **(DFP)** (5.0 mM) in 200 µL of 1:1 MeCN/PBS (pH = 7.4, ThermoFisher Scientific# 10010023) was incubated in the dark at rt. At each designated time point (0 h, 1 h, 2 h, 3 h, 6 h, 24 h) a 20 µL aliquot was withdrawn from the reaction mixture, diluted to 100 µL using 1:1 MeCN/H_2_O and subjected to HPLC (20–100% MeCN + 0.1% TFA over 25 mins, flow rate = 1 mL/min). Each HPLC peak was confirmed to be the expected, unreacted reagent by MS analyses.

**Figure S3:**
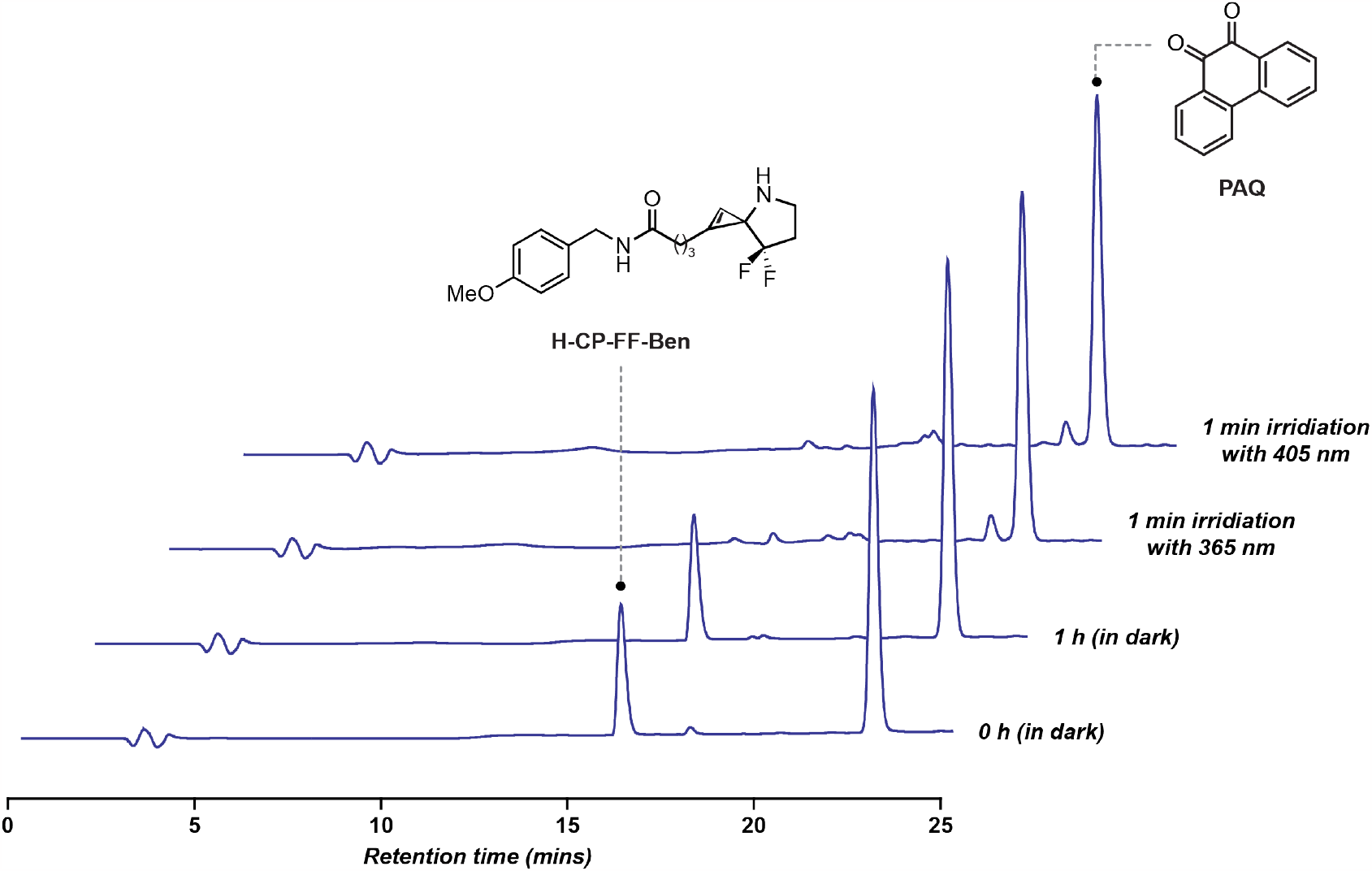
HPLC analyses of no reaction between uncaged cyclopropene H-CP-FF-Ben and 9,10-Phenanthrenequinone PAQ. A reaction mixture consisting of uncaged cyclopropene **H-CP-FF-Ben** (5.0 mM) and 9,10-Phenanthrenequinone **PAQ** (5.0 mM) in 200 µL of 1:1 MeCN/PBS (pH = 7.4, ThermoFisher Scientific# 10010023) was incubated in the dark at rt. At each designated time point (0 h, 1 h, 1 min irradiation with 365 nm, and 1 min irradiation with 405 nm) a 20 µL aliquot was withdrawn from the reaction mixture, diluted to 100 µL using 1:1 MeCN/H_2_O and subjected to HPLC (20–100% MeCN + 0.1% TFA over 25 mins, flow rate = 1 mL/min). Each HPLC peak was confirmed to be the expected, unreacted reagent by MS analyses.

**Figure S4:**
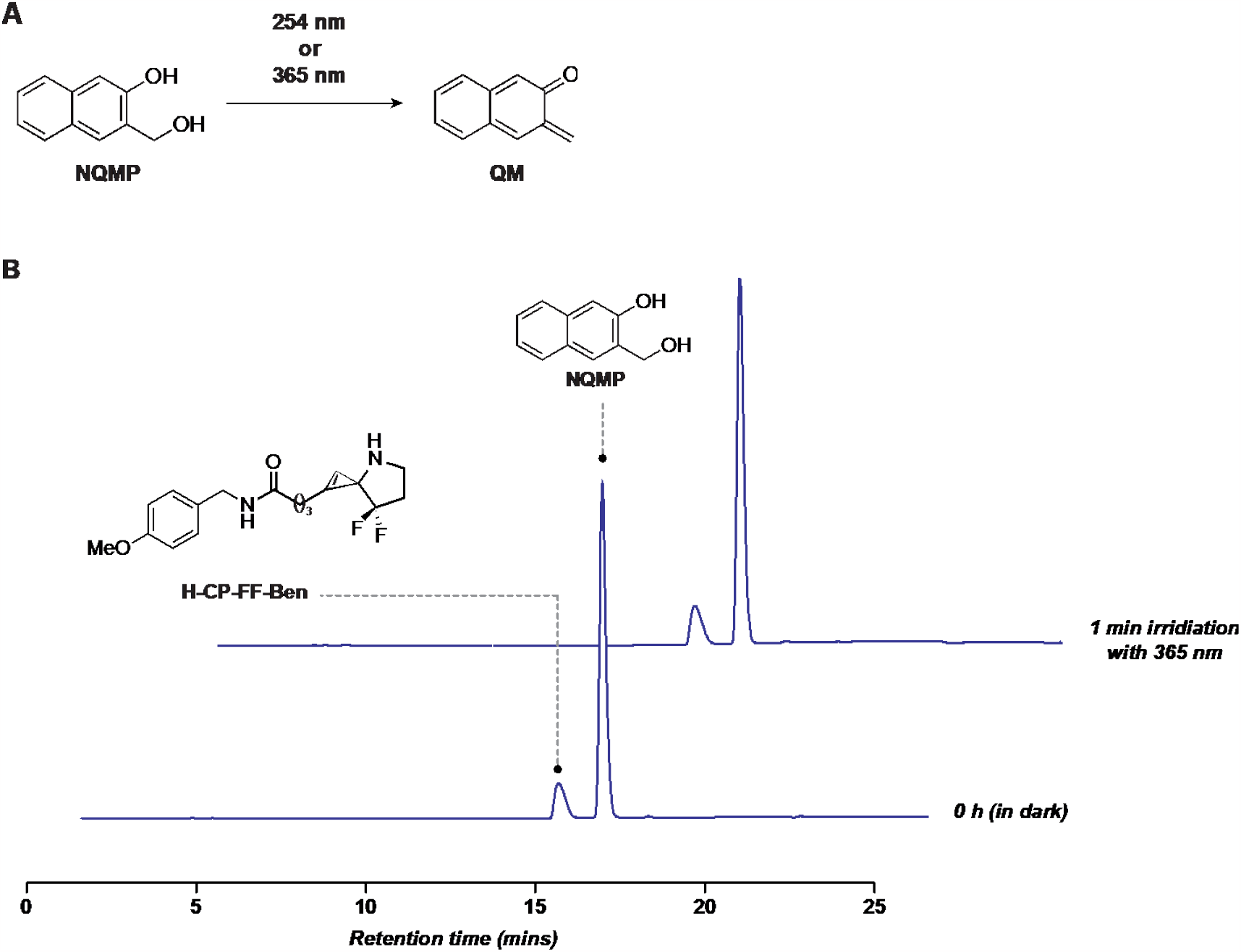
HPLC analyses of no reaction between uncaged cyclopropene H-CP-FF-Ben and 3- (hydroxymethyl)-2-naphthol NQMP. **(A)** *o*-naphthoquinone methide **QM** was generated by treating 3- (hydroxymethyl)-2-naphthol **NQMP** with 365 nm or 254 nm of light. **(B)** A reaction mixture consisting of uncaged cyclopropene **H-CP-FF-Ben** (5.0 mM) and 3-(hydroxymethyl)-2-naphthol **NQMP** (5.0 mM) in 200 µL of 1:1 MeCN/PBS (pH = 7.4, ThermoFisher Scientific# 10010023) was incubated in the dark at rt. At each designated time point (0 h and 1 min irradiation with 365 nm) a 40 µL aliquot was withdrawn from the reaction mixture, diluted to 200 µL using 1:1 MeCN/H_2_O and subjected to HPLC (20–100% MeCN + 0.1% TFA over 25 mins, flow rate = 1 mL/min). Each HPLC peak was confirmed to be the expected, unreacted reagent by MS analyses.

**Figure S5:**
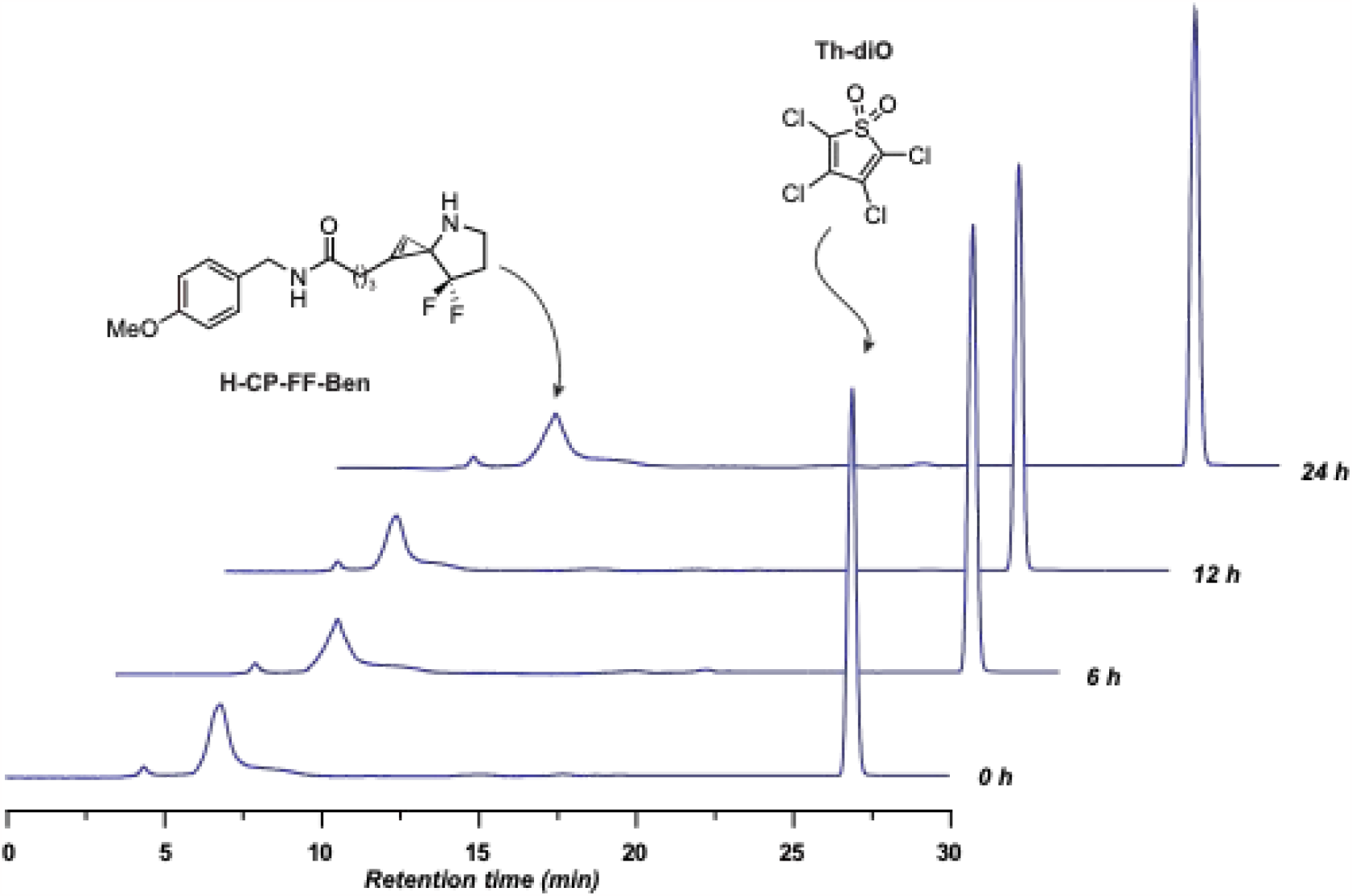
HPLC analyses of no reaction between uncaged cyclopropene H-CP-FF-Ben and thiophene dioxide Th-diO. A reaction mixture consisting of uncaged cyclopropene **H-CP-FF-Ben** (5.0 mM) and thiophene dioxide **Th-diO** (5.0 mM) in 200 µL of 1:1 MeCN/PBS (pH = 7.4, ThermoFisher Scientific# 10010023) was incubated in the dark at rt. At each designated time point (0 h, 6 h, 12 h, 24 h) a 40 µL aliquot was withdrawn from the reaction mixture, diluted to 200 µL using 1:1 MeCN/H_2_O and subjected to HPLC (30–100% MeCN + 0.1% TFA over 25 mins, flow rate = 1 mL/min). Each HPLC peak was confirmed to be the expected, unreacted reagent by MS analyses.

**Figure S6:**
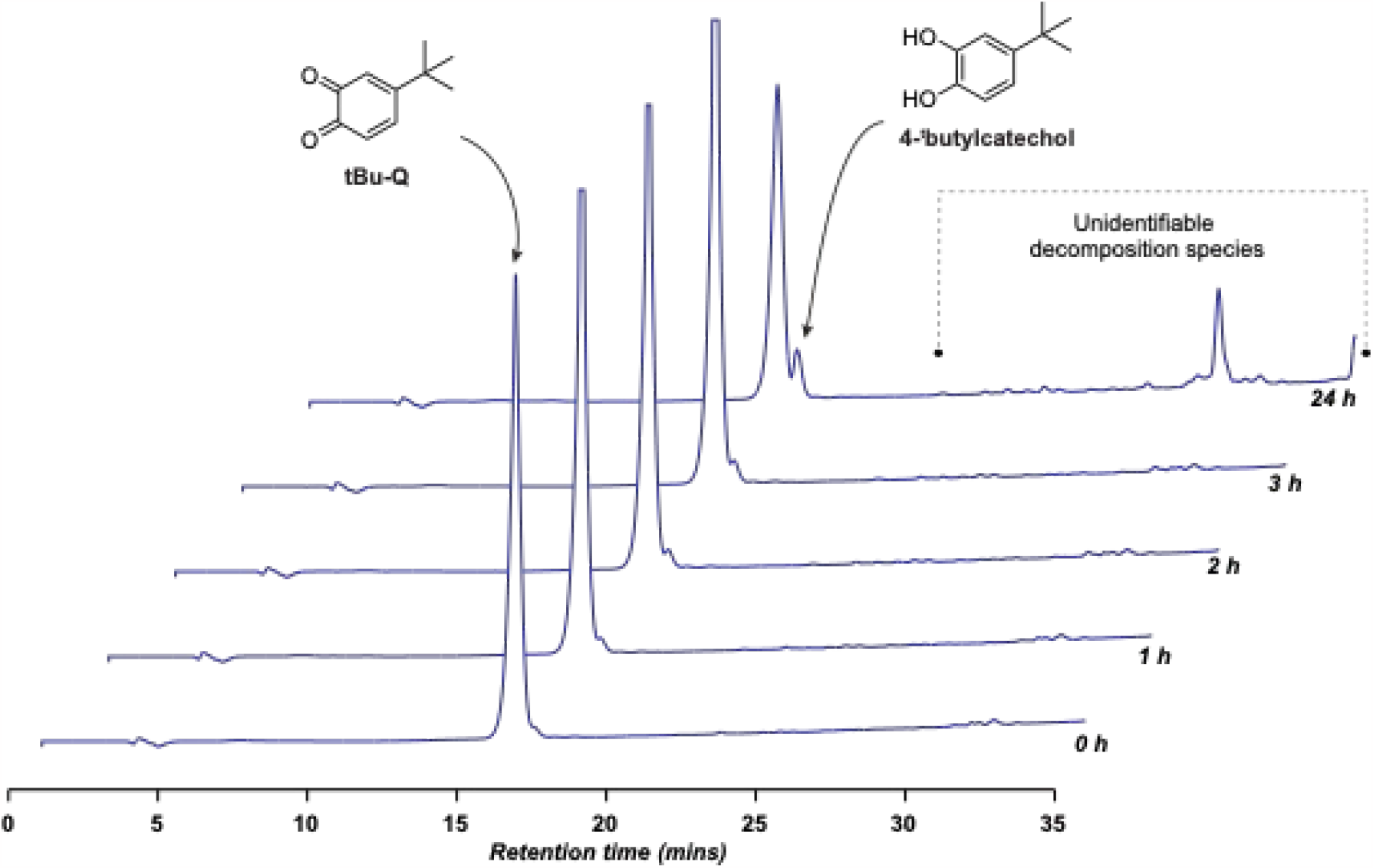
HPLC analyses of reduction of 4-tertbutylquinone tBu-Q into 4-tertbutylcatechol in 1:1 MeCN/PBS. A reaction mixture consisting of 4-tertbutylquinone **tBu-Q** (5.0 mM) in 500 µL of 1:1 MeCN/PBS (pH = 7.4, ThermoFisher Scientific# 10010023) was incubated in the dark at rt. At each designated time point (0 h, 1 h, 2 h, 3 h, 24 h) a 50 µL aliquot was withdrawn from the reaction mixture, diluted to 100 µL using 1:1 MeCN/H_2_O and subjected to reversed phase HPLC (30–100% MeCN + 0.1% TFA over 25 mins, flow rate = 1 mL/min). Each HPLC peak was analyzed by MS. 4-tertbutylquinone **tBu-Q** was found to reduce back to 4-tertbutylcatechol in aqueous buffer at neutral pH. Catechol **DA1-1** further decomposes into unidentifiable species under these reaction conditions.

**Figure S7:**
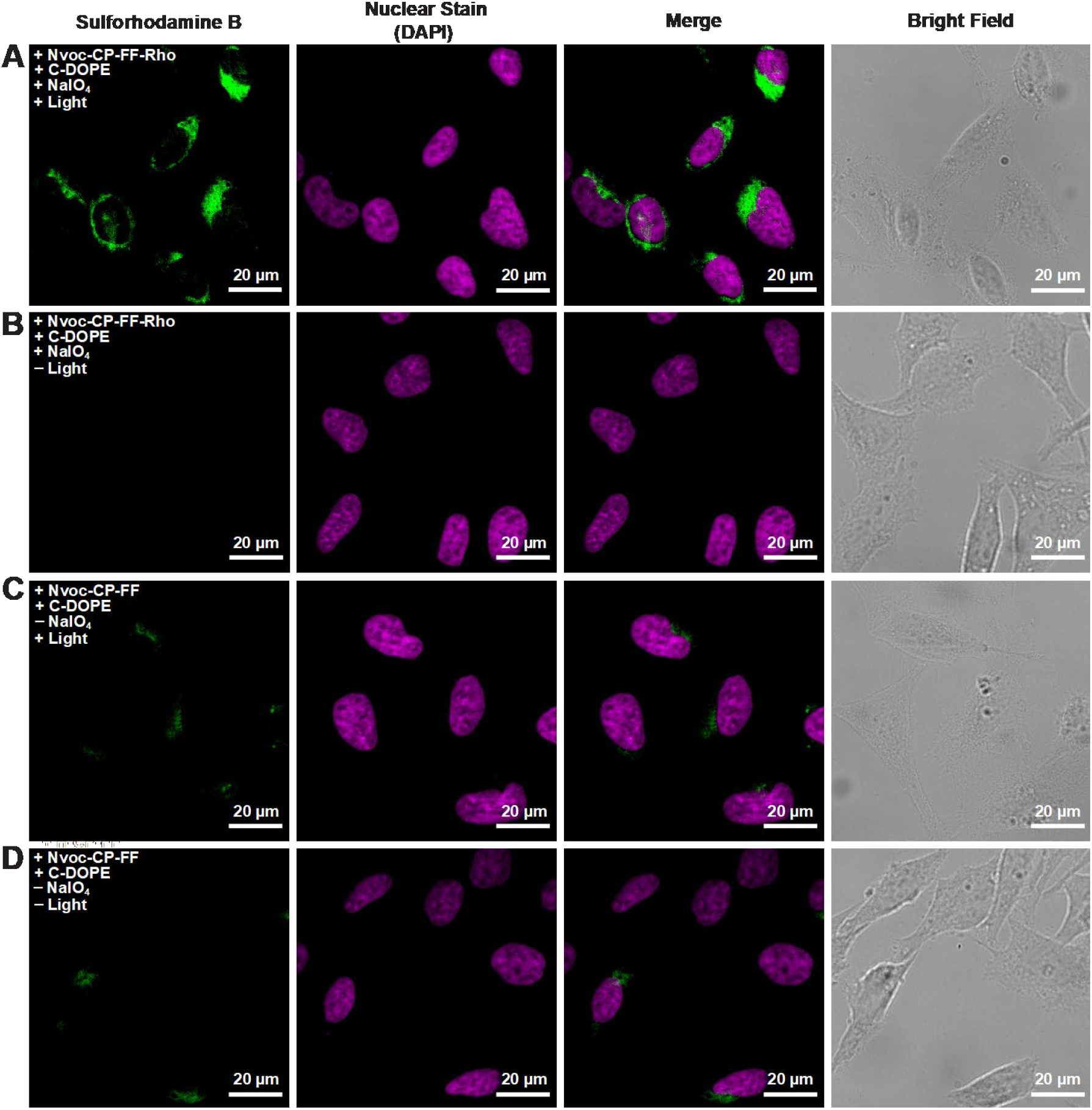
Comparison of the bioorthogonal labeling efficiency of Nvoc-CP-FF and C-DOPE with and without light and sodium periodate. (**A**) With 365 nm light exposure and oxidant. (**B**) Without 365 nm light exposure, with oxidant. (**C**) With 365 nm light exposure, without oxidant. (D) Without 365 nm light exposure and without oxidant. (Scale bars 20 µm)

**Scheme 1:**
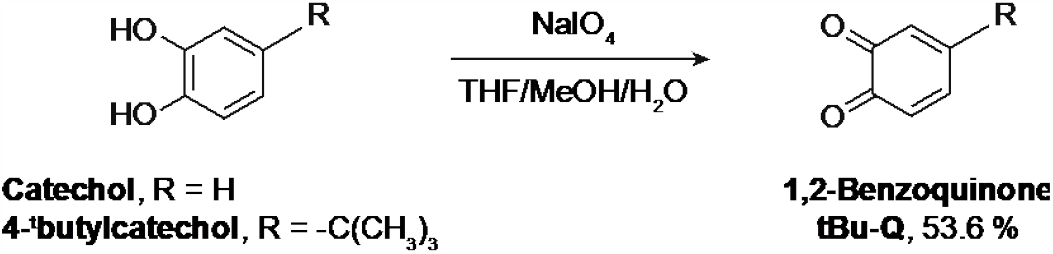
Methods for synthesis of compounds tBu-Q.

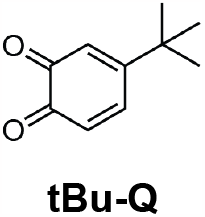

Compound **tBu-Q** was synthesized based on slight modification of a previously described protocol.^6^ To a solution of 4-^t^butylcatechol (2.0 g, 166.22 mmol, 1 eq) in THF/MeOH/H_2_O (130 mL/ 12 mL/ 30 mL) at rt was added sodium periodate (2.57 g, 213.89 mmol, 1 eq) in one portion (Methanol was added to completely solubilize the starting material). The deep red/brown suspension was stirred for 1 h, concentrated *in vacuo*, and diluted with DCM/H_2_O. The DCM layer was dried over anhydrous Na_2_SO_4_, concentrated *in vacuo* and purified by two successive flash chromatography (25 g SiO_2_, 10% EtOAc/Hexanes) to obtain **tBu-Q** as a brown solid (1.06 g, 53.6%). R_f_ = 0.55 (30% EtOAc/Hexanes, visualized w/ UV). The NMR spectra was consistent with the previously reported literature spectra of this compound.^6 1^H NMR (400 MHz, CDCl_3_): δ = 7.10 (dd, *J* = 10.4, 2.4 Hz, 1H), 6.21 (d, *J* = 10.4 Hz, 1H), 6.07 (d, *J* = 2.4 Hz, 1H), 1.07 (s, 9H). ^13^C NMR (101 MHz, CDCl_3_): δ = 180.00, 179.98, 162.06, 140.03, 129.02, 123.28, 77.16, 53.39, 35.32, 27.41. COSY NMR (400 MHz, CDCl_3_, attached) were obtained under same conditions. MS (ESI): Calcd C_10_H_13_O_2_ [M+H]^+^: 165.1, found: 165.1.

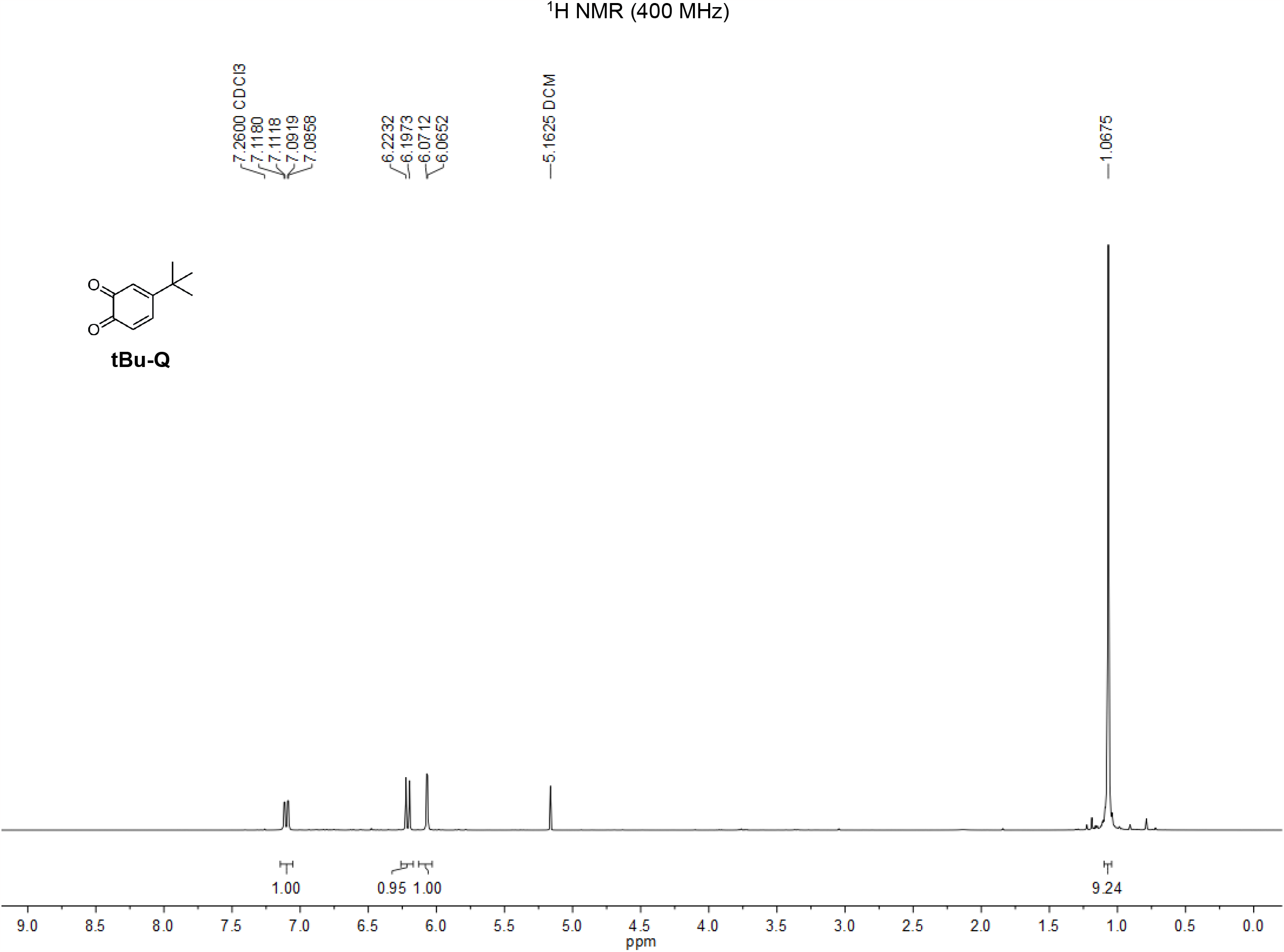

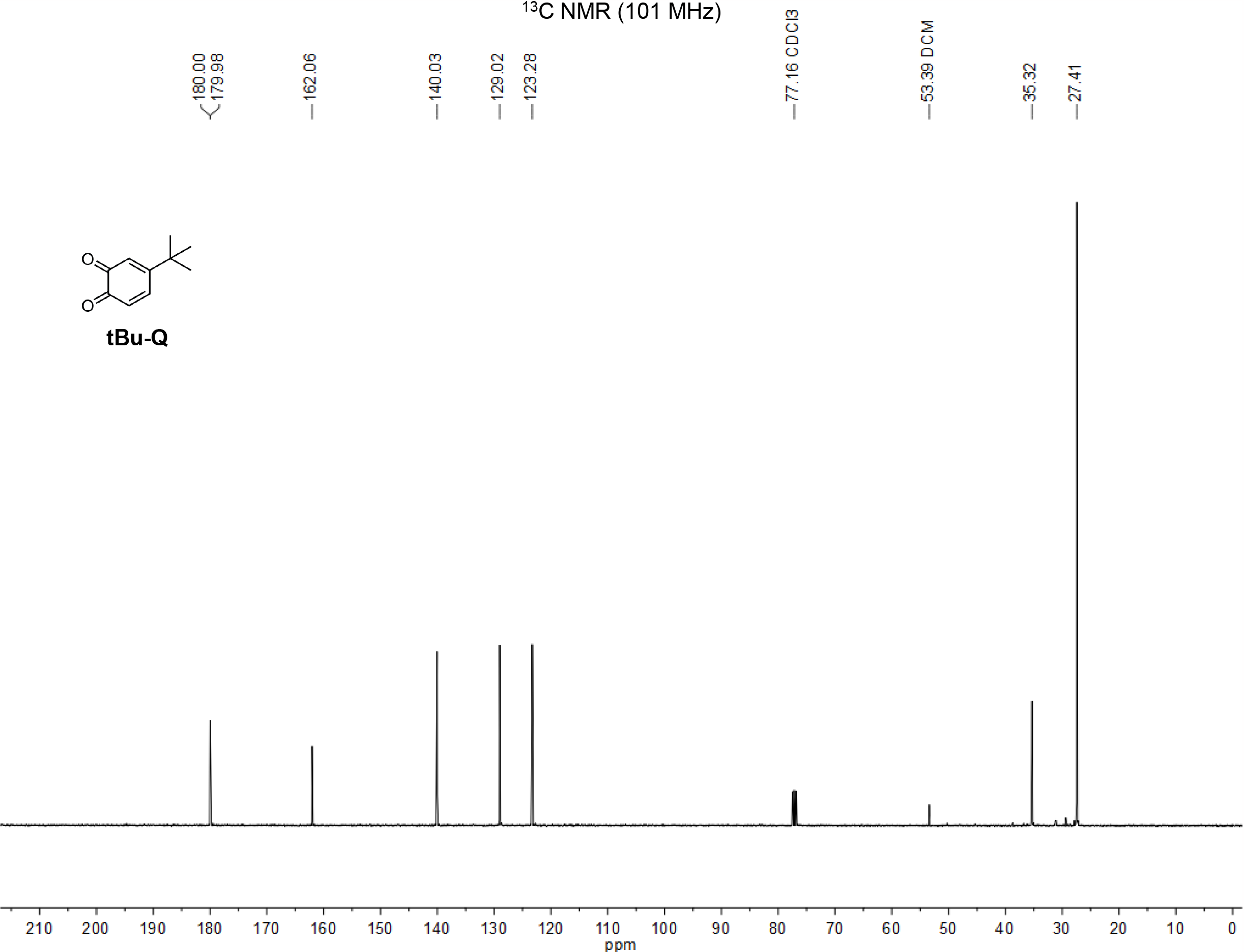

**Scheme 2:**
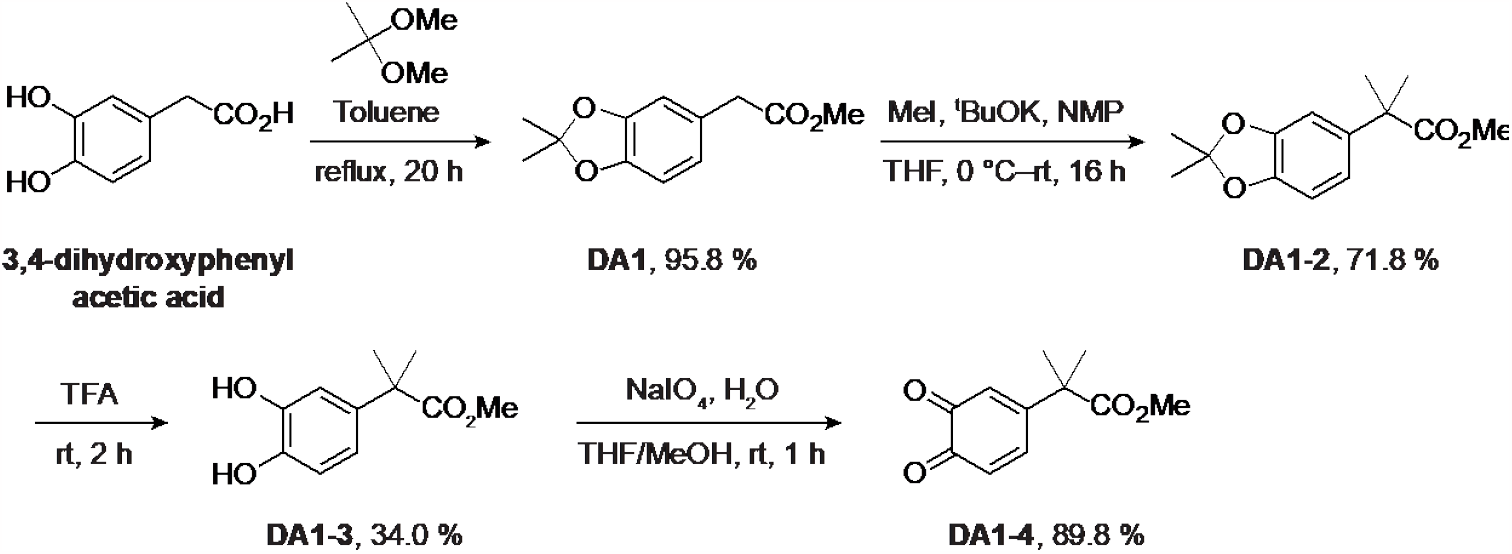
Methods for synthesis of compounds DA1, DA1-2, DA-1-3 and DA1-4.

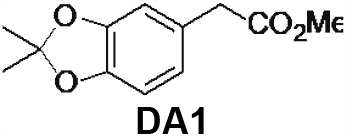

To 3,4-dihydroxyphenylacetic acid (1 g, 5.95 mmol, 1 eq) in Toluene (50 mL) was added 2,2-dimethoxypropane (4.37 mL, 35.67 mmol, 6 eq) and *p*-toluenesulfonic acid (0.11 g, 0.59 mmol, 0.1 eq). The mixture was refluxed for 16 hours. The reaction mixture was washed with water two times, then washed with brine, and the organic phase was collected. The organic layer was further dried over anhydrous Na_2_SO_4_, filtered through filter paper, and concentrated *in vacuo*. The crude product was purified by flash column chromatography on silica gel using 10% EtOAc/hexanes (v/v) to obtain compound **DA1-9** (1.27 g, 95.8%). R_f_ = 0.48 (10% EtOAc/Hexanes, visualized w/ UV). ^1^H NMR (400 MHz, CDCl_3_) δ = 6.68-6.66 (m, 3H), 3.69 (s, 3H), 2.52 (s, 2H), 1.66 (s, 6H). ^13^C NMR (100 MHz, CDCl_3_): δ = 172.50, 147.84, 146.81, 127.08, 121.97, 118.21, 109.71, 108.31, 52.25, 41.07, 26.11. LRMS (ESI): Calcd C_12_H_15_O_4_ [M+H]+: 223.1, Found: 223.1.

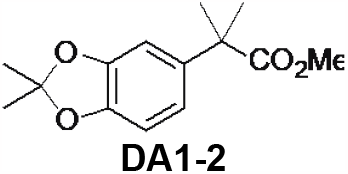

To compound **DA1** (0.59 g, 2.65 mmol, 1 eq) in THF/NMP (50 mL, v:v=1:1) was added ^t^BuOK (1.30 g, 10.6 mmol, 4 eq). The mixture was allowed to stir at 0 ^o^C for 10 min, and iodomethane (0.66 ml, 10.6 mmol, 4 eq) was slowly added into the solution for an additional 2 hours. The reaction mixture was extracted using DCM and water. The combined organic layer was further washed by water and brine, dried over anhydrous Na_2_SO_4_, filtered through filter paper, and concentrated *in vacuo*. The crude product was purified by flash column chromatography on silica gel using 10% EtOAc/hexanes (v/v) to obtain compound **DA1-2** (0.48 g, 71.8%). R_f_ = 0.32 (10% EtOAc/Hexanes, visualized w/ UV). ^1^H NMR (400 MHz, CDCl_3_) δ = 6.75-6.73 (m, 2H), 6.67-6.64 (m, 1H), 3.65 (s, 3H), 1.66 (s, 6H), 1.53 (s, 6H). ^13^C NMR (100 MHz, CDCl_3_): δ = 177.53, 147.71, 146.29, 138.18, 118.18, 109.58, 107.97, 106.62, 52.43, 46.39, 26.98, 26.15. HRMS (ESI): Calcd C_14_H_19_O_4_ [M+H]^+^: 251.1283, Found: 251.1240.

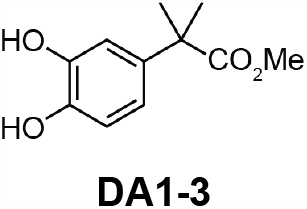

To compound **DA1-2** (0.21 g, 0.84 mmol, 1 eq) was cooled 0 ^o^C at and TFA (5 mL) was added. The mixture was allowed to warm up gradually to room temperature and stirred for 2 hours. The volatiles were removed *in vacuo*. The crude product was purified by flash column chromatography on silica gel using 50% EtOAc/hexanes (v/v)) to obtain compound **DA1-3** (0.06 g, 34.0%). R_f_ = 0.39 (50% EtOAc/Hexanes, visualized w/ UV and KMnO_4_). ^1^H NMR (400 MHz, CDCl_3_) δ = 6.87-6.86 (d, *J* = 2 Hz, 1H), 6.80-6.73 (m, 2H), 5.38-5.29 (br, 2H), 3.65 (s, 3H), 1.53 (s, 6H). ^13^C NMR (100 MHz, CDCl_3_): δ = 178.82, 143.79, 142.88, 137.28, 118.06, 115.37, 113.42, 52.76, 46.13, 26.64. HRMS (ESI): Calcd C_11_H_14_O_4_ [M+Na]^+^: 233.0790, Found: 233.0779

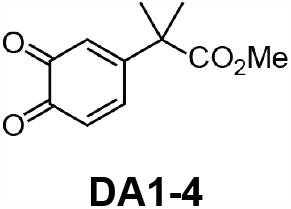

To compound **DA1-3** (59 mg, 0.28 mmol, 1 eq) in THF/MeOH (v:v=1:0.4, 3 mL) was added aqueous NaIO_4_ (60 mg, 0.28 mmol in 2.5 mL water). The mixture was allowed to stir at room temperature for 1 hour. The methanol was removed *in vacuo*. The reaction mixture was extracted using DCM and water. The combined organic layer was further washed by water and brine, dried over anhydrous Na_2_SO_4_, filtered through filter paper and concentrated *in vacuo*. The crude product was purified by flash column chromatography on silica gel using 30% EtOAc/hexanes (v/v) to obtain compound **DA1-4** (52 mg, 89.8%). R_f_ = 0.17 (30% EtOAc/Hexanes, visualized w/ UV and KMnO_4_). ^1^H NMR (700 MHz, CDCl_3_) δ = 6.96-6.94 (dd, *J* = 10.5, 2.1 Hz, 1H), 6.38-6.36 (d, *J* = 10.5 Hz, 1H), 6.32-6.31 (d, *J* = 2.1 Hz, 1H), 3.72 (s, 3H), 1.53 (s, 6H). ^13^C NMR (176 MHz, CDCl_3_): δ = 179.98, 179.95, 174.49, 156.06, 140.54, 129.70, 125.16, 53.12, 47.44, 24.05, 23.95. HRMS (ESI): Calcd C_11_H_12_O_4_ [M+Na]^+^: 231.0633, Found: 231.0592.

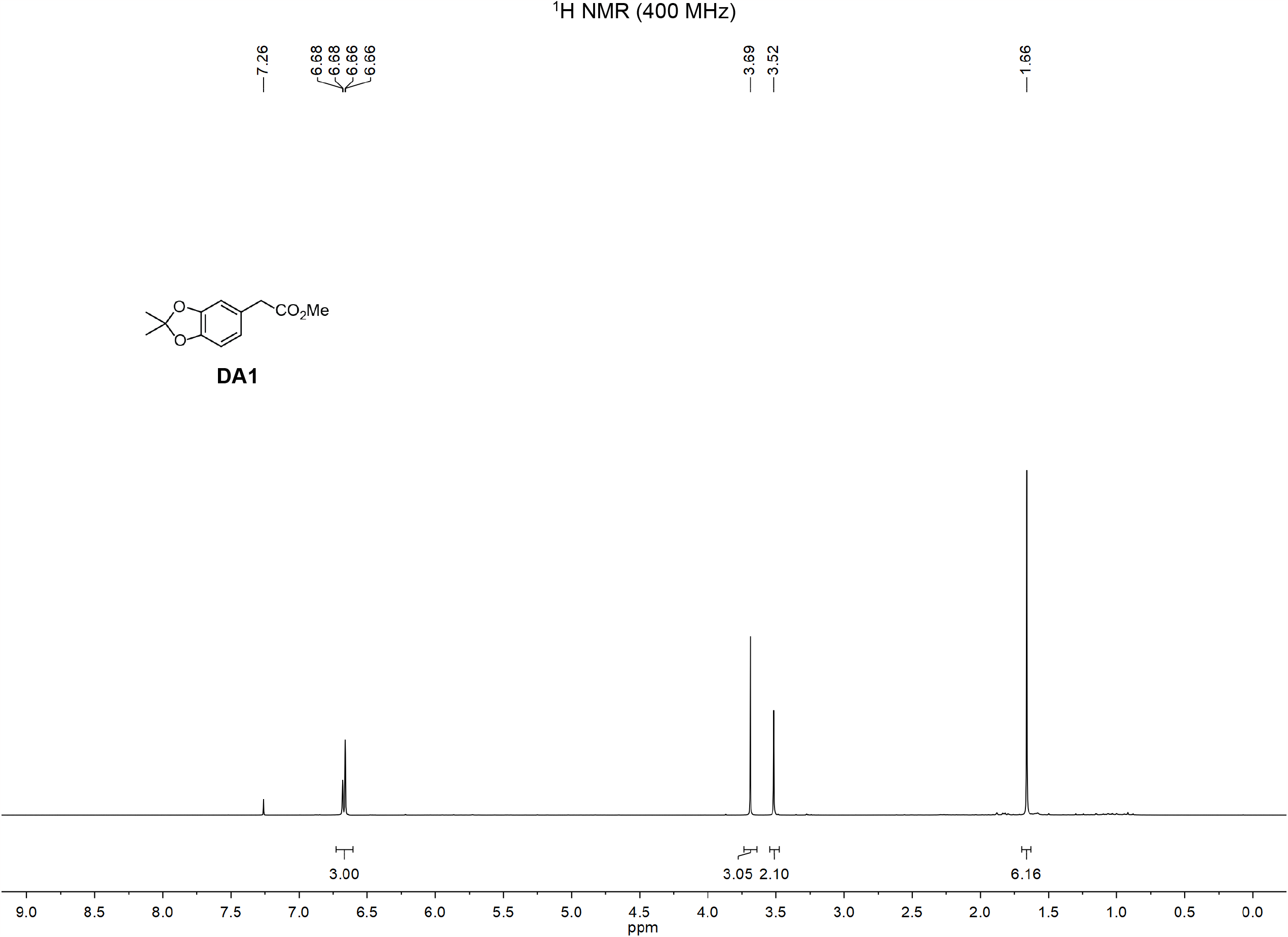

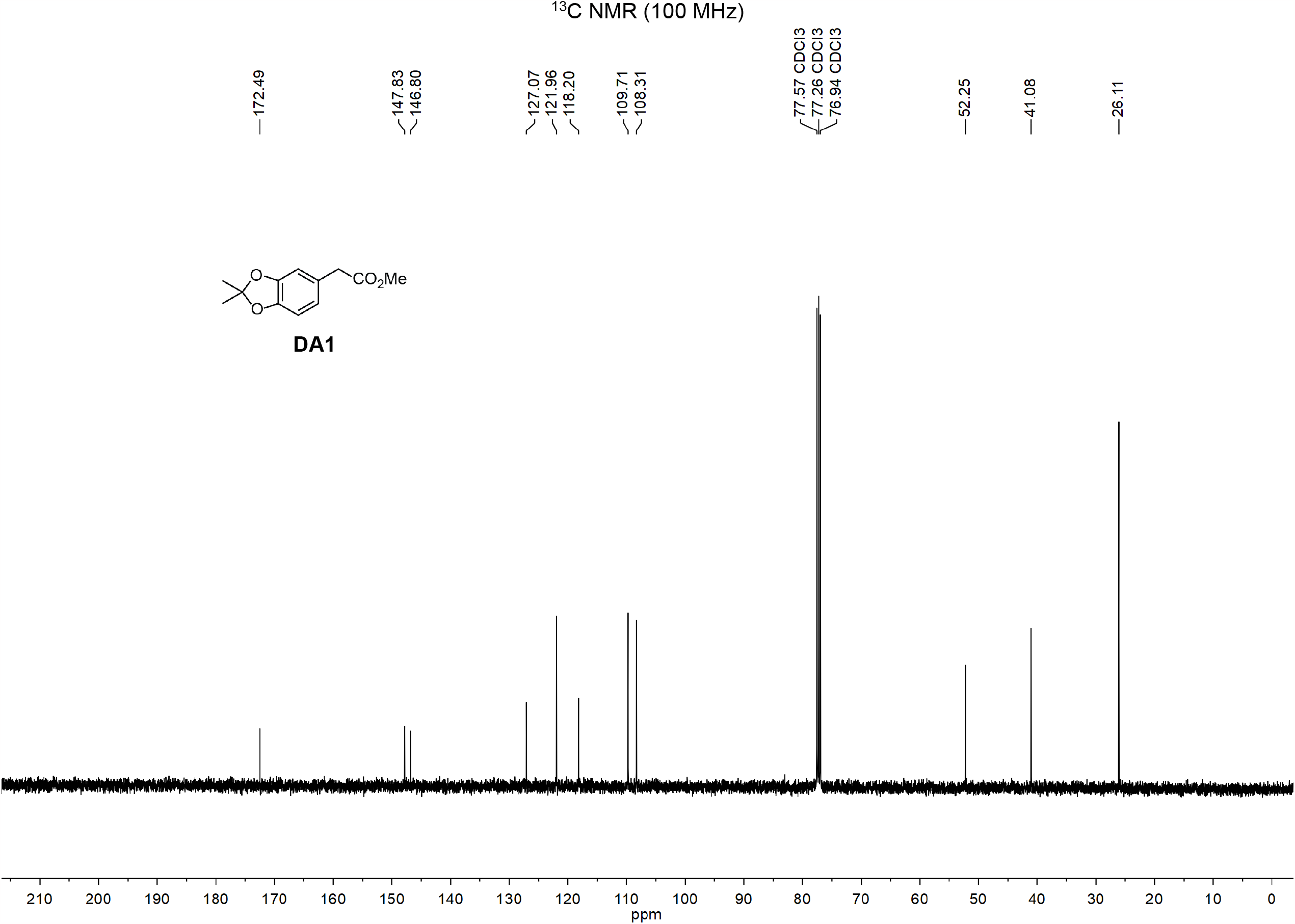

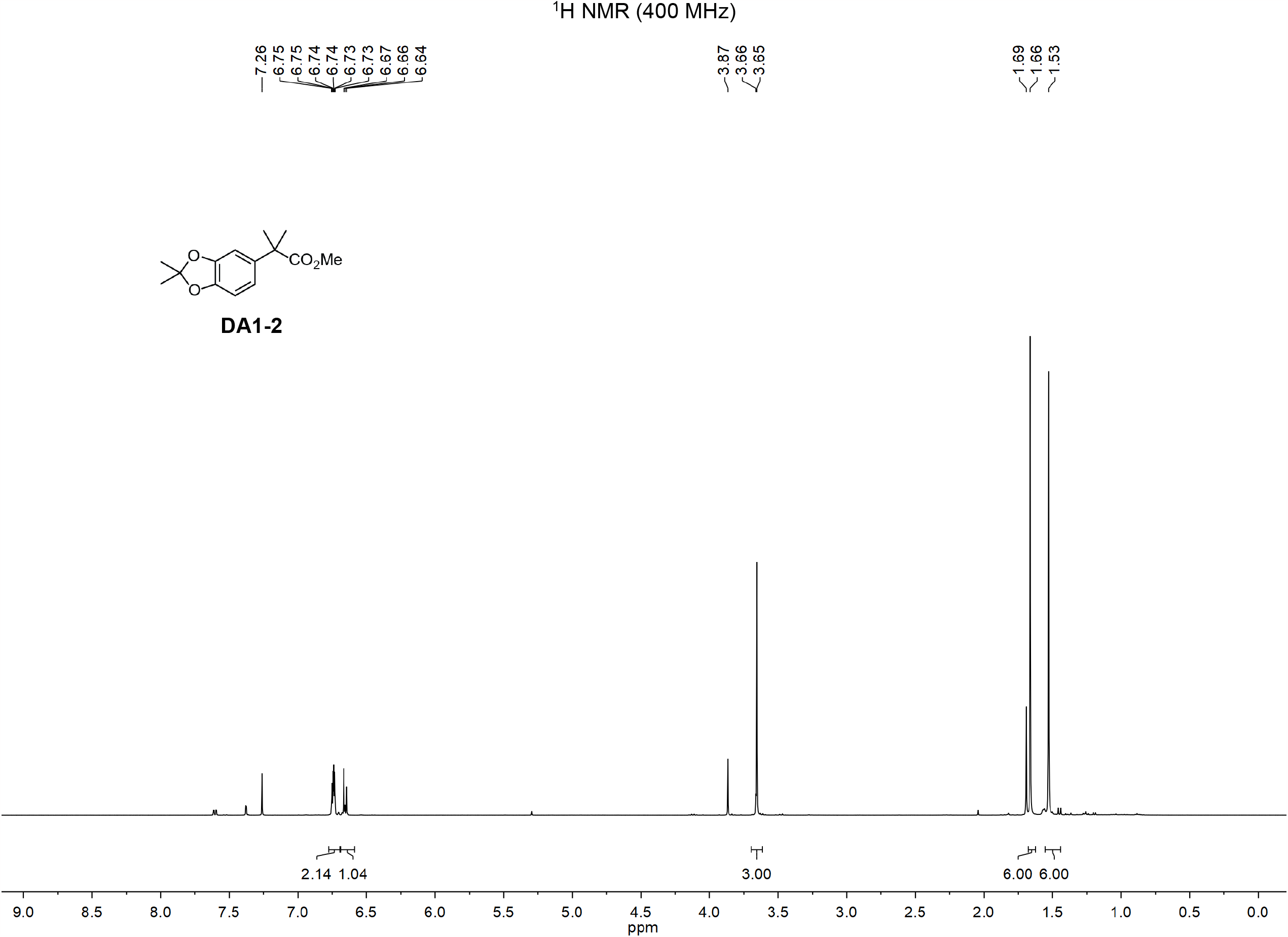

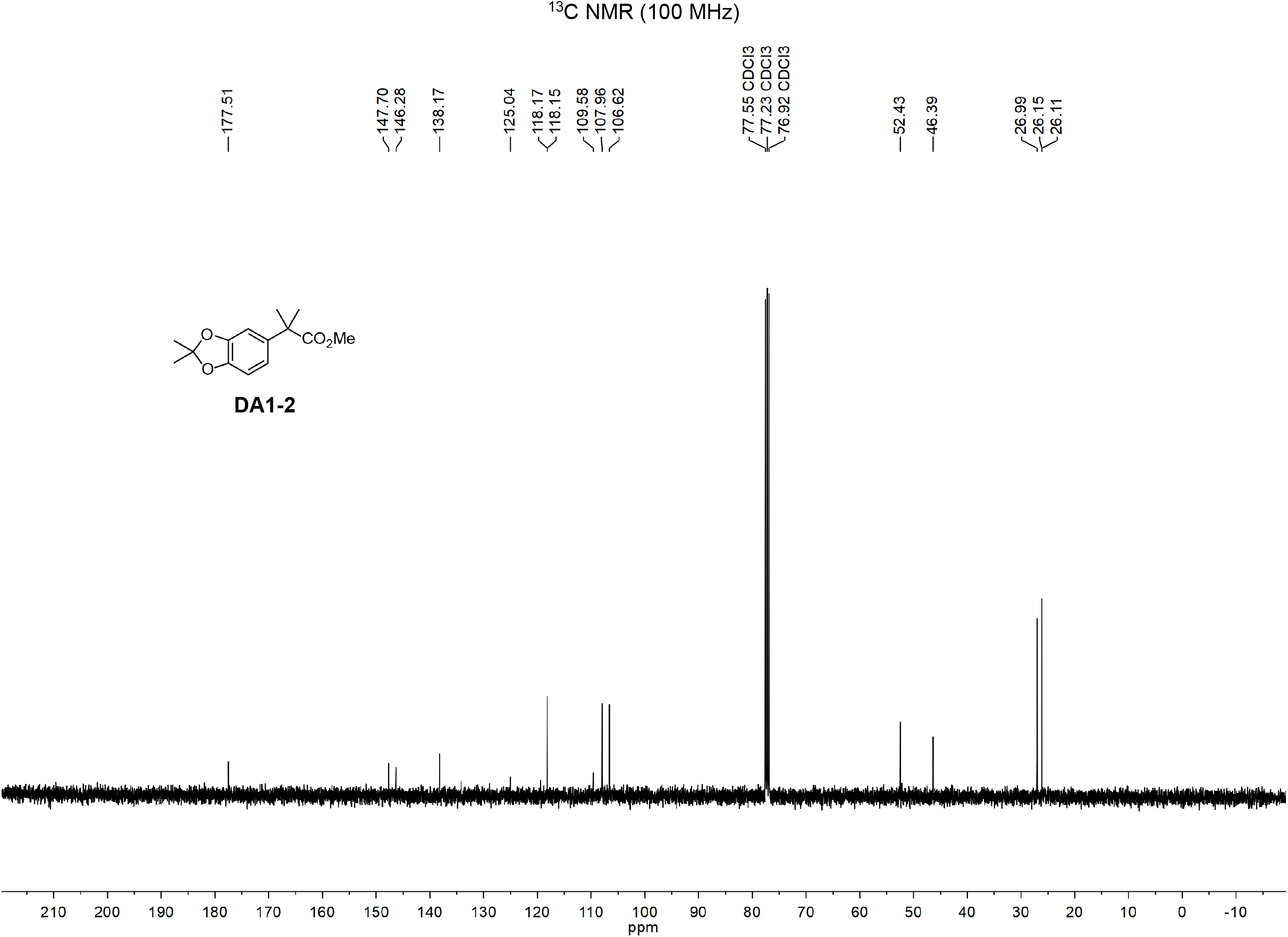

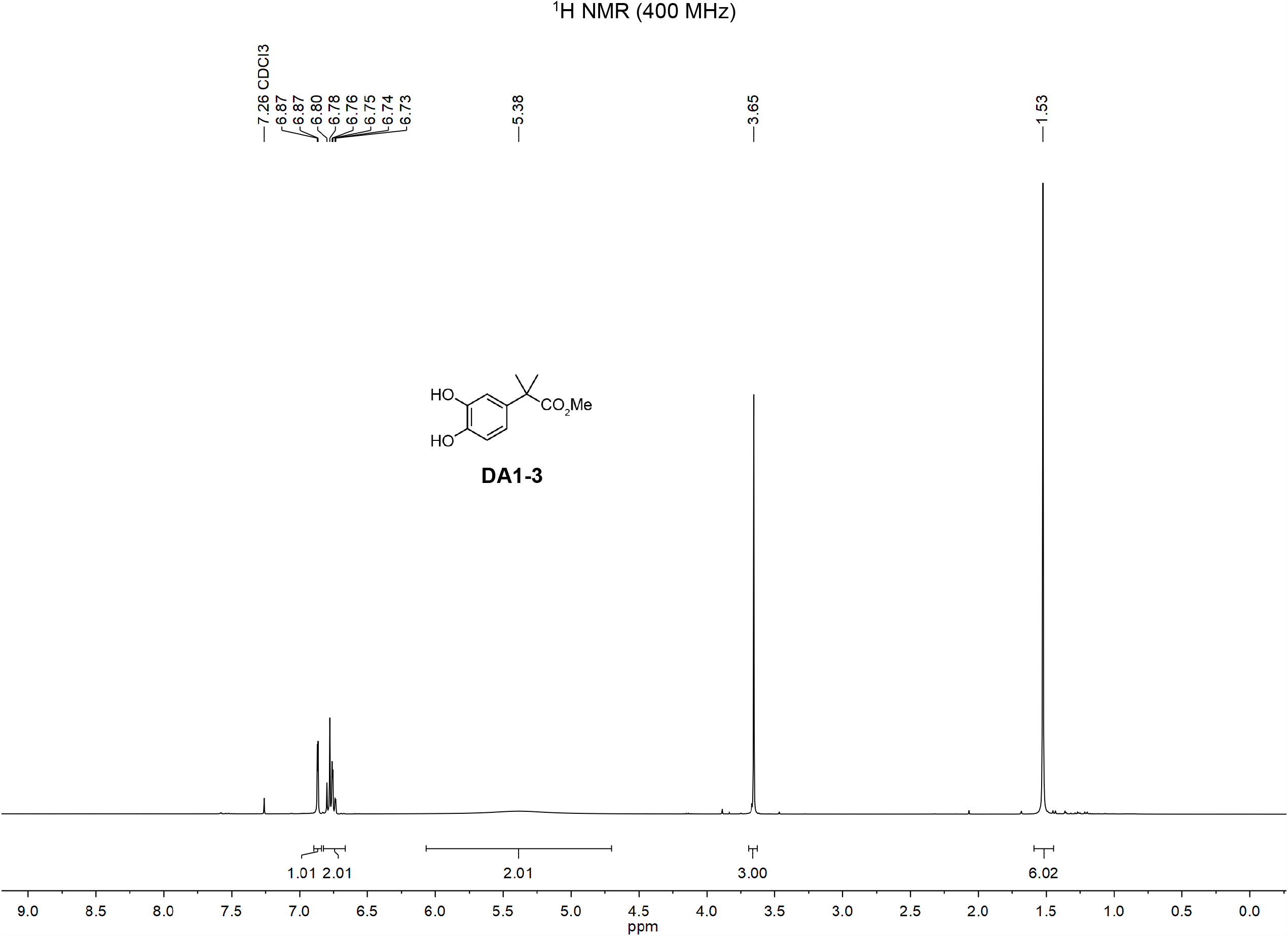

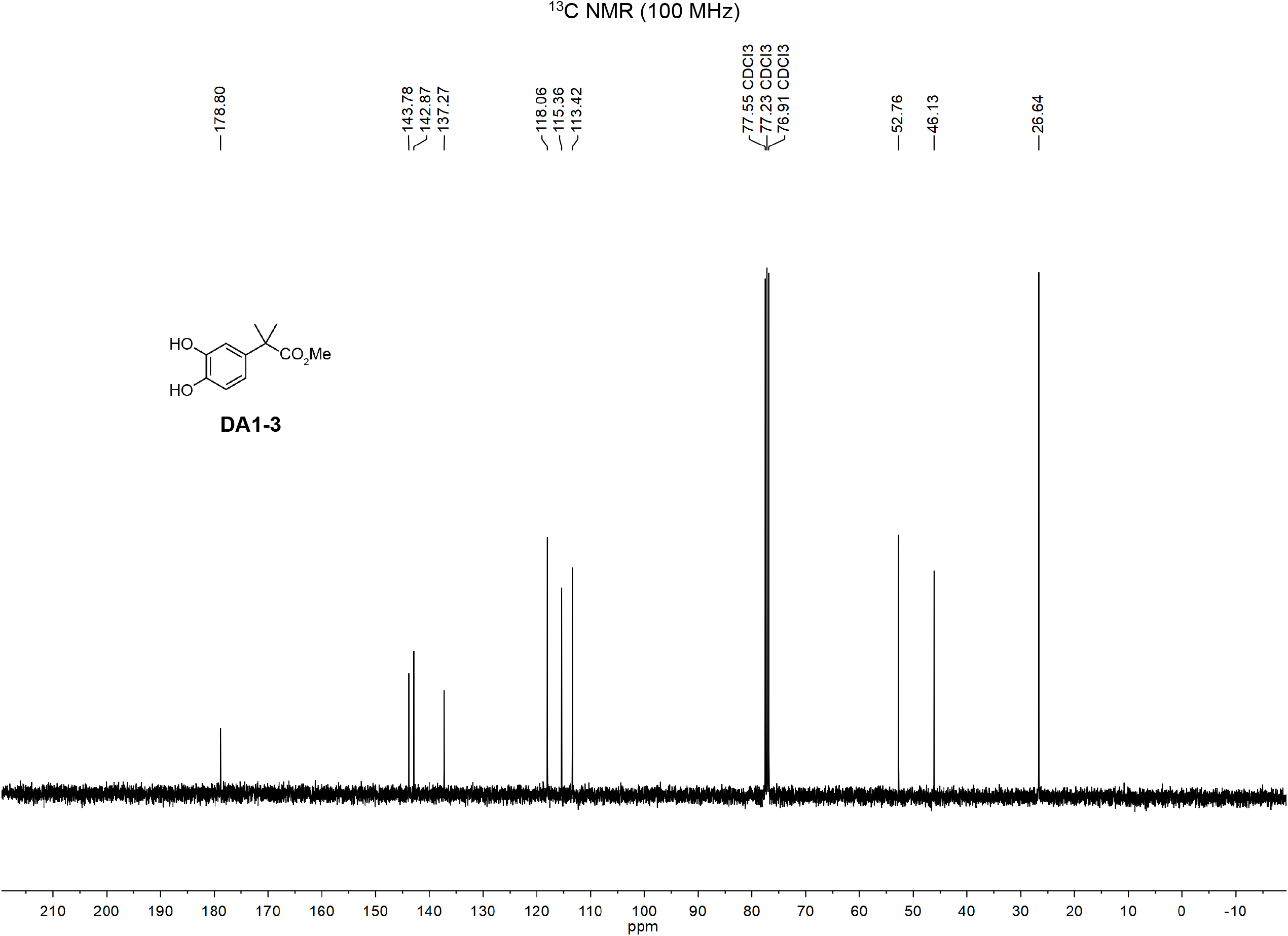

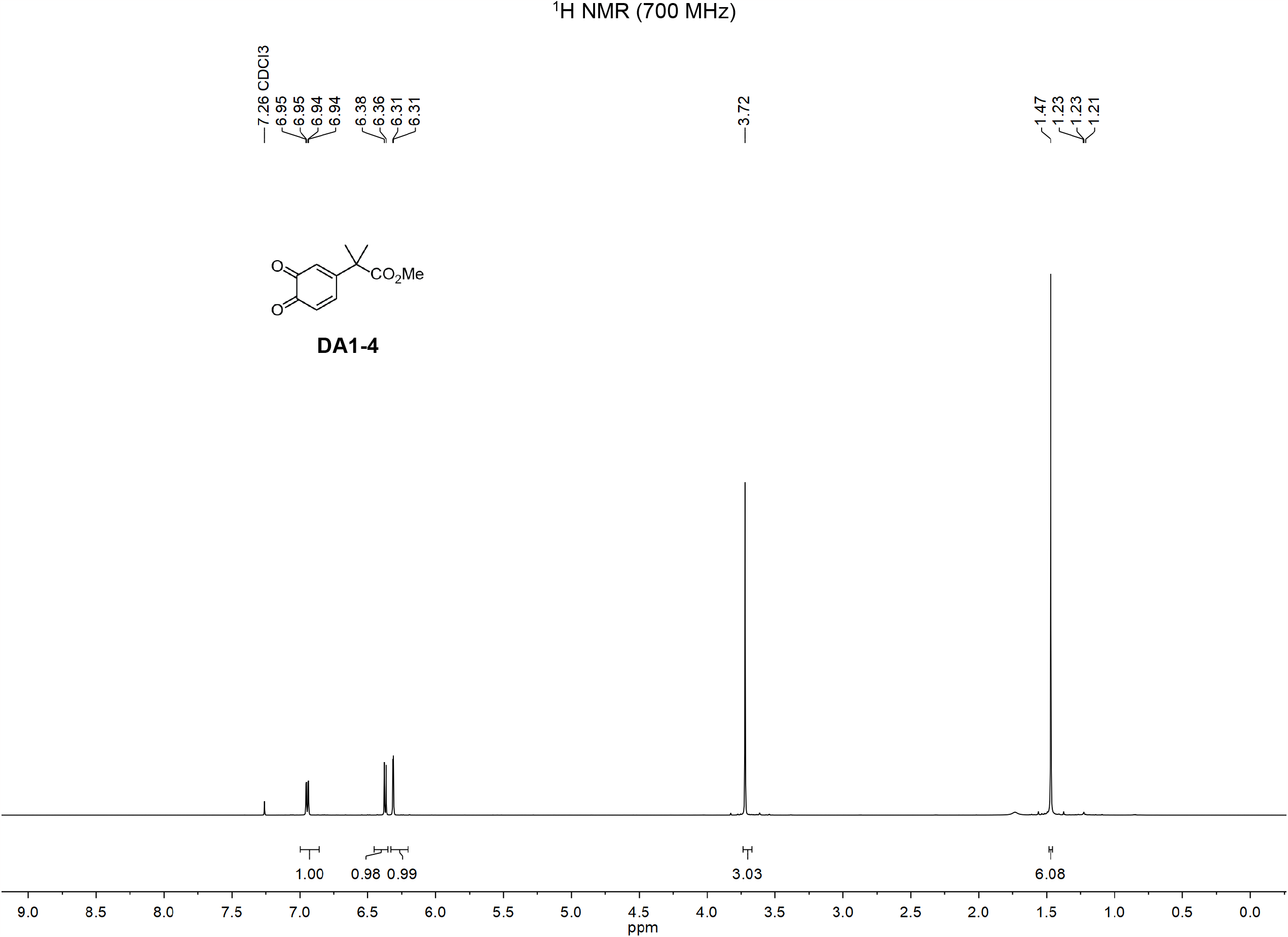

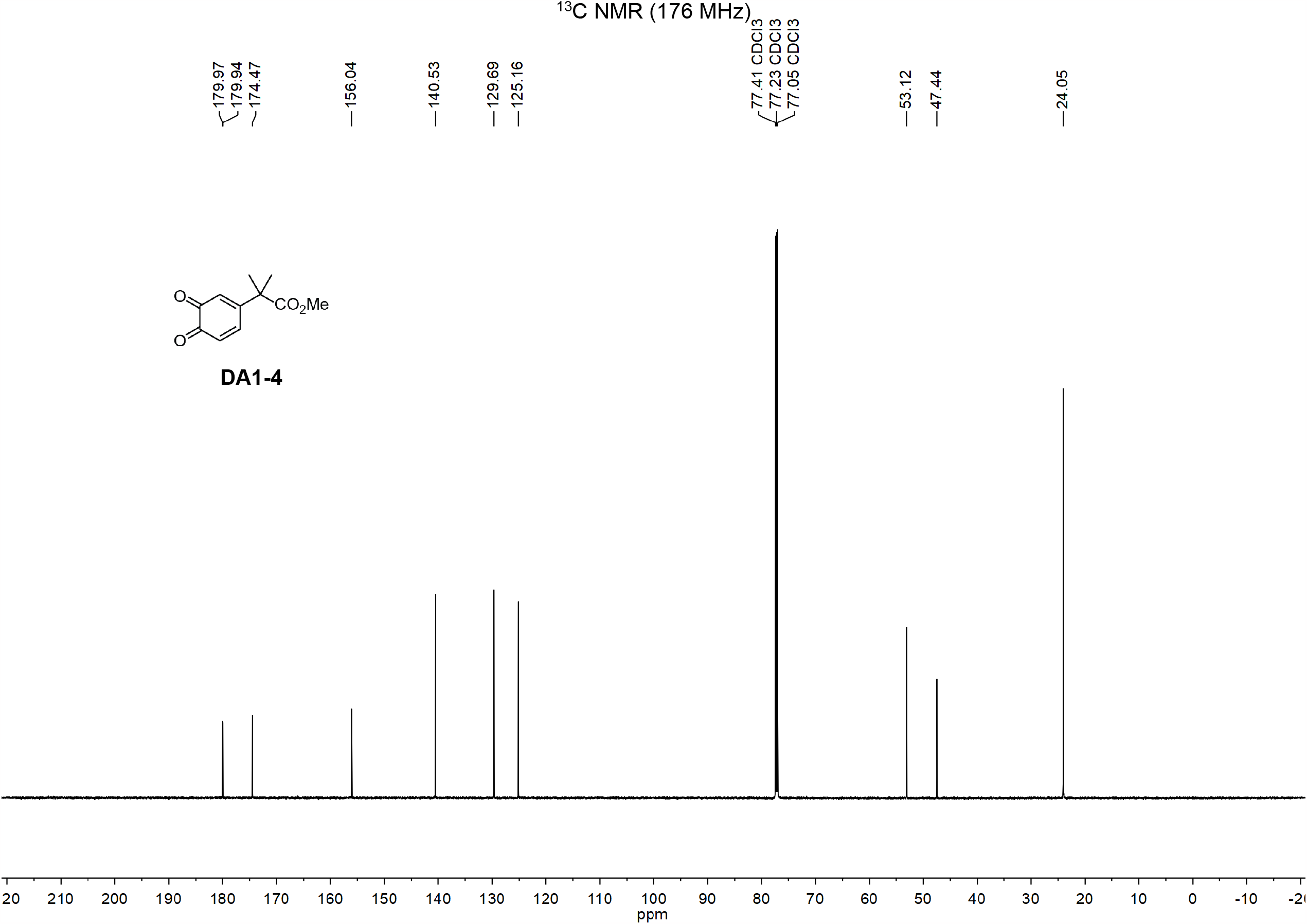

**Scheme 3:**
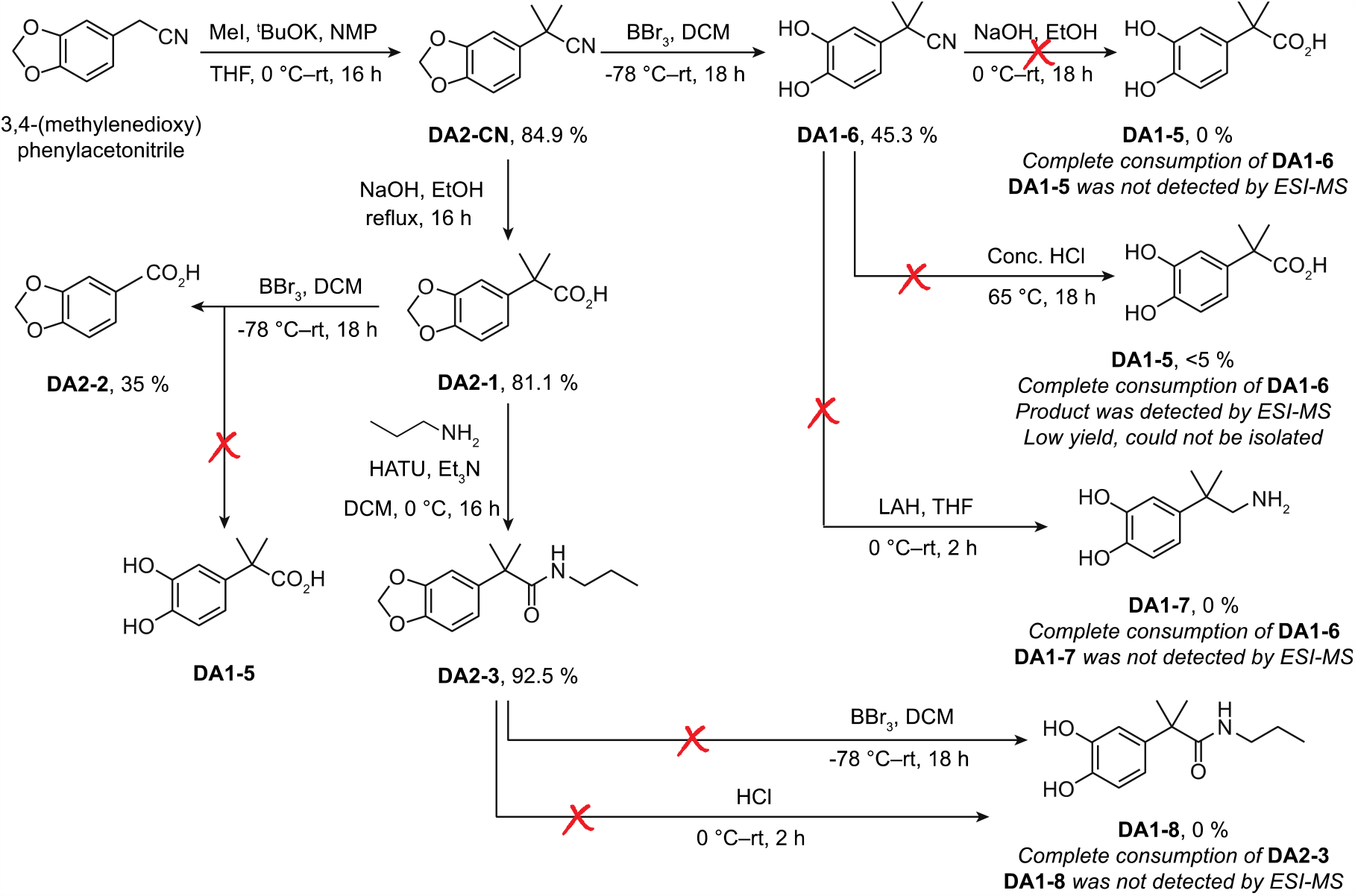
Methods for synthesis of compounds DA2-CN, DA1-6, DA2-1, DA2-3 and DA2-2.

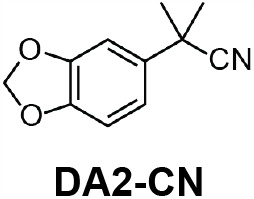

To an ice-cold suspension of 3,4-(methylenedioxy)phenylacetonitrile (0.5 g, 3.101 mmol, 1 eq) in THF/NMP (20 mL, v:v=1:1) was added ^t^BuOK (1.392 g, 12.404 mmol, 4 eq). The mixture was allowed to stir at 0 ^o^C for 10 min, and iodomethane (0.772 ml, 12.404 mmol, 4 eq) was slowly added into the solution for an additional 2 hour. The reaction mixture was extracted using DCM and water. The combined organic layer was further washed by water and brine, dried over anhydrous Na_2_SO_4_, filtered through filter paper and concentrated *in vacuo*. The crude product was purified by flash column chromatography on silica gel using 10% EtOAc/hexanes (v/v) to obtain compound **DA2-CN** (0.586 g, 84.9%). R_f_ = 0.67 (20% EtOAc/Hexanes, visualized w/ UV). ^1^H NMR (700 MHz, CDCl_3_) δ = 6.95-6.92 (m, 2H), 6.79-6.78 (d, *J* = 7.7 Hz, 1H), 5.97 (s, 2H), 1.68 (s, 6H). ^13^C NMR (176 MHz, CDCl_3_): δ = 148.33, 147.29, 135.52, 124.78, 119.59, 108.49, 106.08, 101.59, 37.03, 29.48. HRMS (ESI): Calcd C_11_H_11_NO_2_ [M+H]^+^: 190.0868, Found: 190.0868

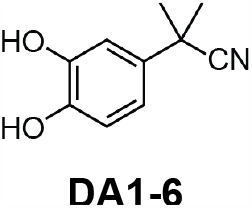

To a solution of compound **DA2-CN** (0.251 g, 1.327 mmol, 1eq) in DCM (10 mL) under N_2_ at -78 ^o^C was slowly added liquid BBr_3_ (0.996 g, 3.981 mmol, 3 eq). The mixture was allowed to warm up gradually to room temperature and stirred overnight. The reaction was quenched gently with water. The mixture was extracted with DCM and water, followed by brine washing. The combined organic layer was dried over anhydrous Na_2_SO_4_, concentrated *in vacuo*, and purified by flash chromatography over silica gel to give compound **DA1-6** (0.106 g, 45.3%). R_f_ = 0.26 (30% EtOAc/Hexanes, visualized w/ UV and KMnO_4_). ^1^H NMR (400 MHz, CDCl_3_) δ = 7.03 (s, 1H), 6.87-6.86 (d, *J* = 0.8 Hz, 1H), 5.96-5.85 (br, 1H), 5.42-5.29 (br, 1H), 1.69 (s, 6H). ^13^C NMR (101 MHz, CDCl_3_): δ = 143.98, 143.51, 134.35, 124.81, 117.45, 115.69, 112.91, 36.83, 29.42. LRMS (ESI): Calcd C_10_H_11_NO_2_ [M+H]^+^: 178.1, Found: 178.1.

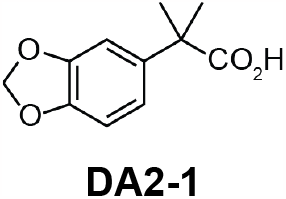

To compound **DA2-CN** (0.61 g, 3.203 mmol, 1 eq) in EtOH (10 mL) was added aqueous NaOH (1.79 g, 44.8 mmol in 10 mL water). The mixture was allowed to stir at 120 ^o^C for 16 hours. The methanol was removed *in vacuo*. The solution was acidified by 2N aqueous HCl to pH = 1. The reaction mixture was extracted using DCM and water. The combined organic layer was further washed by water and brine, dried over anhydrous Na_2_SO_4_, filtered through filter paper, and concentrated *in vacuo*. The crude product was purified by flash column chromatography on silica gel using 40% EtOAc/hexanes (v/v) to obtain compound **DA2-1** (0.540 g, 81.08%). R_f_ = 0.38 (40% EtOAc/Hexanes, visualized w/ UV). ^1^H NMR (700 MHz, CDCl_3_) δ = 6.91 (s, 1H), 6.87-6.85 (d, *J* = 8.4 Hz, 1H), 6.78-6.76 (d, *J* = 8.4 Hz, 1H), 5.95 (s, 2H), 1.56 (s, 6H). ^13^C NMR (176 MHz, CDCl_3_): δ = 182.47, 147.95, 146.66, 137.96, 119.14, 108.25, 107.07, 101.32, 46.13, 26.61. HRMS (ESI): Calcd C_11_H_12_O_4_ [M+H]^+^: 209.0814, Found: 209.0886.

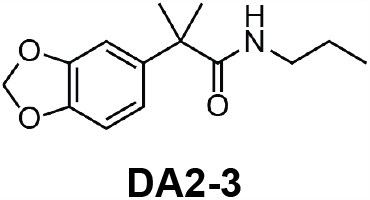

To an ice-cold suspension of compound **DA2-1** (24 mg, 0.12 mmol, 1 eq) in DCM (5 mL) was added HATU (48 mg, 0.13 mmol, 1.1 eq) and triethylamine (0.041 ml, 0.29 mmol, 2.5 eq). The mixture was stirred at 0 ^o^C for 5 min, and propylamine (0.011 ml, 0.13 mmol, 1.1 eq) was slowly added into the solution at 0 ^o^C. The reaction was stirred for an additional 16 hours. The reaction mixture was extracted using DCM and water. The combined organic layer was further washed by water and brine, dried over anhydrous Na_2_SO_4_, filtered through filter paper, and concentrated *in vacuo*. The crude product was purified by flash column chromatography on silica gel using 30% EtOAc/hexanes (v/v) to obtain compound **DA2-3** (27 mg, 92.5%). R_f_ = 0.75 (60% EtOAc/Hexanes, visualized w/ UV and KMnO_4_). ^1^H NMR (400 MHz, CDCl_3_) δ = 6.83-6.76 (m, 3H), 5.95 (s, 2H), 5.22 (br, 1H), 3.14-3.08 (q, *J* = 12.8, 7.2, 2H),1.51-1.30 (m, 2H), 0.82-0.78 (t, *J* = 7.2, 3H). ^13^C NMR (176 MHz, CDCl_3_): δ = 177.59, 148.13, 146.64, 139.49, 119.60, 108.35, 107.56, 101.33, 47.01, 41.59, 27.54, 22.93, 11.43. HRMS (ESI): Calcd C_14_H_19_NO_3_ [M+H]^+^: 250.1443, Found: 250.1452.

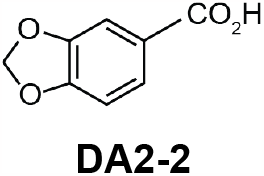

To compound **DA2-1** (0.05 g, 0.24 mmol, 1 eq) in DCM (5 mL) under N_2_ and -78 ^o^C was added BBr_3_ (0.041 mL, 0.24 mmol, 1 eq). The mixture was allowed to stir at rt gradually for overnight. The reaction was quenched gently by water. The mixture was extracted with DCM and water, followed by brine washing. The combined organic layer was dried over anhydrous Na_2_SO_4_, concentrated *in vacuo*, and purified by flash chromatography over silica gel to give compound **DA2-2** (0.106 g, 45.3%). R_f_ = 0.75 (60% EtOAc/Hexanes, visualized w/ UV and KMnO4). ^1^H NMR (400 MHz, DMSO-d_6_): = 12.74 (s, 2H), 7.54 (dd, *J* = 8.1, 1.7 Hz, 1H), 7.36 (d, *J* = 1.7 Hz, 1H), 7.00 (d, *J* = 8.2 Hz, 1H), 6.12 (s, 2H). ^13^C NMR (101 MHz, CD_3_OD): δ = 169.33, 153.20, 149.21, 126.53, 125.75, 110.29, 108.85, 103.33. HRMS (ESI): Calcd C_8_H_6_O_4_ [M+H]^+^: 166.0344, Found: 167.0350.

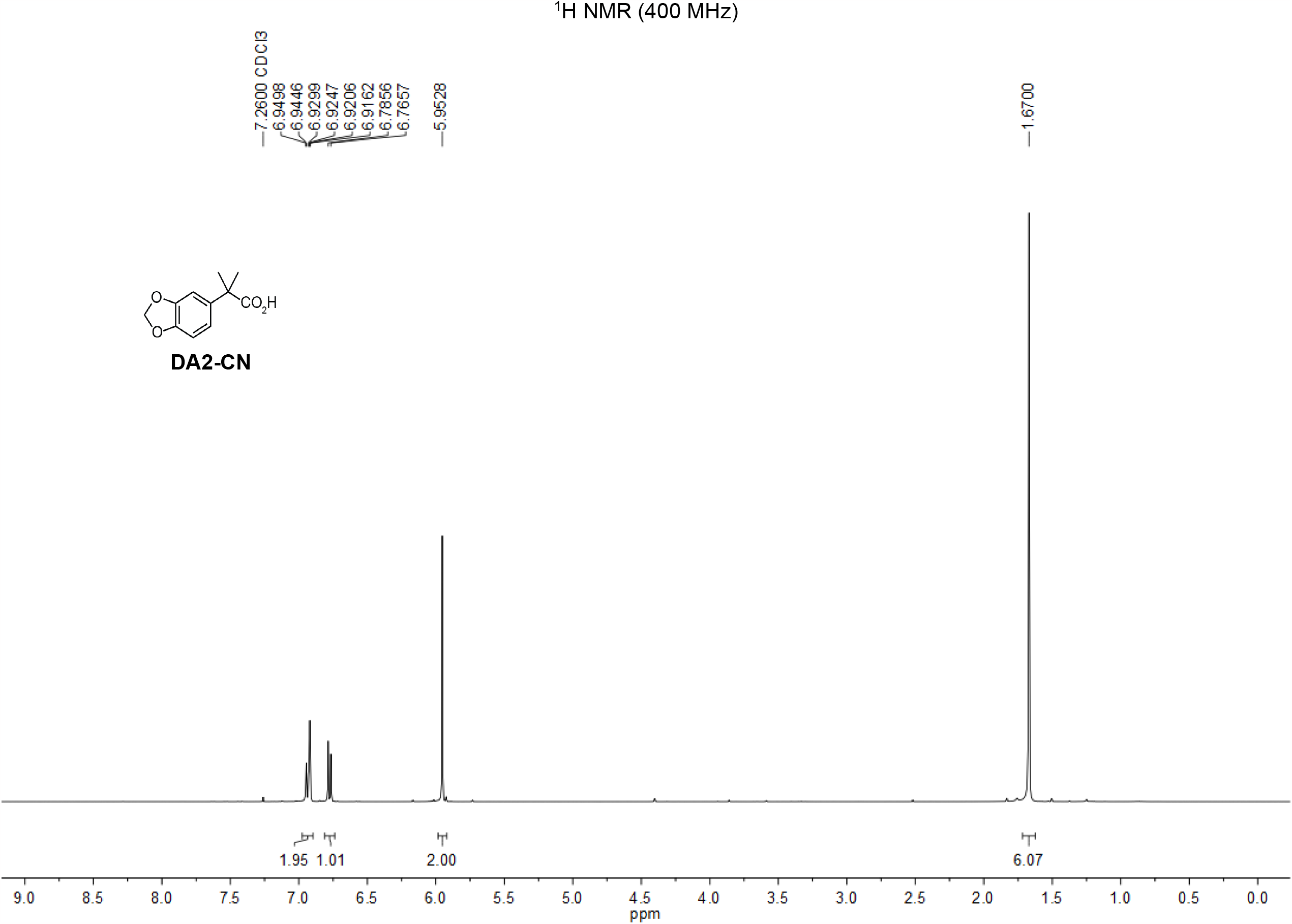

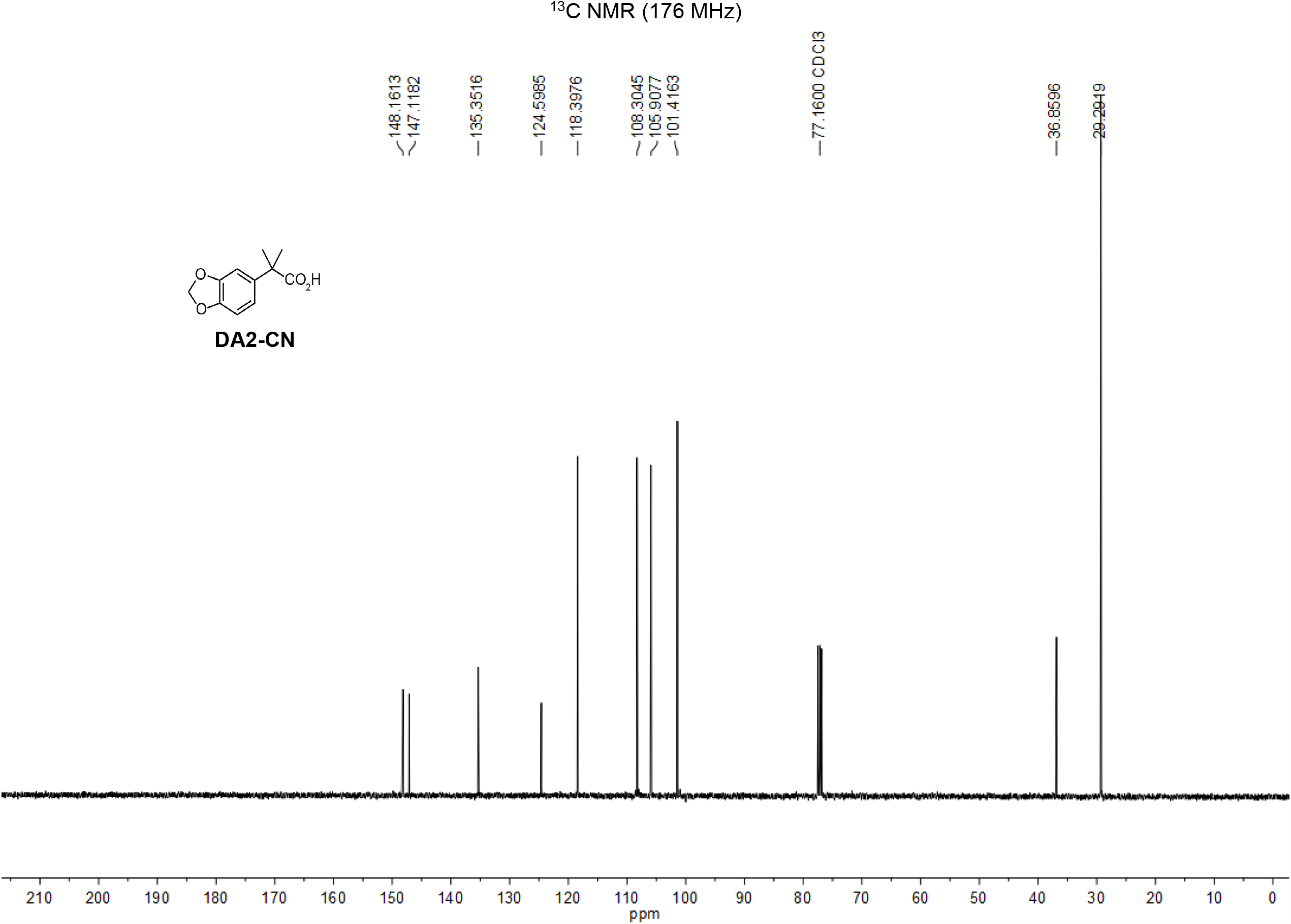

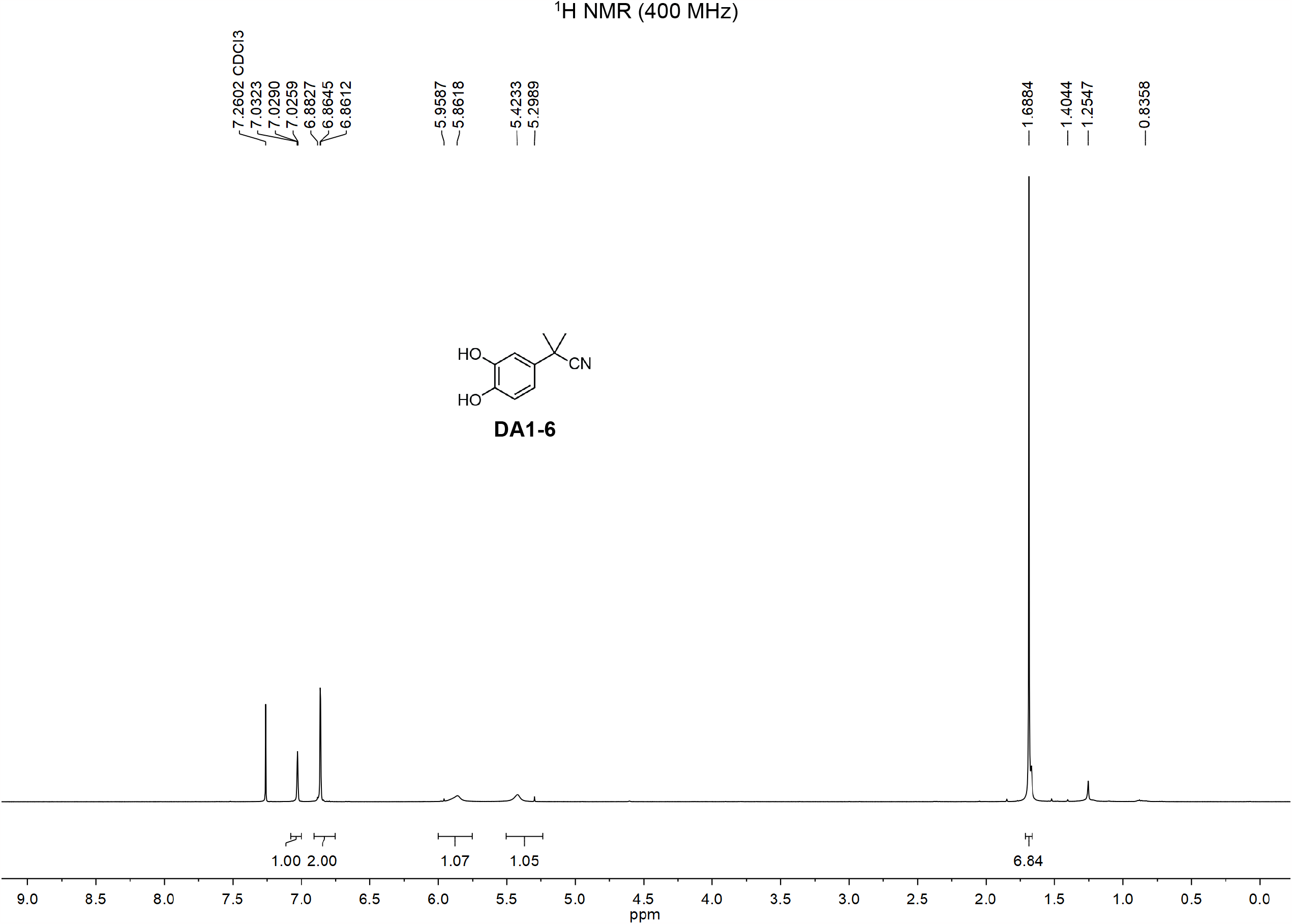

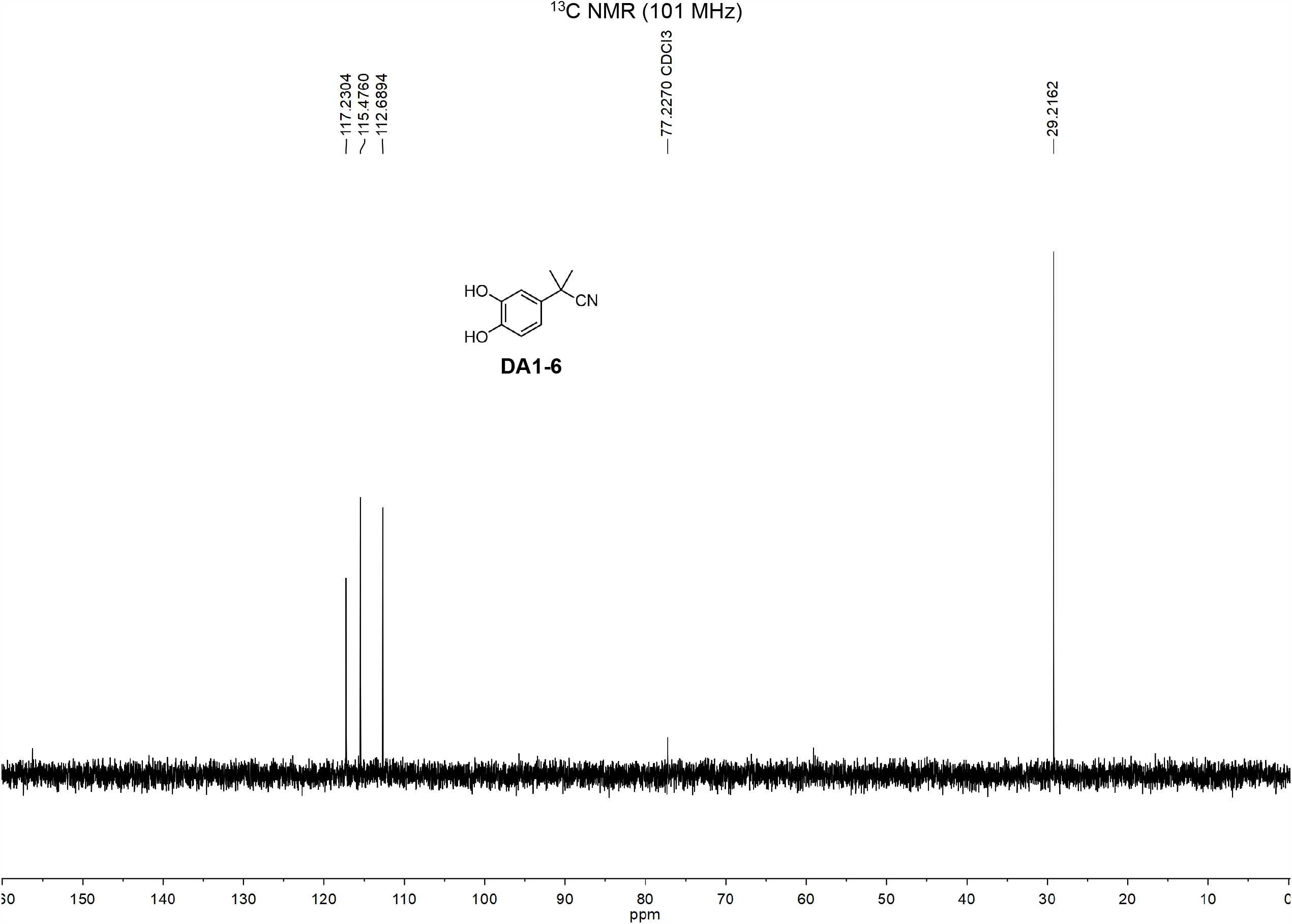

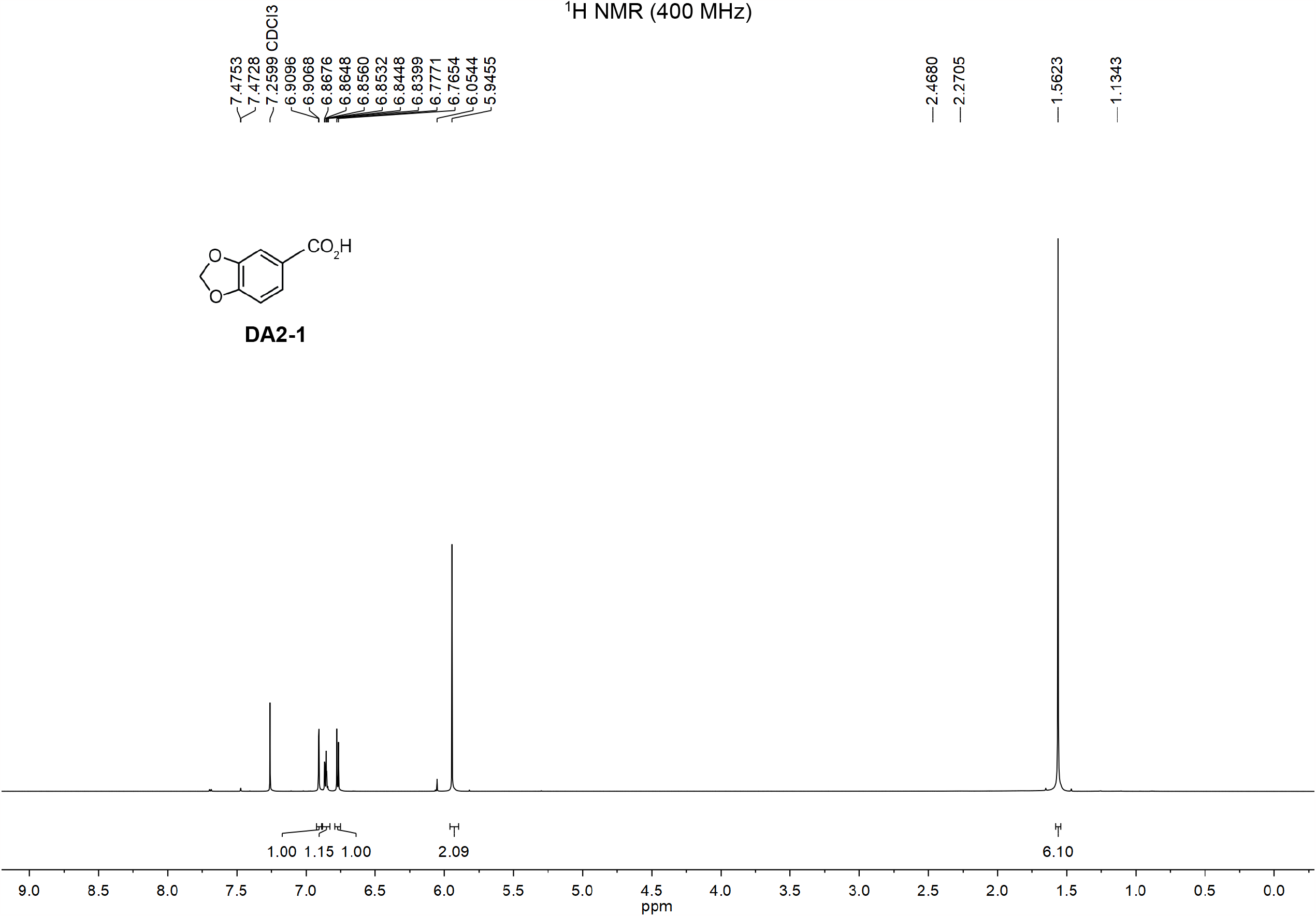

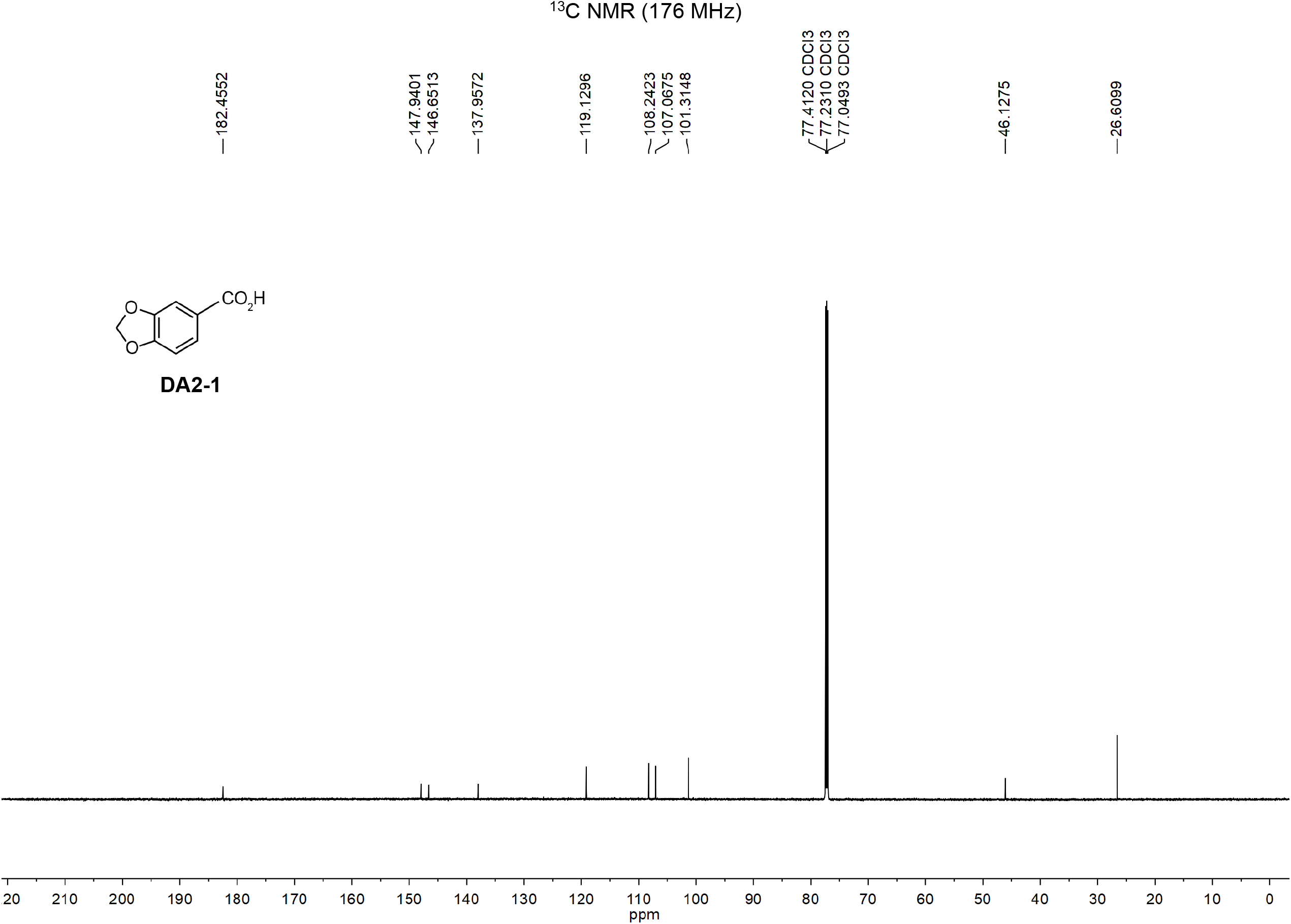

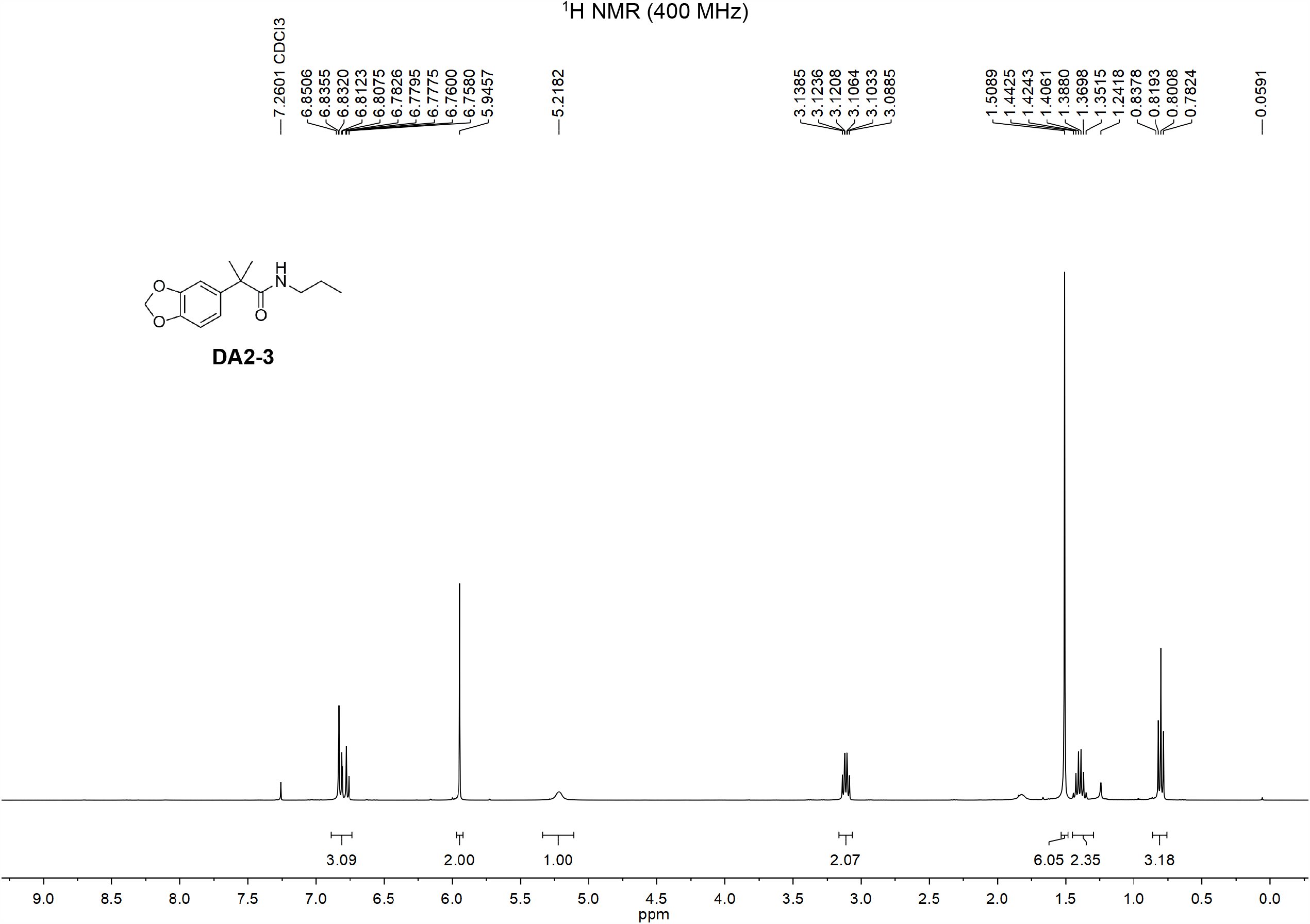

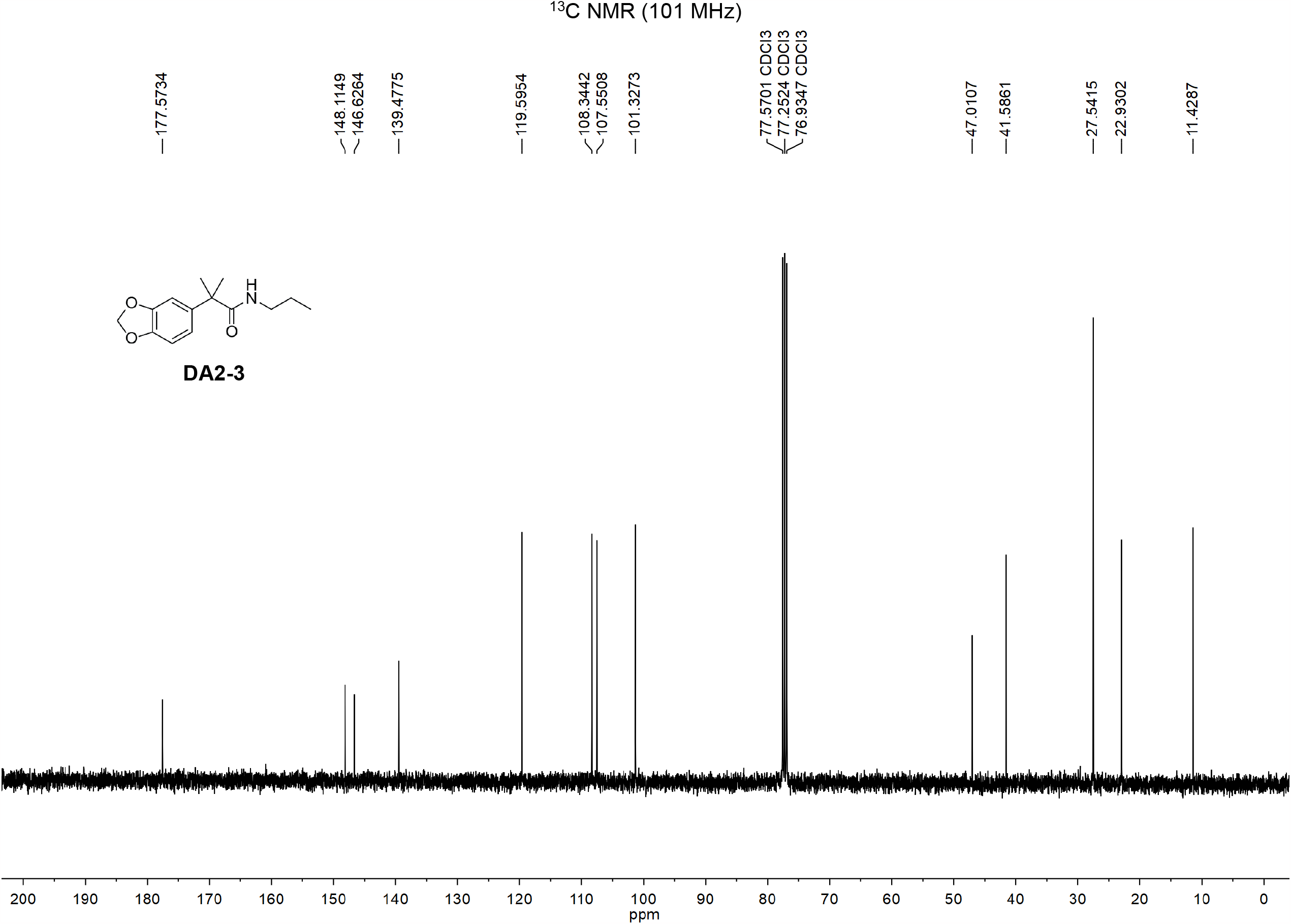

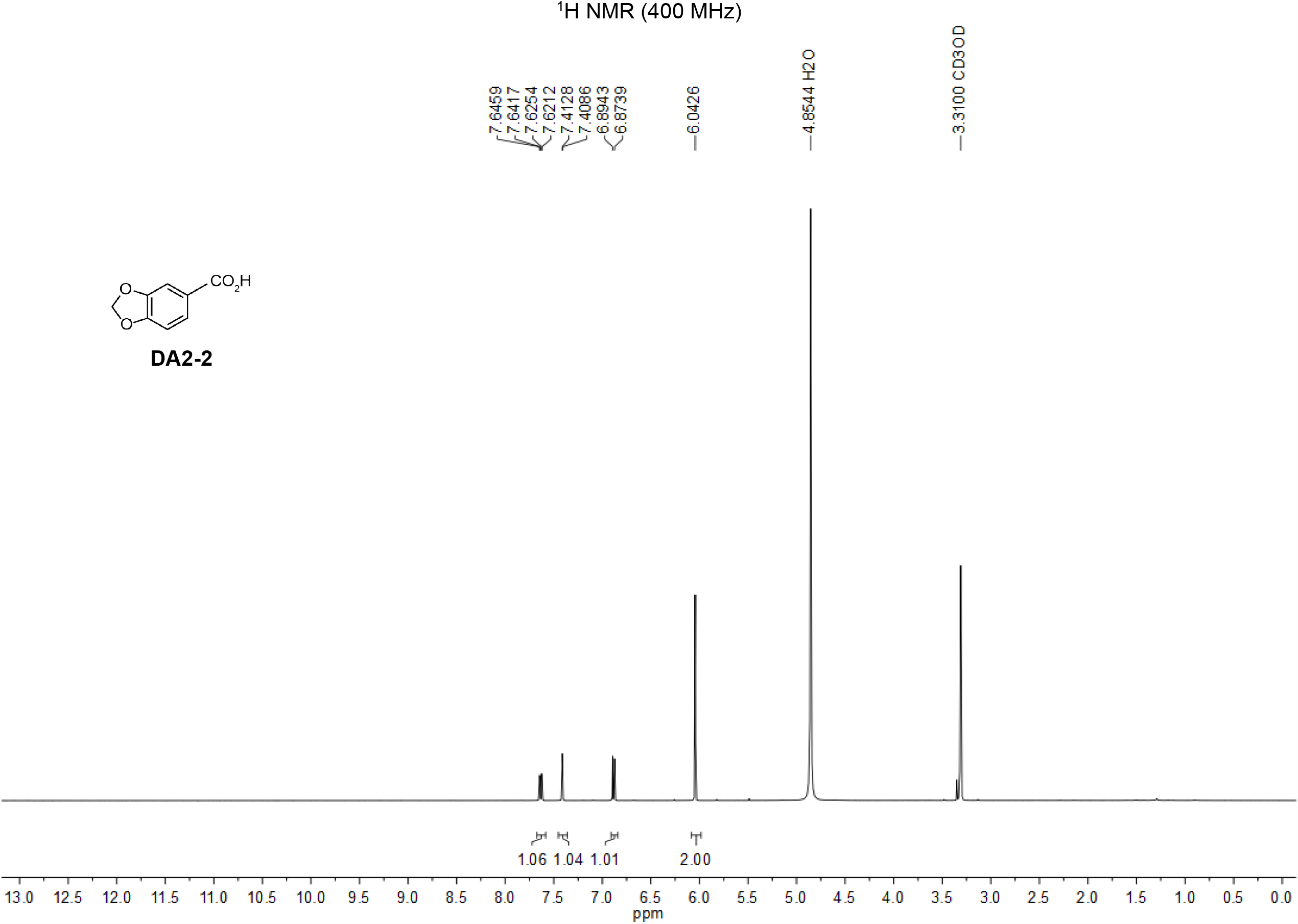

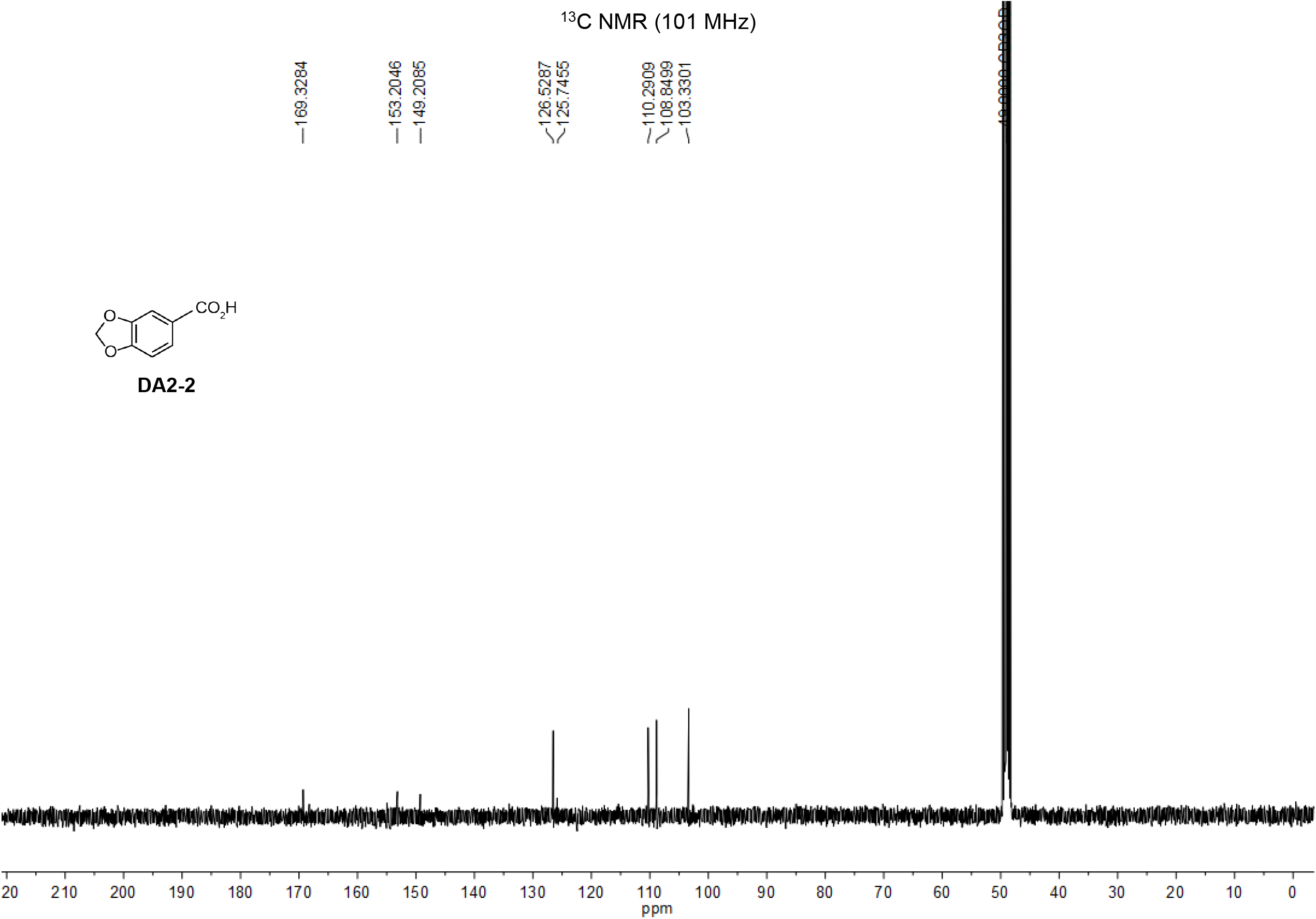

**Scheme 4:**
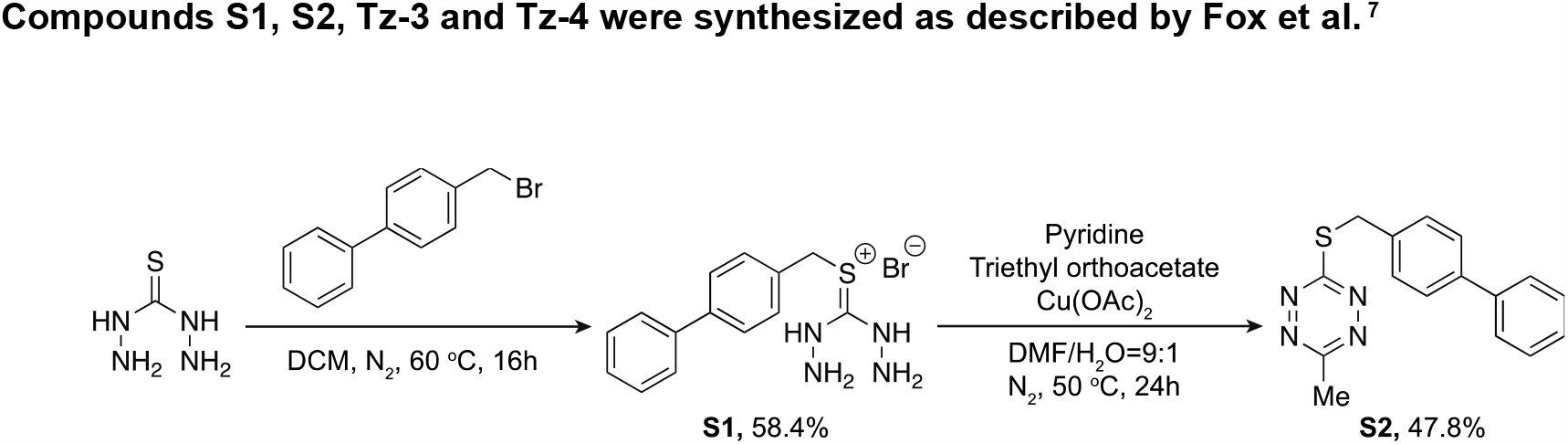
Methods for synthesis of compounds S1, S2, Tz-3 and Tz-4.

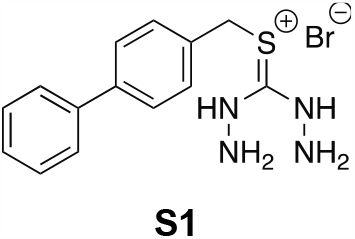

To a thiocarbohydrazine (1.73 g, 16.3 mmol, 1.0 eq) in ethanol (40 mL, 0.40 M) was added 4-bromomethyl biphenyl (4.04 g, 16.3 mmol, 1.0 eq). The mixture was stirred at 60 ^o^C overnight. A white cloud-like solid formed during the reaction. The mixture was allowed to cool down to room temperature, and diethyl ether (50 mL) was added to the mixture. The white solids were filtered and collected by vacuum filtration, washed with diethyl ether three times, and dried under the Schlenk line to obtain compound **S1** (3.37 g, 58.4%). ^1^H NMR (400 MHz, CD_3_OD) δ = 7.59-7.53 (m, 4H), 7.43 (d, *J* = 8 Hz, 2H), 7.36 (t, *J* = 7.2 Hz, 2H), 7.26 (t, *J* = 7.2 Hz, 1H), 4.24 (s, 2H). ^13^C NMR (100 MHz, CDCl_3_): δ = 142.78, 141.73, 130.97, 130.86, 130.09, 128.83, 128.76, 128.58, 128.10, 128.05. LRMS (ESI+): Calcd C_14_H_17_BrN_4_S [M-Br]^+^: 273.1, Found: 273.1.

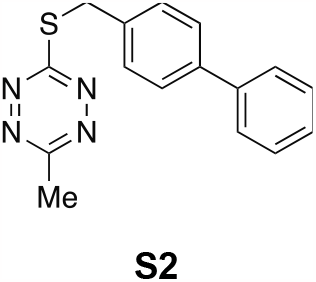

Compound **S1** (0.915 g, 2.59 mmol, 1.0 eq), pyridine (0.418 mL, 5.18 mmol, 2.0 eq), and DMF (10 mL, 0.25 M) were mixed in RBF under an inert atmosphere at 50 ^o^C. Triethyl orthoacetate (0.949 mL, 5.18 mmol, 2.0 eq) was slowly added to the reaction using the syringe pump at 0.5 mL/hr. The mixture was stirred at 60 ^o^C overnight, and DMF/H_2_O (5 mL, DMF: H_2_O=9:1) was added after cooling to room temperature. Cu(II)(OAc)_2_ was added to the solutions and let the reaction stir for another 24 hours with air purging. The reddish-purple solid will be crushed after adding H_2_O (50 mL). Collect the red solid through vacuum filtration and washed with cold water three times. The crude product was purified by flash chromatography using 60% DCM/hexanes (v/v) to obtain compound **S2**. R_f_ = 0.52 (60% DCM/Hexanes, visualized w/ UV). ^1^H NMR (400 MHz, CDCl_3_) δ = 7.58-7.52 (m, 6H), 7.41 (d, *J* = 7.2 Hz, 2H), 7.33 (t, *J* = 7.2 Hz, 1H), 4.58 (s, 2H), 2.98 (s, 3H). ^13^C NMR (100 MHz, CDCl_3_): δ = 174.83, 165.56, 141.07, 140.75, 134.95, 129.93, 129.04, 127.67, 127.29, 34.65, 20.93. LRMS (ESI+): Calcd C_16_H_15_N_4_S+ [M+H]^+^: 295.1, Found: 295.1.

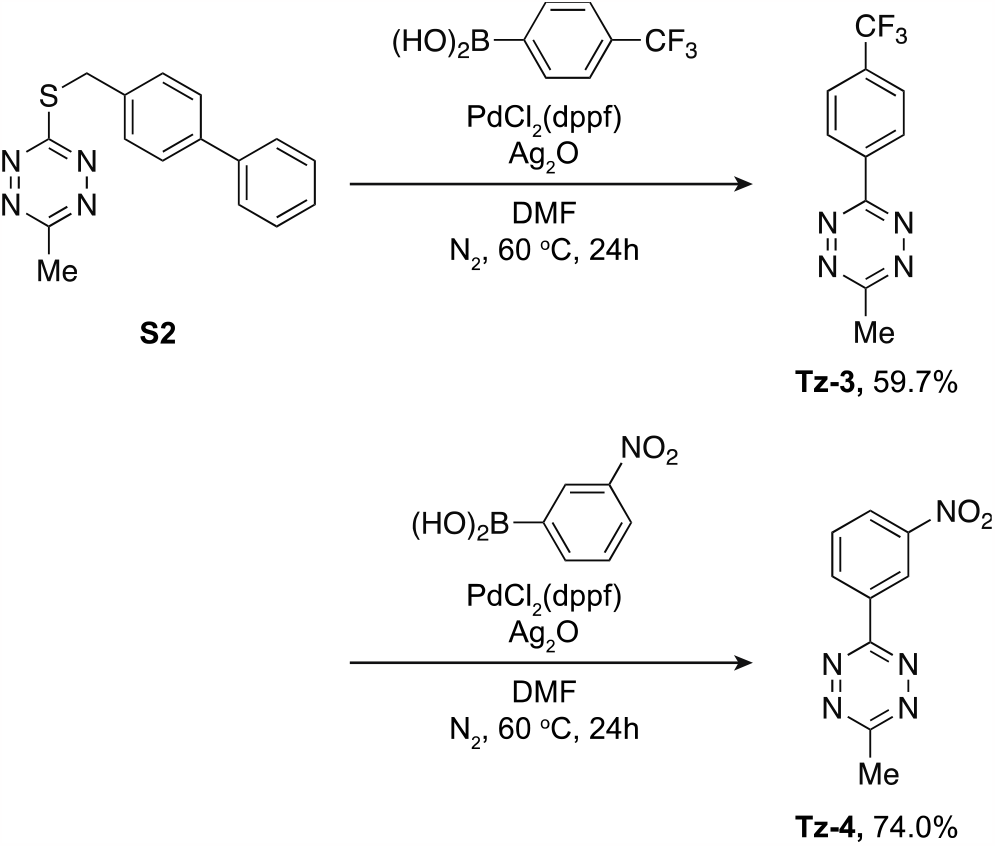

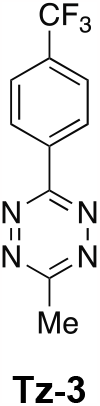

To compound **S2** (0.373 g, 1.27 mmol, 1.0 eq) in DMF (12 mL, 0.1M) under N_2_ was added 4- (Trifluoromethyl)phenylboronic acid (0.458 g, 2.41 mmol, 1.9 eq), PdCl_2_(dppf) (139 mg, 0.19 mmol, 0.15 eq) and Ag_2_O (0.736 g, 3.18 mmol, 2.5 eq). The mixture was stirred at 60 ^o^C overnight. The mixture was allowed to back to room temperature, then DMF was removed *in vacuo*. The crude product was purified by flash chromatography over silica gel using 50% DCM/hexanes (v/v) to give compound **Tz-3** (0.182 g, 59.7%) as the pink solid. R_f_ = 0.70 (50% DCM/Hexanes, visualized w/ UV). ^1^H NMR (400 MHz, CDCl_3_): = 8.72 (d, *J* = 8 Hz, 2H), 7.85 (d, *J* = 8 Hz, 2H), 3.14 (s, 3H). ^13^C NMR (101 MHz, CDCl_3_): δ = 168.12, 163.51, 128.48, 126.47. 126.43, 53.66, 21.51. ^19^F NMR (380 MHz, CDCl_3_): δ = 63.11. LRMS (ESI+): Calcd C_10_H_8_F_3_N_4_+ [M+H]^+^: 241.1, Found: 241.1.

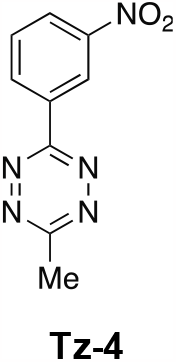

To compound **S2** (0.104 g, 0.35 mmol, 1.0 eq) in DMF (10 mL) under N_2_ was added 3-Nitrophenylboronic acid (0.112 g, 0.67 mmol, 1.9 eq), PdCl_2_(dppf) (38 mg, 0.053 mmol, 0.15 eq) and Ag_2_O (0.204 g, 0.883 mmol, 2.5 eq). The mixture was stirred at 60 ^o^C for 24 hours. The mixture was allowed to back to room temperature, then DMF was removed *in vacuo*. The crude product was purified by flash chromatography over silica gel using 50% DCM/hexanes (v/v) to give compound **Tz-4** (0.057 g, 74.0%) as the pink solid. R_f_ = 0.38 (50% DCM/Hexanes, visualized w/ UV). ^1^H NMR (400 MHz, CDCl_3_): = 9.44 (s, 1H), 8.92 (d, *J* = 8 Hz, 1H), 8.46 (d, *J* = 8 Hz, 1H), 7.79 (t, *J* = 8 Hz, 1H), 3.15 (s, 3H). ^13^C NMR (101 MHz, CDCl_3_): δ = 168.41, 162.92, 149.27, 133.88, 133.52, 130.66, 127.13, 123.06, 21.52. LRMS (ESI+): Calcd C_9_H_8_N_5_O_2_+ [M+H]^+^: 218.2, Found: 218.2.

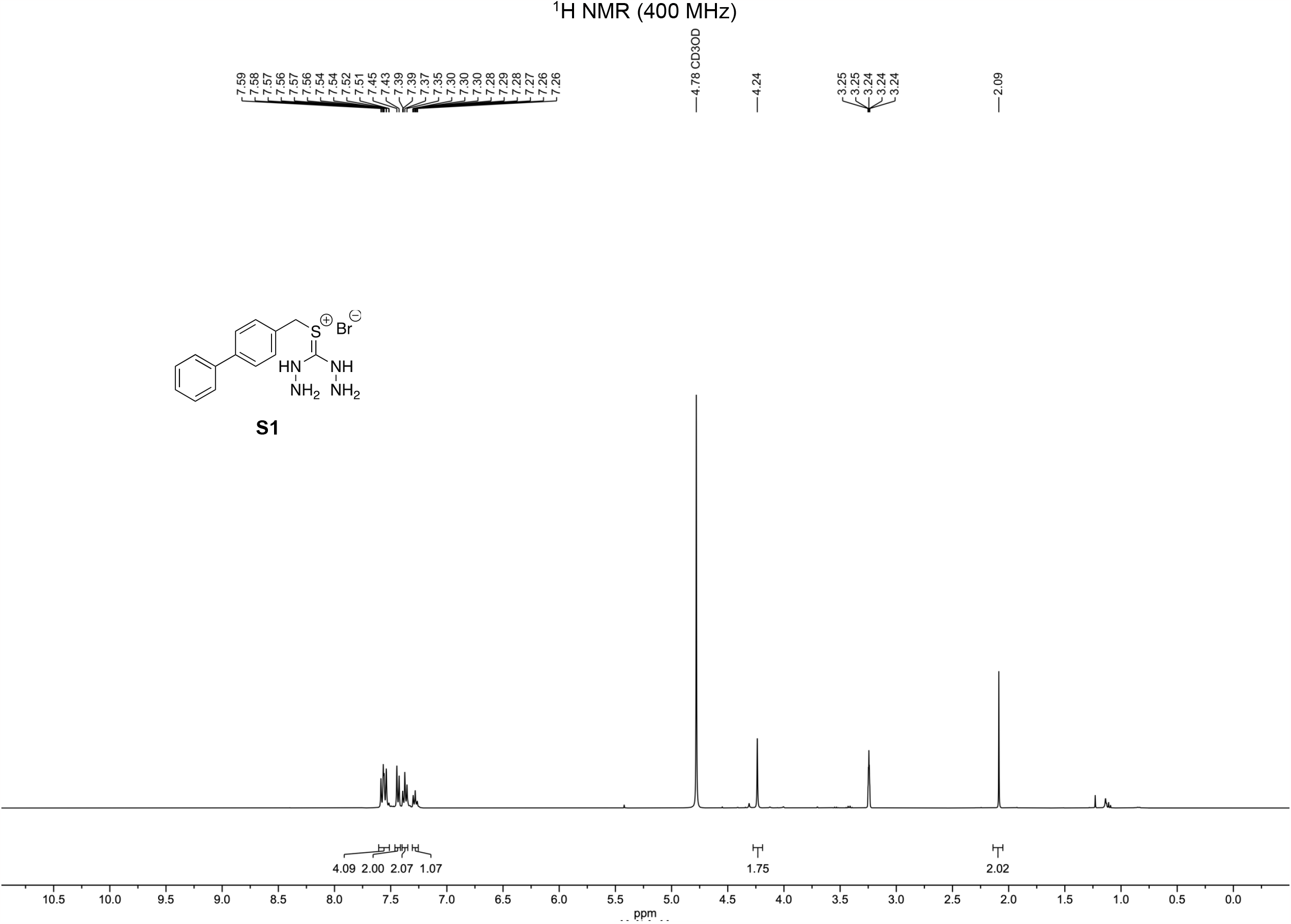

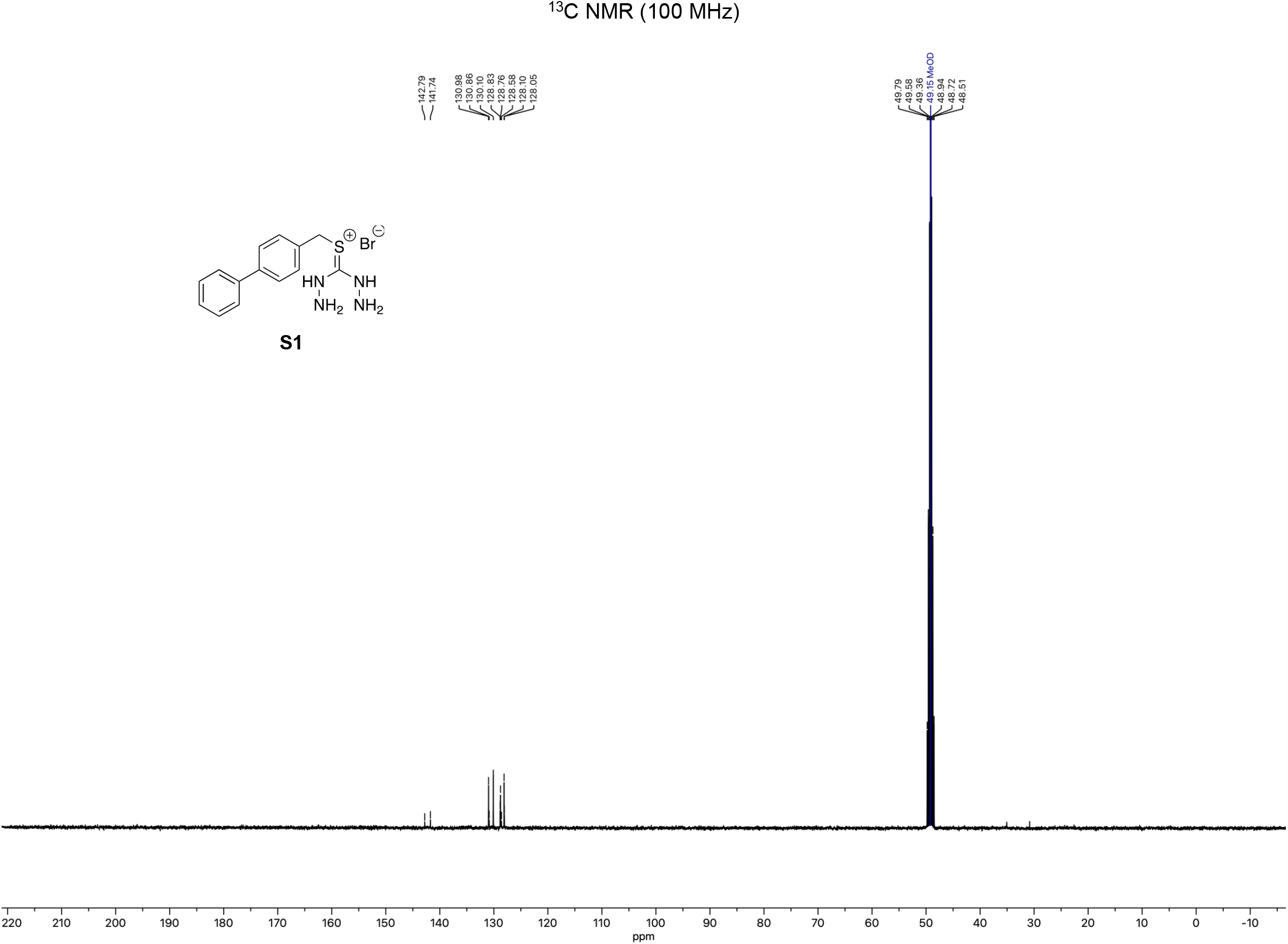

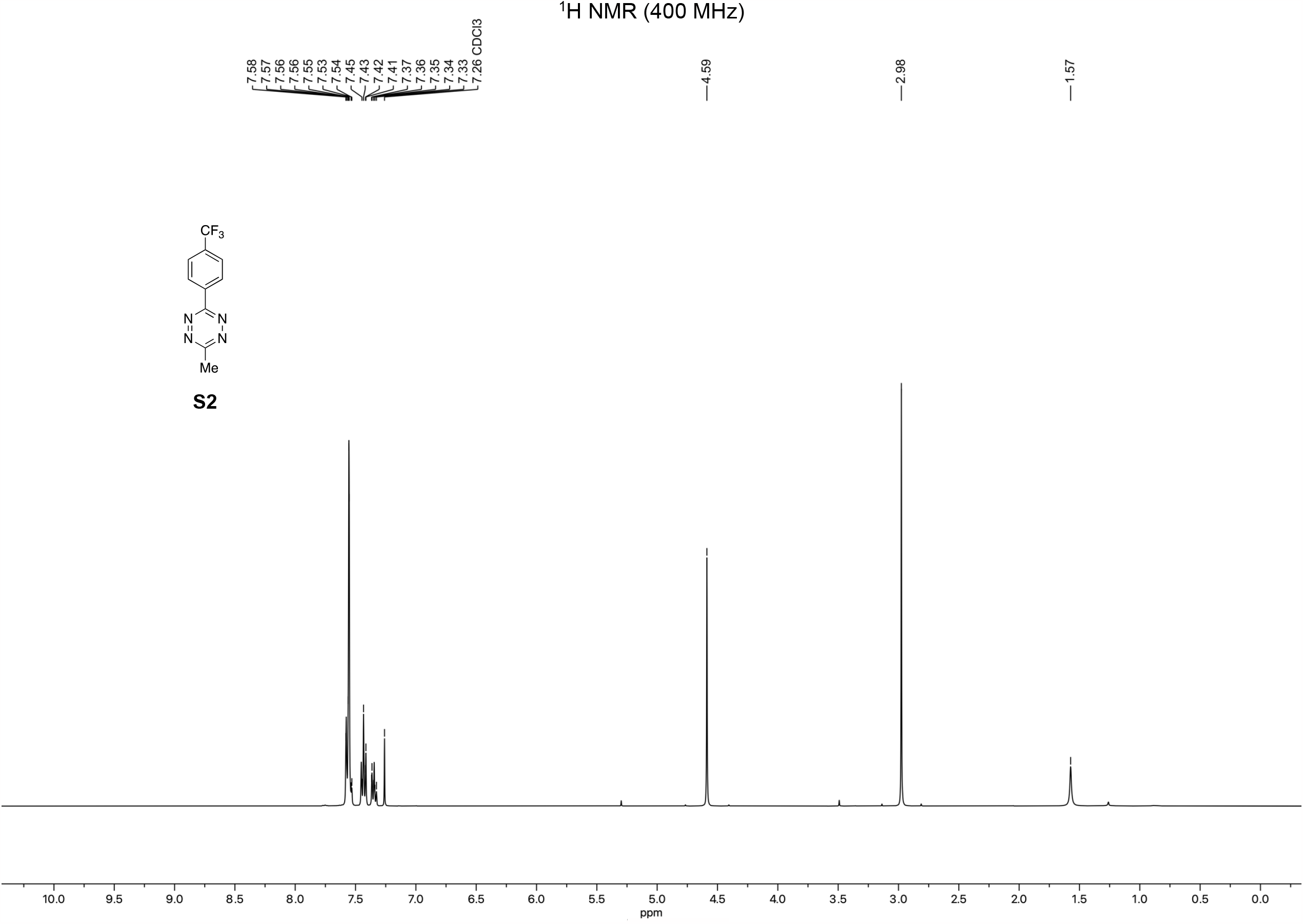

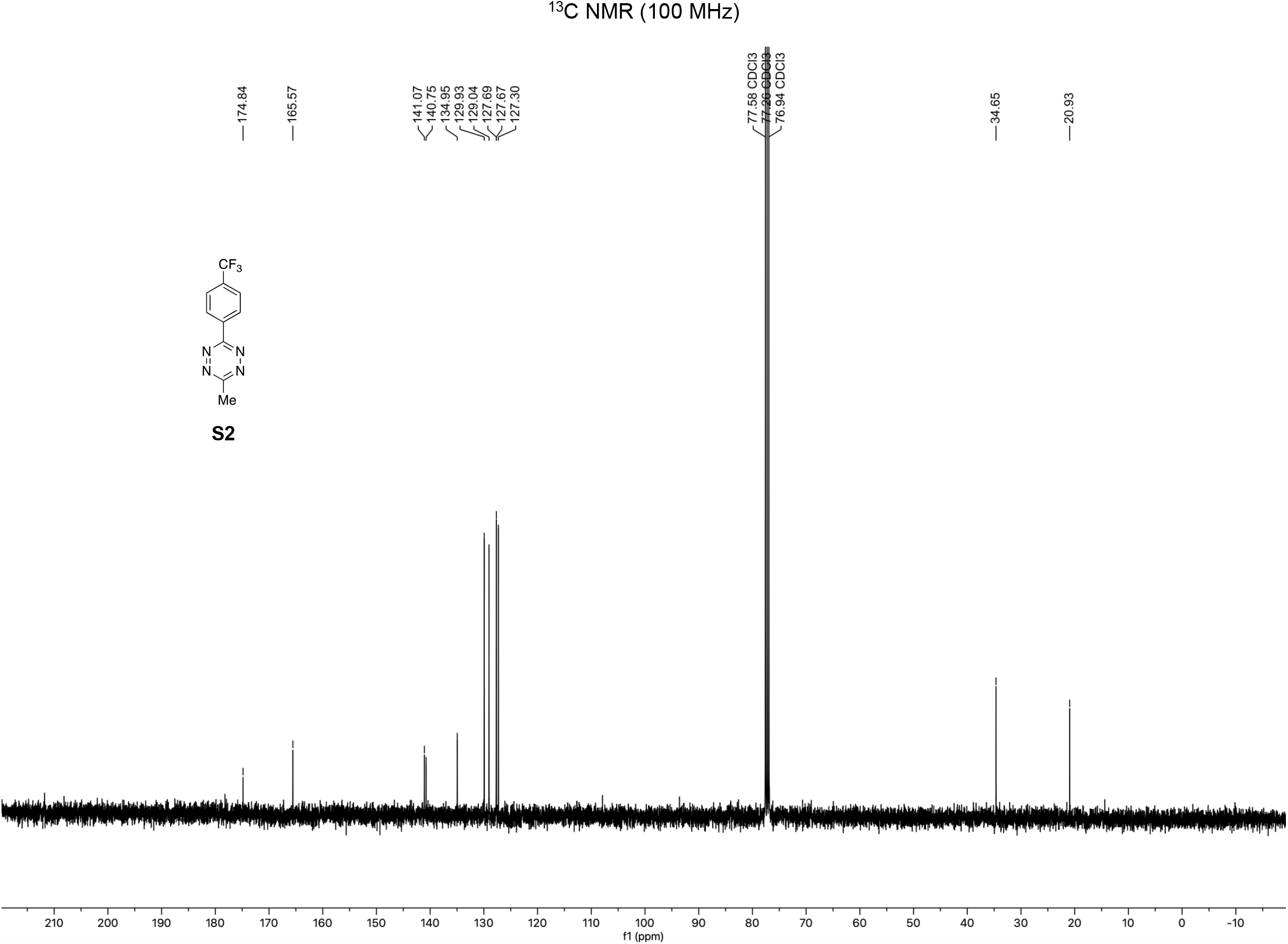

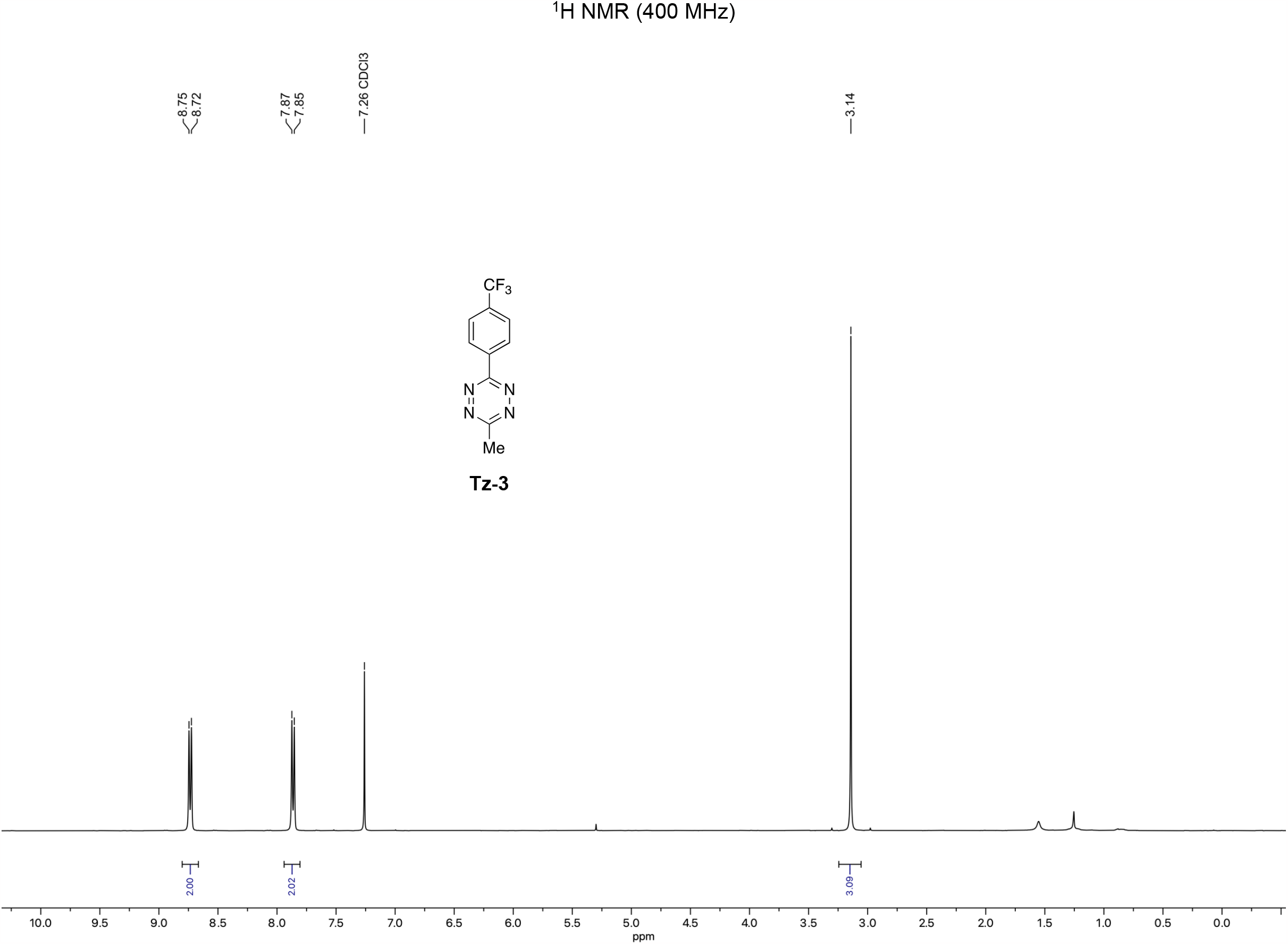

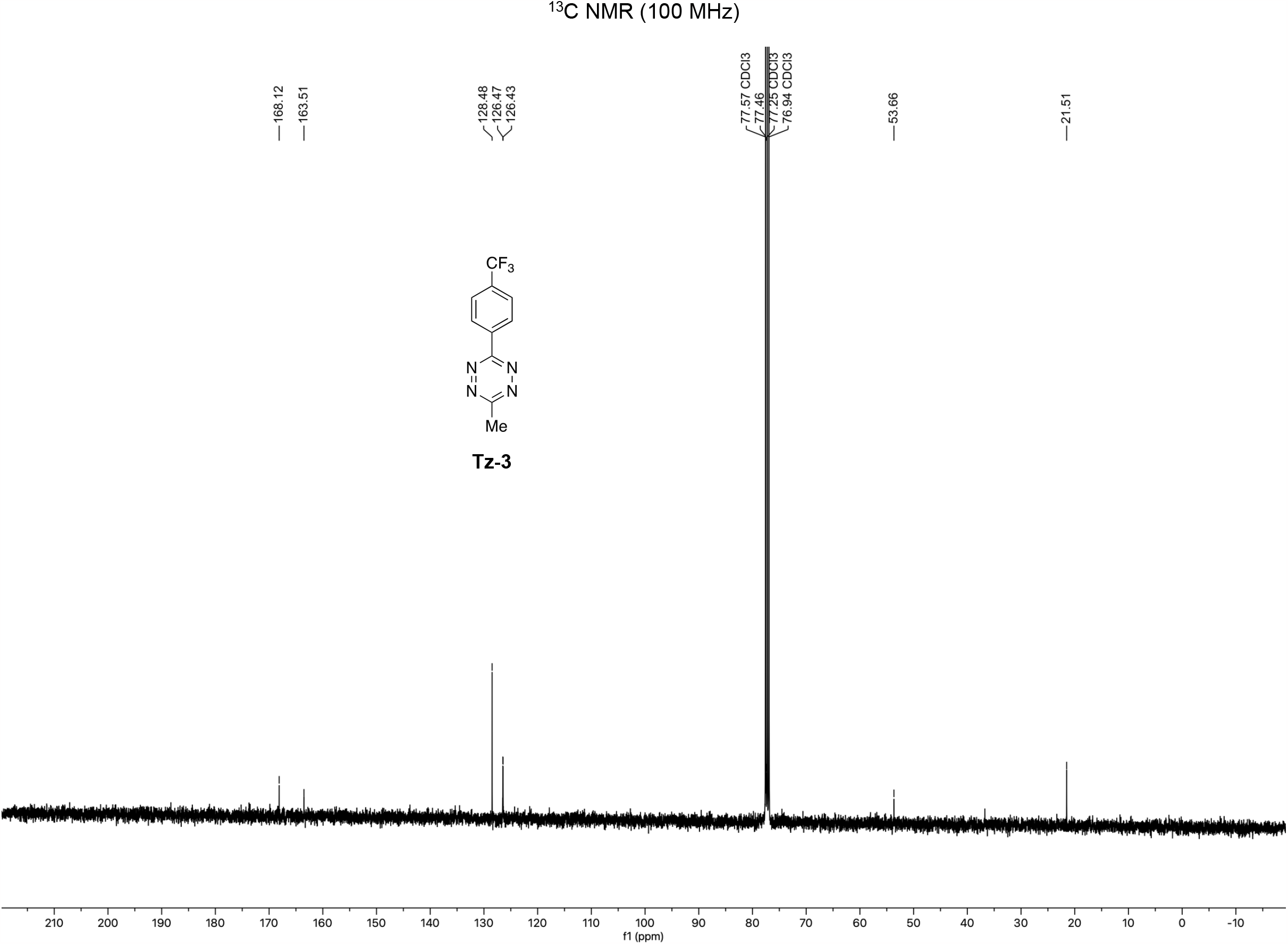

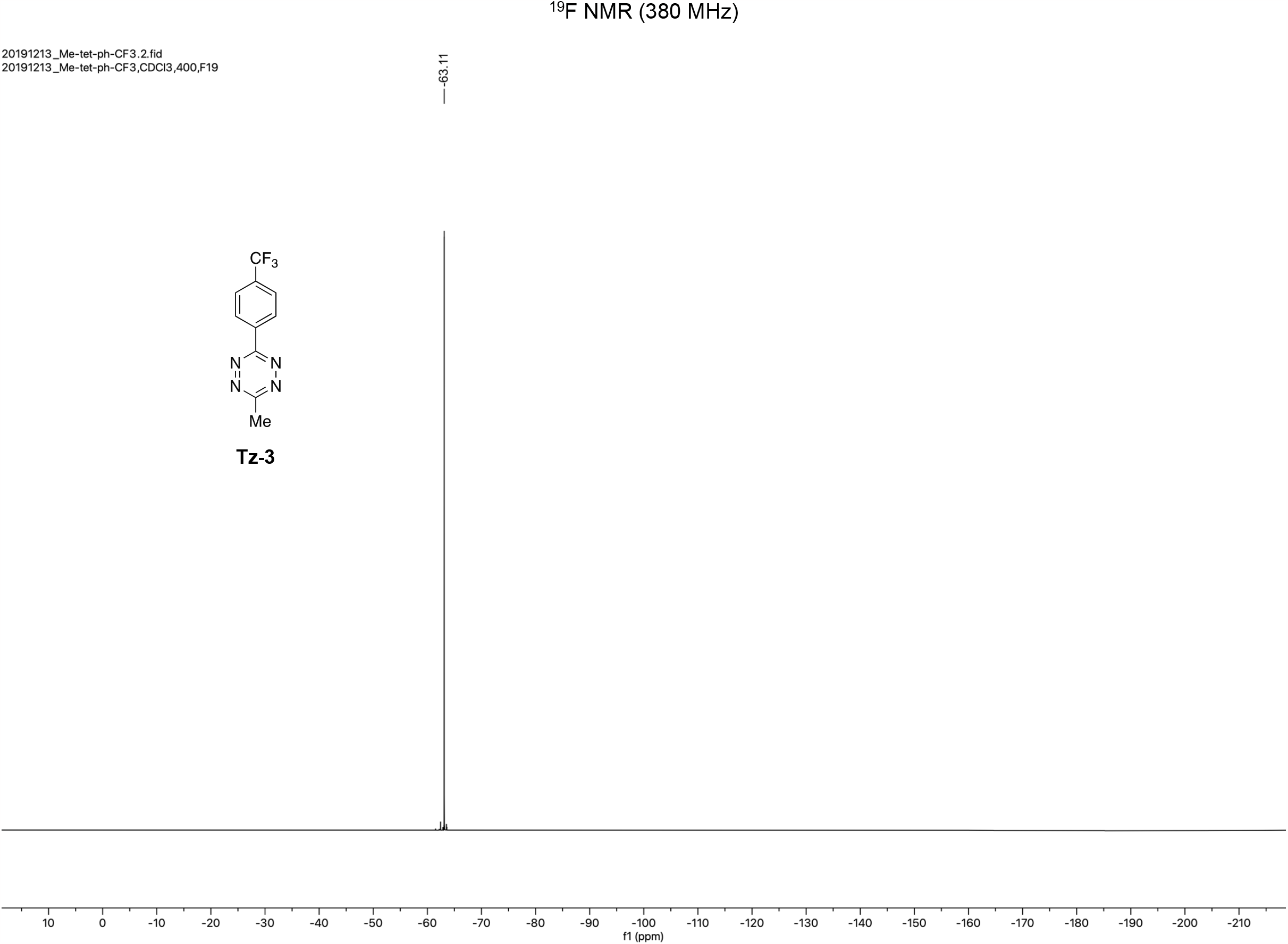

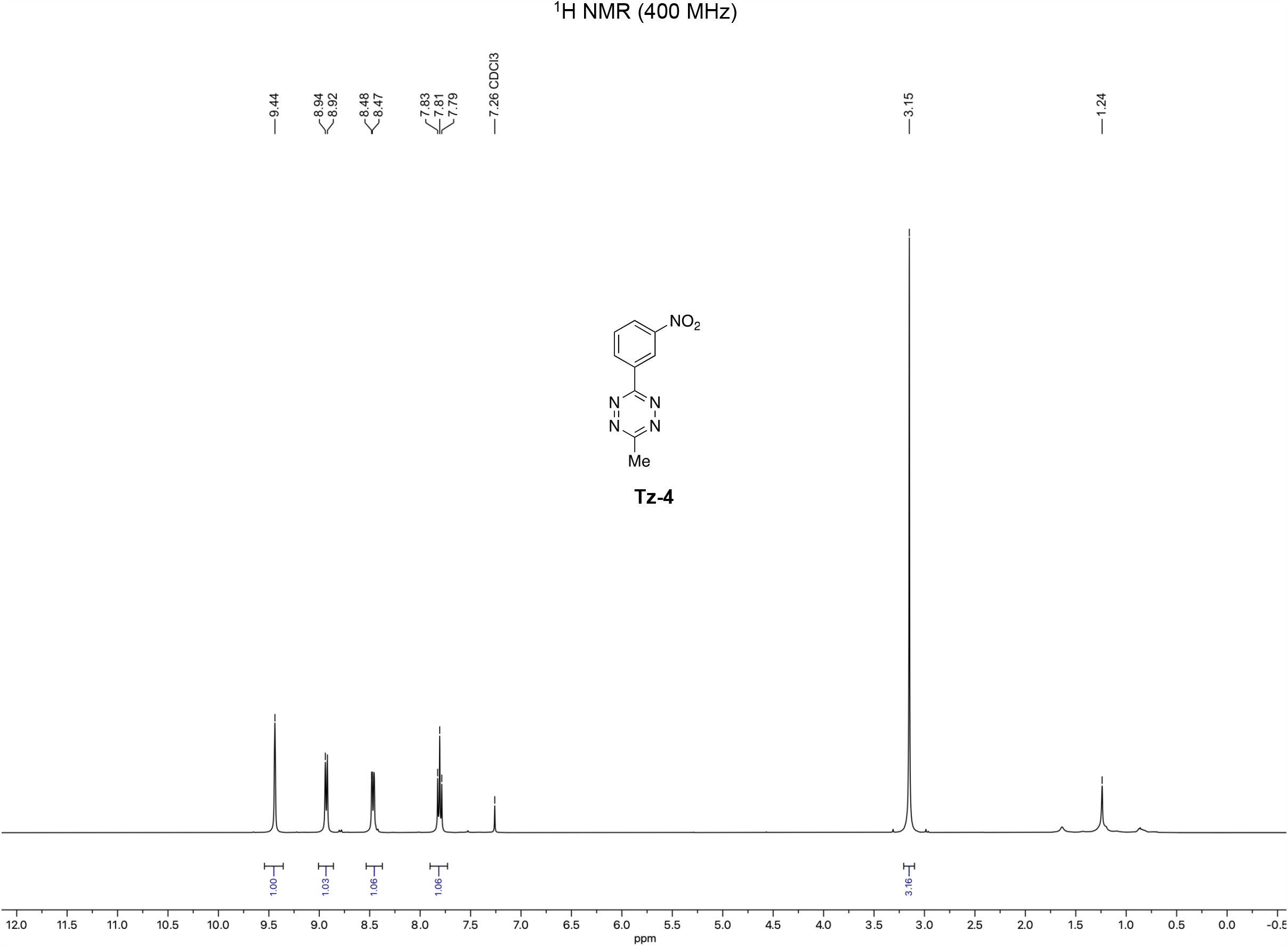

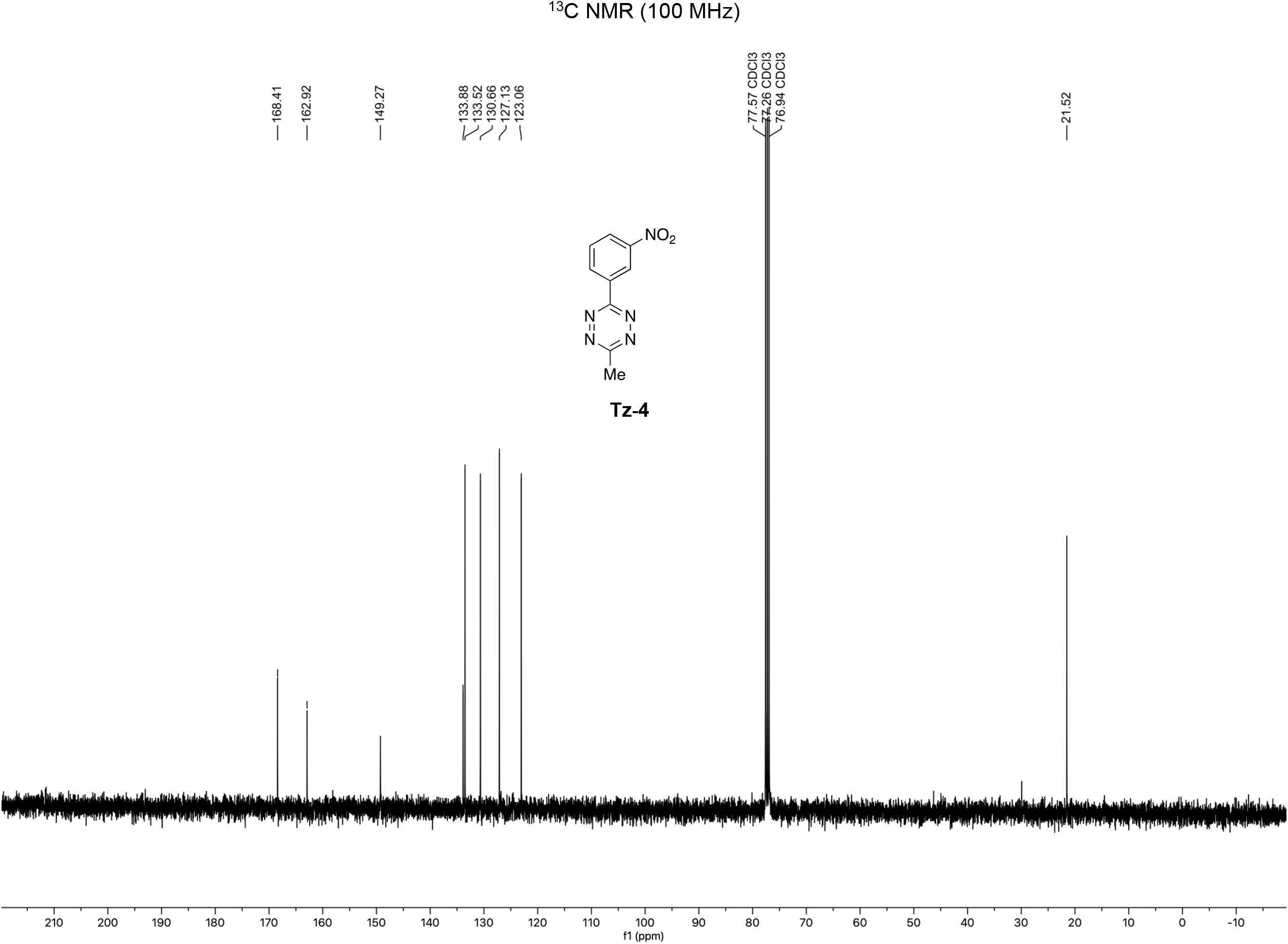

**Scheme 5:**
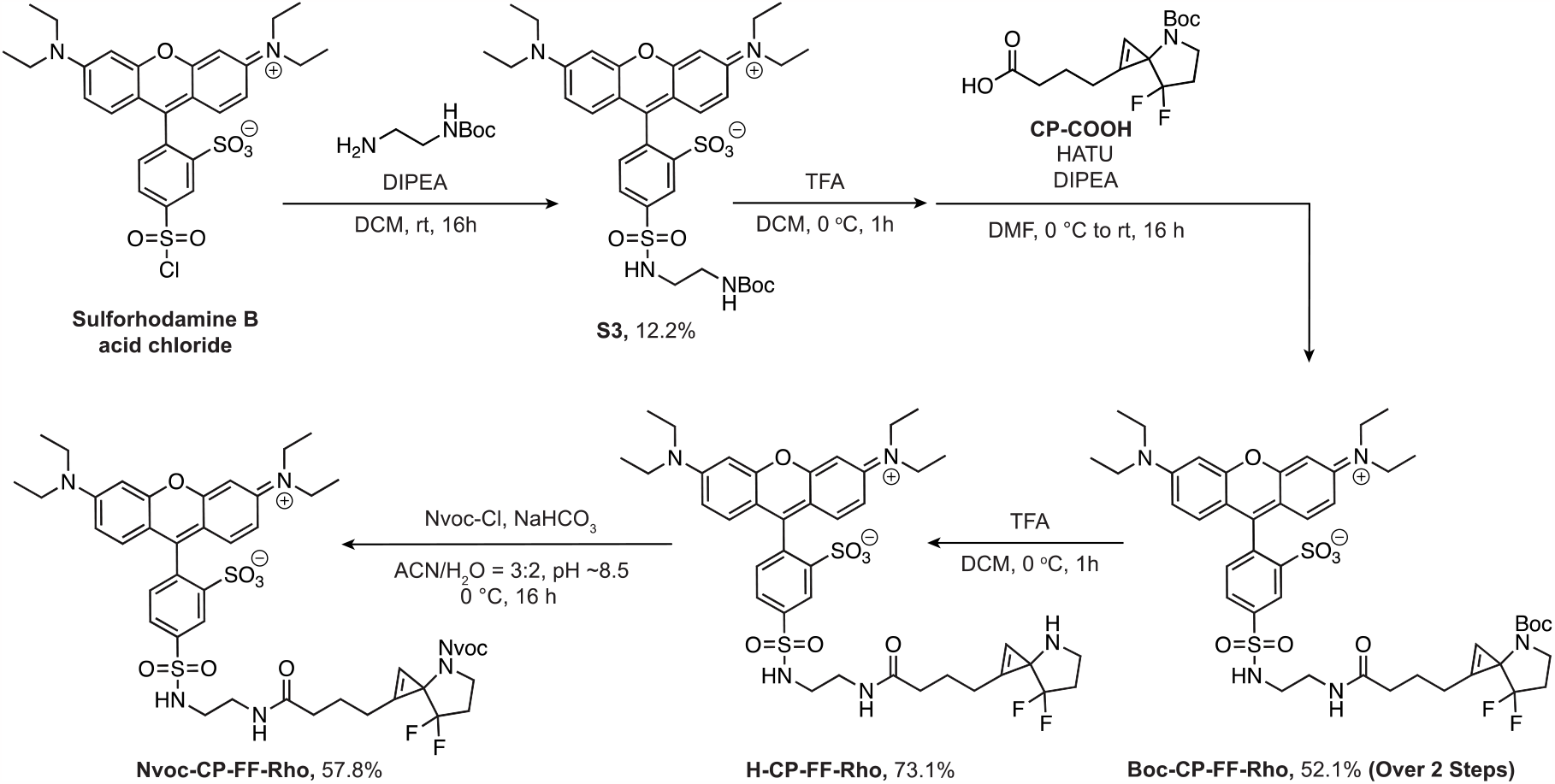
Methods for synthesis of compounds S3, Boc-CP-FF-Rho, H-CP-FF-Rho, & Nvoc-CP-FF-Rho.

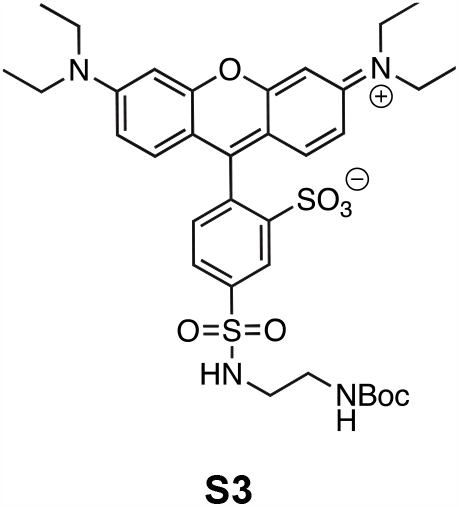

To an ice-cold suspension of Sulforhodamine B acid chloride (88 mg, 0.15 mmol, 1 eq) in DCM (5 mL) was added *N*-Boc-ethylenediamine (33 mg, 0.21 mmol, 1.4 eq) and DIPEA (0.131 ml, 0.75 mmol, 5 eq). The mixture was stirred at room temperature overnight, then DCM was removed by a rotary evaporator. The crude mixture was purified by flash column chromatography on silica gel using 10% MeOH/DCM (v/v) to obtain compound **S3** (13 mg, 12.2%). R_f_ = 0.52 (10% MeOH/DCM, visualized w/ UV). ^1^H NMR (700 MHz, CD_3_OD) δ = 8.6 (s, 1H), 8.22 (d, *J* = 5.6 Hz, 1H), 7.56 (d, *J* = 7.7 Hz,1H), 7.14 (d, *J* = 9.8 Hz, 2H), 7.08 (d, *J* = 9.8 Hz, 2H) 6.98 (s, 7.2, 2H), 3.73-3.66 (m, 8H), 3.03-3.00 (m, 2H), 2.87-2.85 (m, 2H), 1.38 (s, 9H), 1.30 (t, *J* = 8.4 Hz, 12H). ^13^C NMR (176 MHz, CDCl_3_): δ = 159.46, 158.51, 157.42, 156.52, 149.41, 142.47, 133.96, 133.22, 132.97, 130.91, 127.76, 115.64, 115.42, 97.32, 80.29, 47.04,45.51, 43.74, 42.06, 41.70, 33.22, 30.89, 28.86, 28.25, 23.88, 14.59, 13.01, 9.39. HRMS (ESI): Calcd C_34_H_45_N_4_O_8_S_2_+ [M+H]^+^: 701.2601, Found: 701.2668.

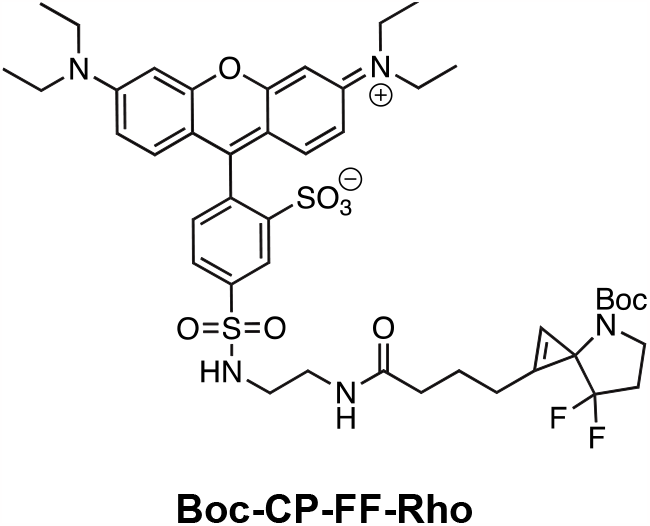

To an ice-cold suspension of **S3** (13 mg, 0.02 mmol, 1 eq) in DCM (5 mL) was added TFA (1 mL). The mixture was stirred at rt for an hour, and then the solvent was removed by a rotary evaporator. The crude mixture was collected without further purification to obtain the Boc-deprotected rhodamine. To an ice-cold suspension of cyclopropene **CP-COOH** (5.7 mg, 0.018 mmol, 1 eq) in DMF (2 mL) was added HATU (8.4 mg, 0.022 mmol, 1.2 eq) and DIPEA (4 μL, 0.02 mmol, 1.2 eq). The mixture was stirred at 0 ^o^C for 10 min, and the Boc-deprotected rhodamine (13.3 g, 0.022 mmol, 1.2 eq) was dissolved in DMF (200 μL) and slowly added into the solution at 0 ^o^C. The mixture was allowed to stir at room temperature overnight, then the reaction was extracted with DCM and water, followed by brine washing. The combined organic layer was dried over anhydrous Na_2_SO_4_, concentrated *in vacuo*, and purified by flash chromatography over silica gel on silica gel using 5% MeOH/DCM (v/v) to give compound **Boc-CP-FF-Rho** (8.5 g, 52.1%) as a pink solid. R_f_ = 0.16 (10% MeOH/DCM). Alternatively, the crude product can be purified by HPLC (R_t_ = 24.6 min, 20-100% ACN over 25 mins, flow rate = 1 mL/min). ^1^H NMR (700 MHz, CD_3_OD) δ = 8.58 (s, 1H), 8.21 (d, *J* = 7 Hz, 1H), 7.51 (d, *J* = 7 Hz, 1H), 7.11 (d, *J* = 14 Hz, 2H), 7.06 (d, *J* = 14 Hz, 2H), 8.87 (s,1H), 3.71-3.67 (m, 8H), 3.53-3.47 (m, 4H), 3.13 (t, *J* = 7 Hz, 2H), 2.91 (t, *J* = 7 Hz, 2H), 2.50-2.47 (m, 2H), 2.21-2.18 (m, 2H), 1.93-1.90 (m, 2H), 1.84-1.80 (m, 2H), 1.41 (s, 9H), 1.29 (t, *J* = 7 Hz, 12H). ^13^C NMR (176 MHz, CD_3_OD): δ = 159.32, 157.30, 149.32, 133.84, 133.08, 132.84, 130.77, 127.54, 115.53, 115.28, 97.16, 49.53, 47.95, 46.89, 43.31, 40.40, 36.17, 34.03, 33.08, 30.76, 30.48, 28.75, 28.73, 24.09, 23.89, 23.74, 23.35, 14.43, 12.83, 9.21. HRMS (ESI): Calcd C_44_H_56_F_2_N_5_O_9_S_2_+ [M+H]^+^: 900.3409, Found: 900.3482.

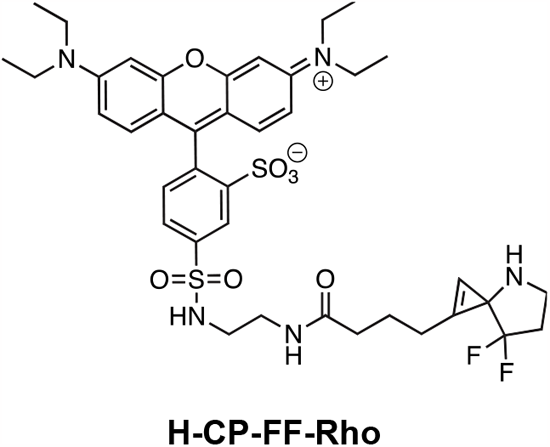

To an ice-cold suspension of **Boc-CP-FF-Rho** (9.2 mg, 0.010 mmol, 1 eq) in DCM (2 mL) was added TFA (2 mL). The mixture was stirred at 0 ^o^C for 1 to 2 hours, and then the reaction mixture was concentrated *in vacuo*. The crude was re-suspended in ACN (0.5 mL) and H_2_O (0.5 mL), and purified by HPLC (R_t_ = 19.7 min, 20-100% ACN over 25 mins, flow rate = 1 mL/min) to obtain **H-CP-FF-Rho** (6.0 mg, 73.1%) as dark red solid. ^1^H NMR (700 MHz, CD_3_OD) δ = 8.57 (s, 1H), 8.22 (d, *J* = 14 Hz, 1H), 7.45 (d, *J* = 14 Hz, 1H), 7.26 (s, 2H), 7.10–7.06 (m, 4H), 6.97 (s, 2H), 6.83 (s, 1H), 3.72– 3.64 (m, 8H), 3.58–3.43 (m, 4H), 3.24–3.15 (m, 4H), 2.98 (t, *J* = 14 Hz, 2H), 2.71–2.63 (m, 4H), 2.30 (t, *J* = 14 Hz, 2H), 1.99 (m, 4H), 1.33 (m, 12H). ^13^C NMR (176 MHz, CD_3_OD): δ = 157.93, 155.90, 131.64, 129.37, 126.21, 114.11, 113.85, 95.77, 48.12, 48.00, 46.54, 45.48, 39.45, 38.96, 34.30, 32.76, 31.67, 29.35, 29.20, 29.07, 29.03, 28.84, 22.48, 22.33, 13.03, 11.41, 7.80. HRMS (ESI): Calcd C_39_H_48_F_2_N_5_O_7_S_2_+ [M+H]^+^: 800.2957, Found: 800.2951.

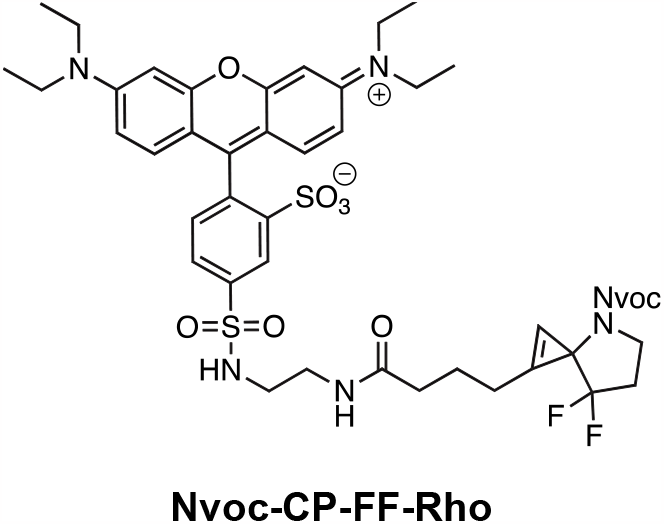

To an ice-cold suspension of **H-CP-FF-Rho** (4.4 mg, 0.005 mmol, 1 eq) in ACN (3 mL) and H_2_O (2 mL) was added sodium carbonate to adjust pH to ∼8.5, and 4,5-dimethoxy-2-nitrobenzyl chloroformate (7.6 mg, 0.027 mmol, 5 eq) at 0 ^o^C. The mixture was allowed to stir at room temperature overnight. The reaction mixture was concentrated *in vacuo*, re-suspended in ACN (0.2 mL) and H_2_O (0.2 mL), and purified by HPLC (R_t_ = 22.7 min, 30-100% CAN over 25 mins, flow rate = 1 mL/min) to obtain **Nvoc-CP-FF-Rho** (3.3 mg, 57.8%) as pink solid. ^1^H NMR (700 MHz, CD_3_OD) δ = 8.57 (s, 1H), 8.21 (d, *J* = 14 Hz, 1H), 7.70 (s, 1H), 7.70–7.52 (m, 2H), 7.51 (d, *J* = 7 Hz, 1H), 7.11 (d, *J* = 7 Hz, 2H), 7.07 (d, *J* = 7 Hz, 2H), 6.92 (s, 2H), 3.89 (s, 6H), 3.69–3.66 (m, 8H), 3.14–3.12 (m, 2H), 2.92 (d, *J* = 7 Hz, 2H), 2.42–2.35 (m, 6H), 2.04–1.5 (m, 8H), 1.31 (t, *J* = 7 Hz, 12H). ^13^C NMR (176 MHz, CD_3_OD) δ 159.25, 157.27, 157.25, 156.46, 155.00, 149.31, 142.39, 133.82, 133.02, 132.85, 130.76, 127.53, 115.54, 115.52, 115.24, 115.22, 109.50, 97.17, 97.16, 56.88, 49.53, 49.40, 46.88, 43.36, 42.18, 40.38, 35.98, 33.08, 30.75, 30.47, 23.98, 23.74, 14.43, 12.83. HRMS (ESI): Calcd C_49_H_57_F_2_N_6_O_13_S_2_+ [M+H]^+^: 1039.3315, Found: 1039.3385.

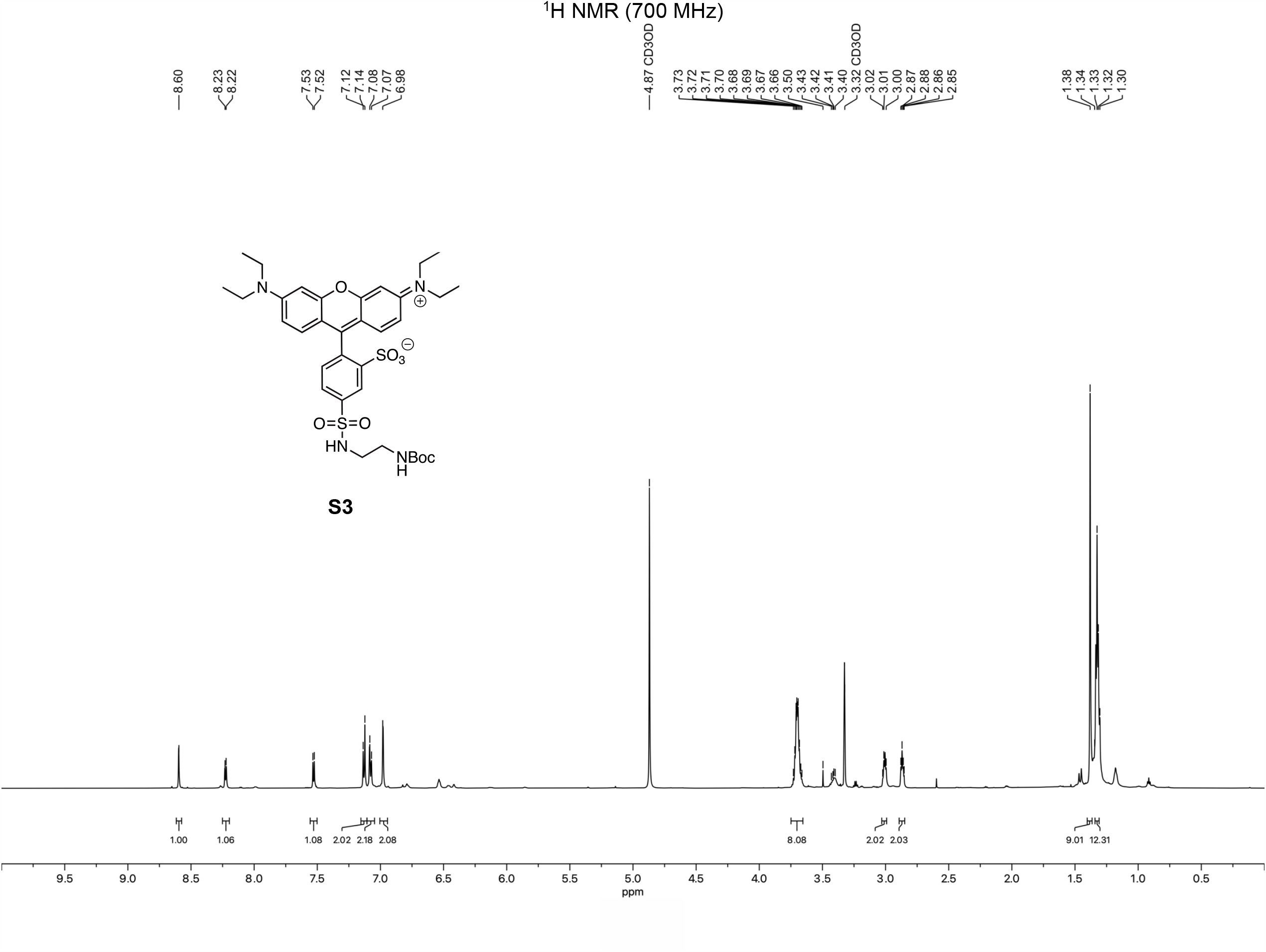

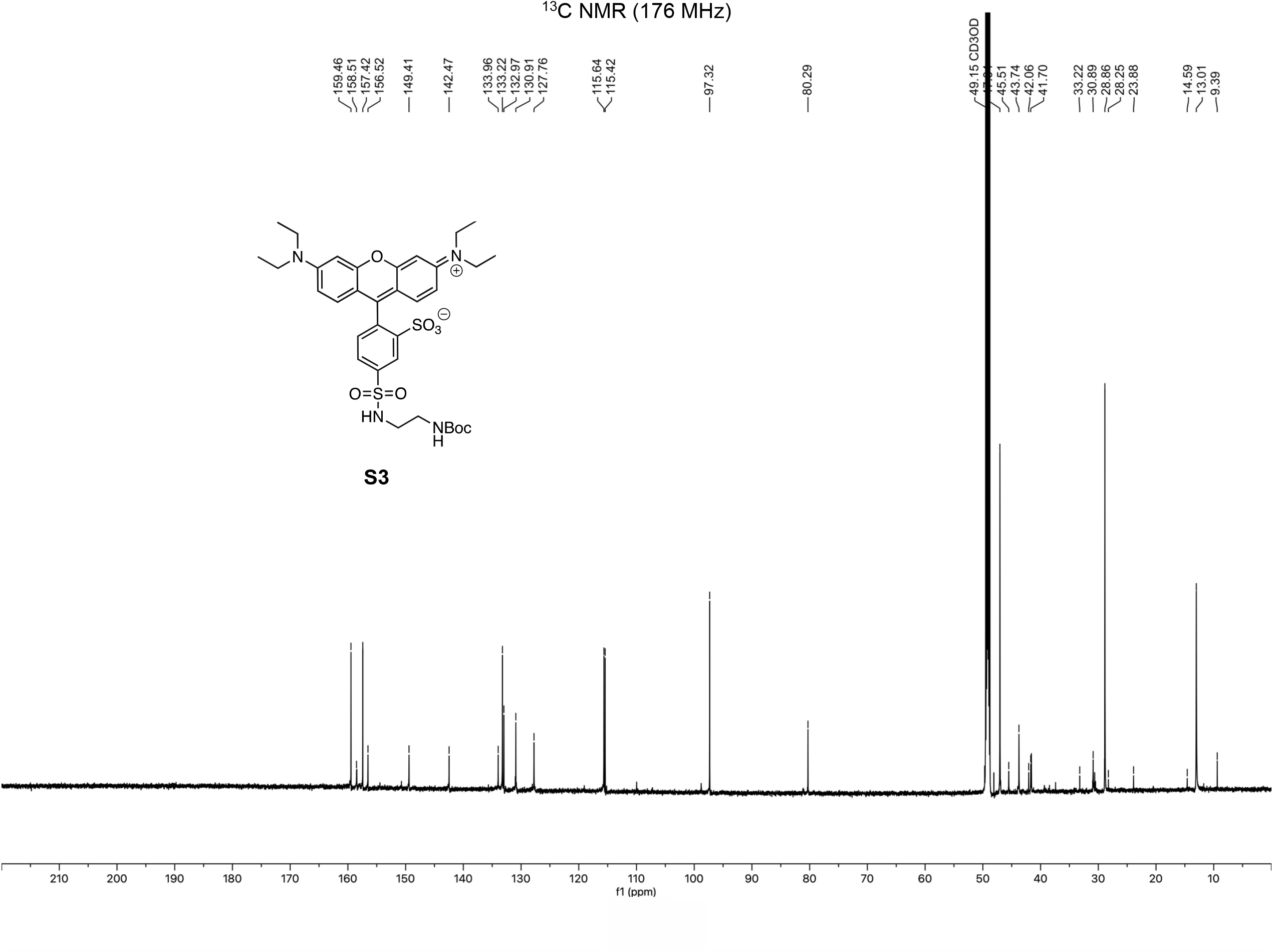

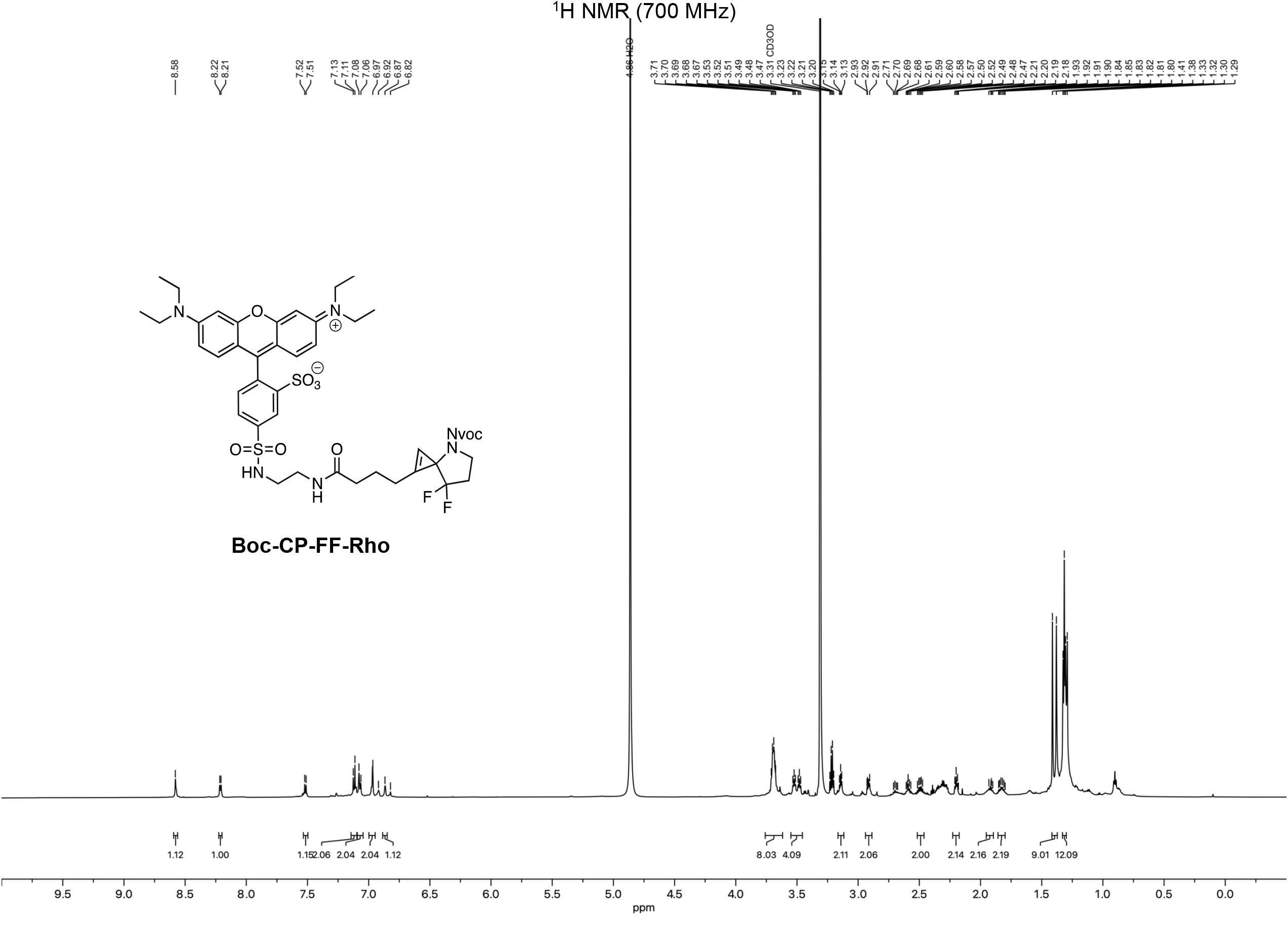

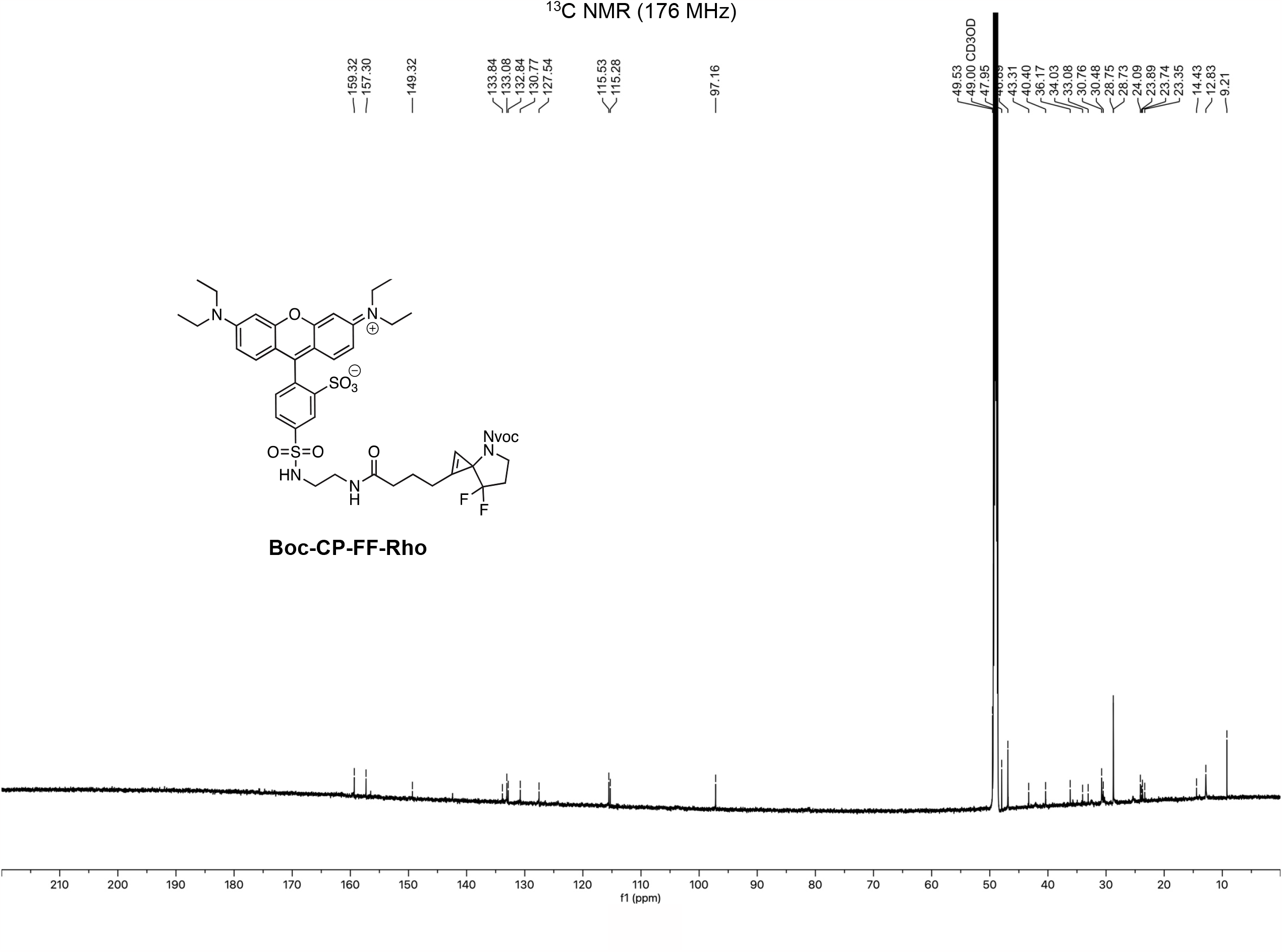

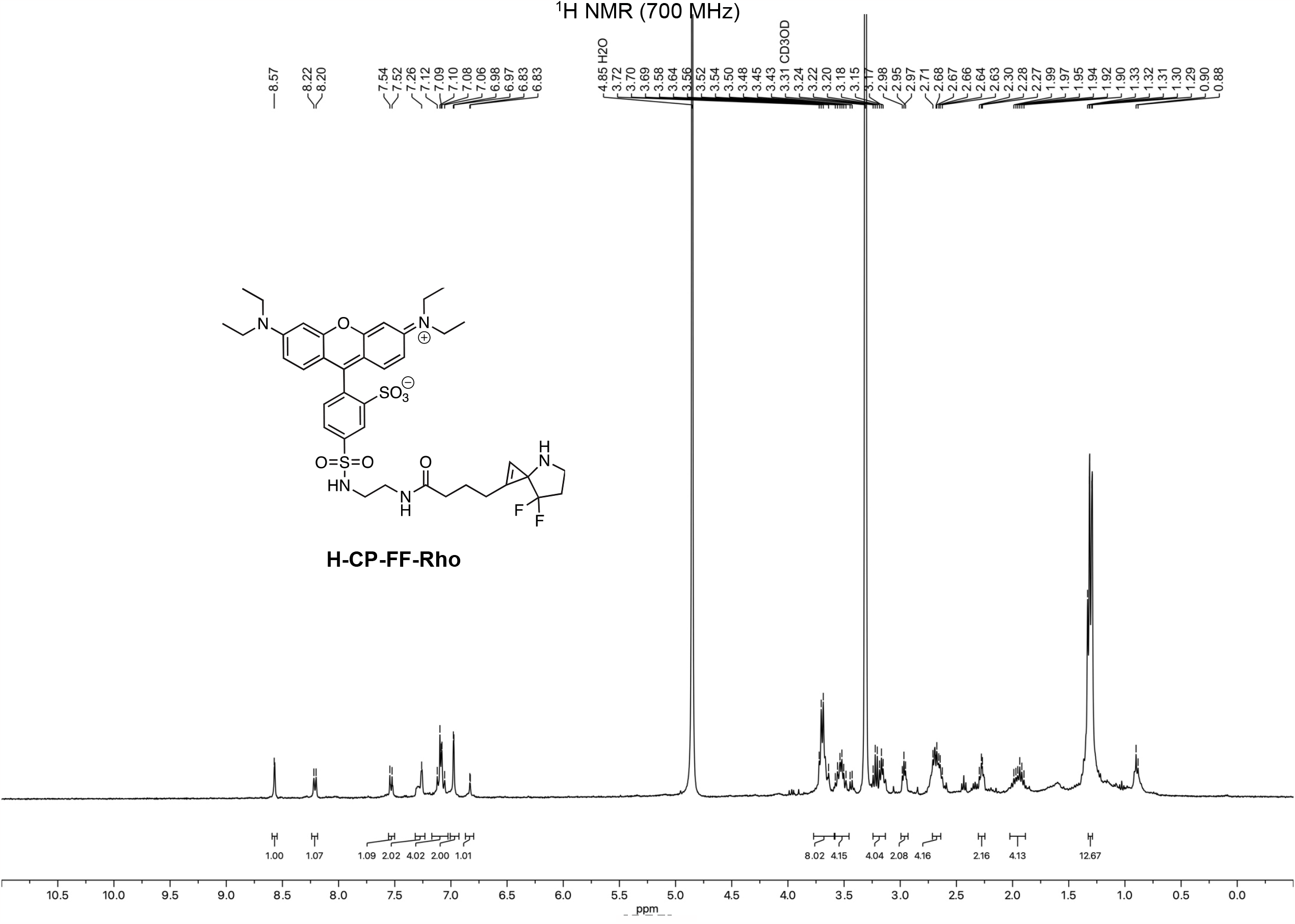

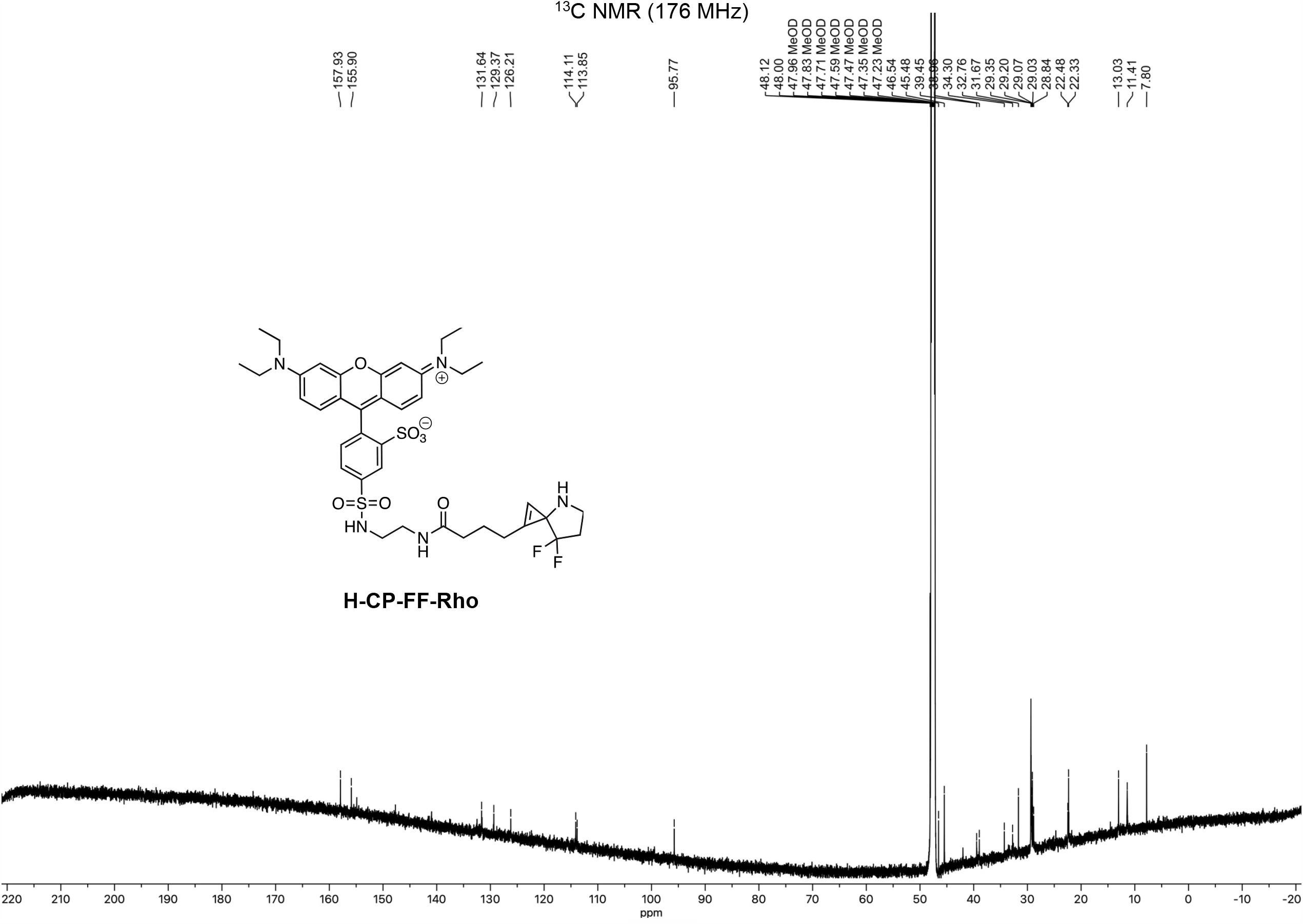

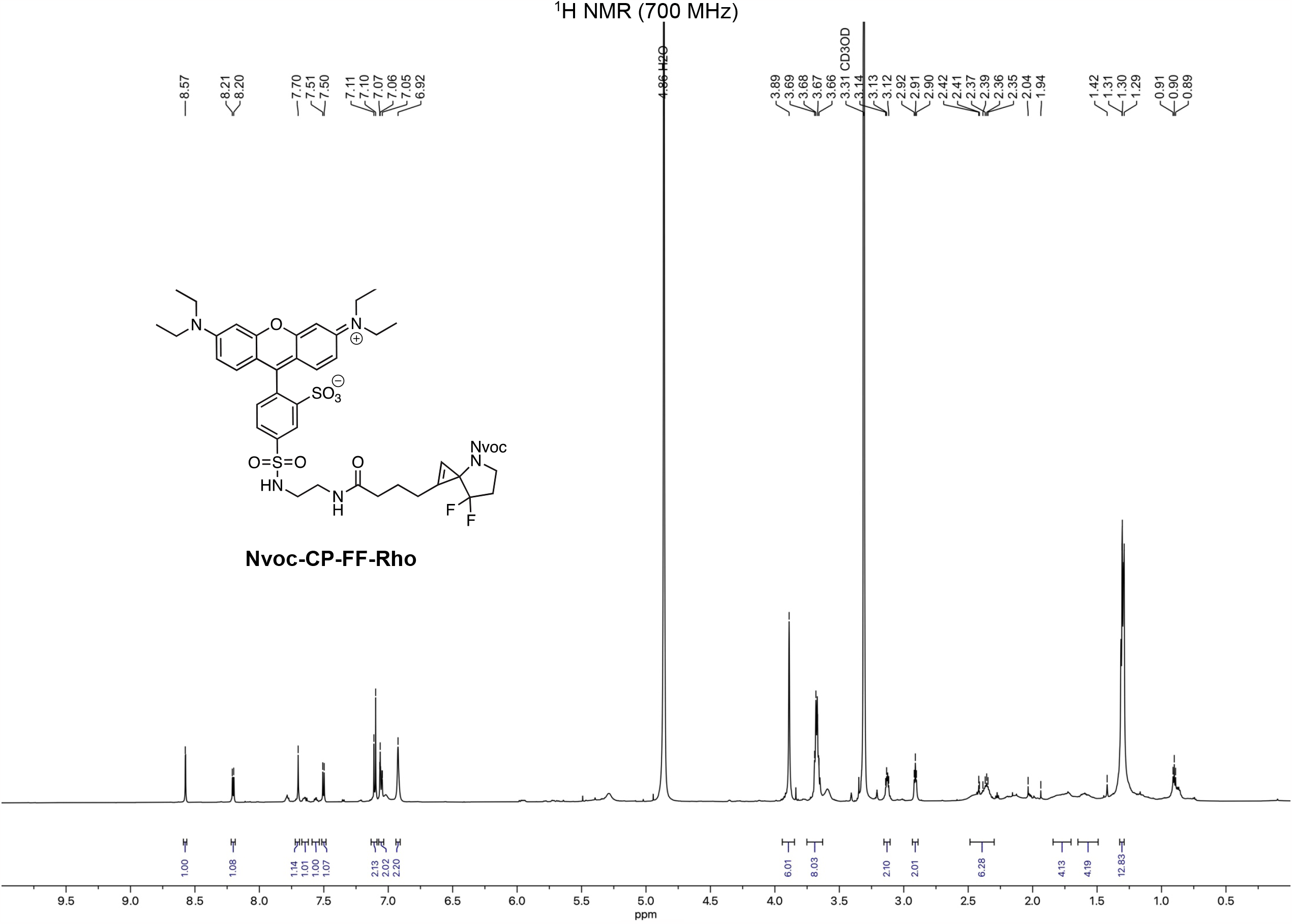

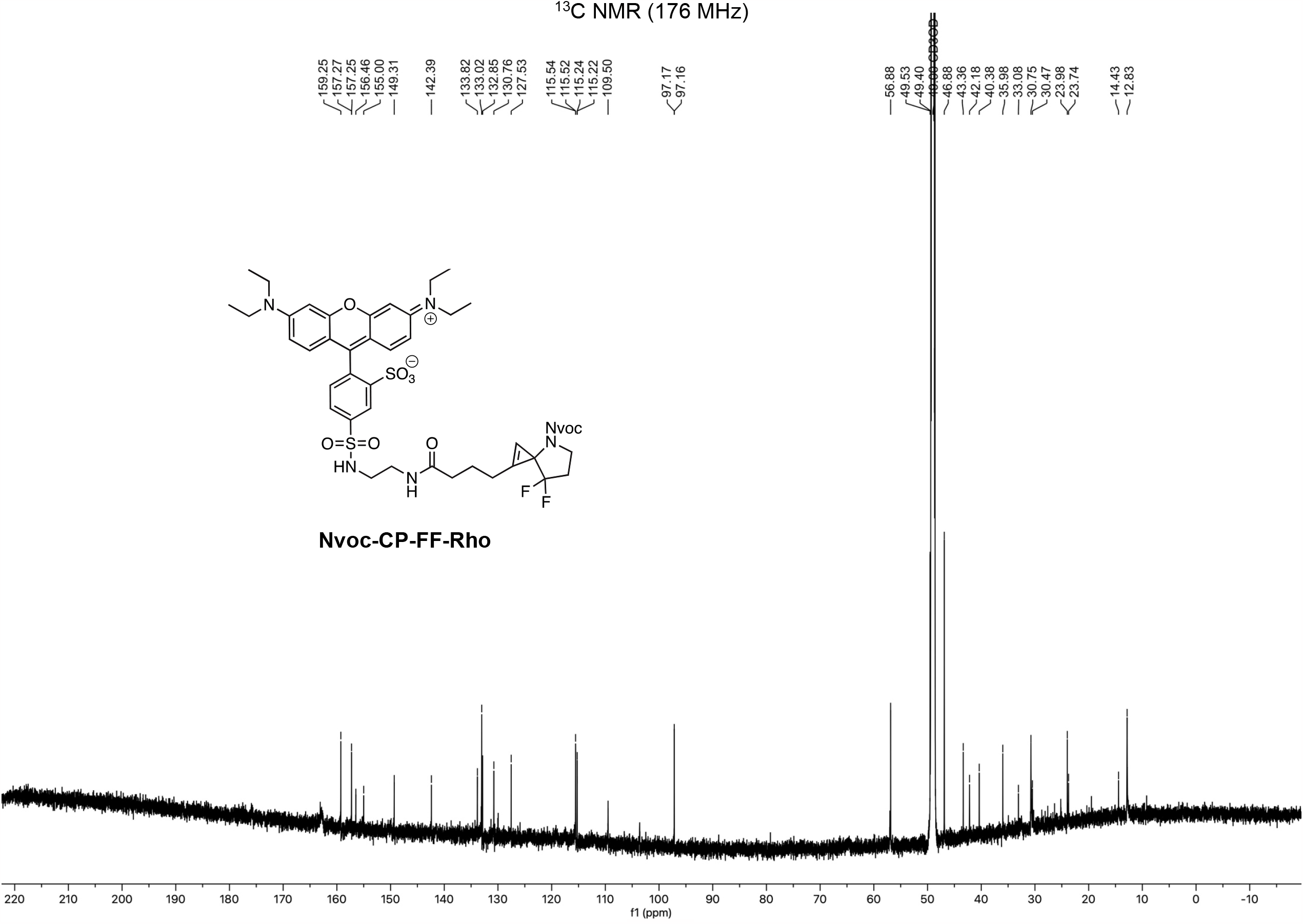

**Scheme 6:**
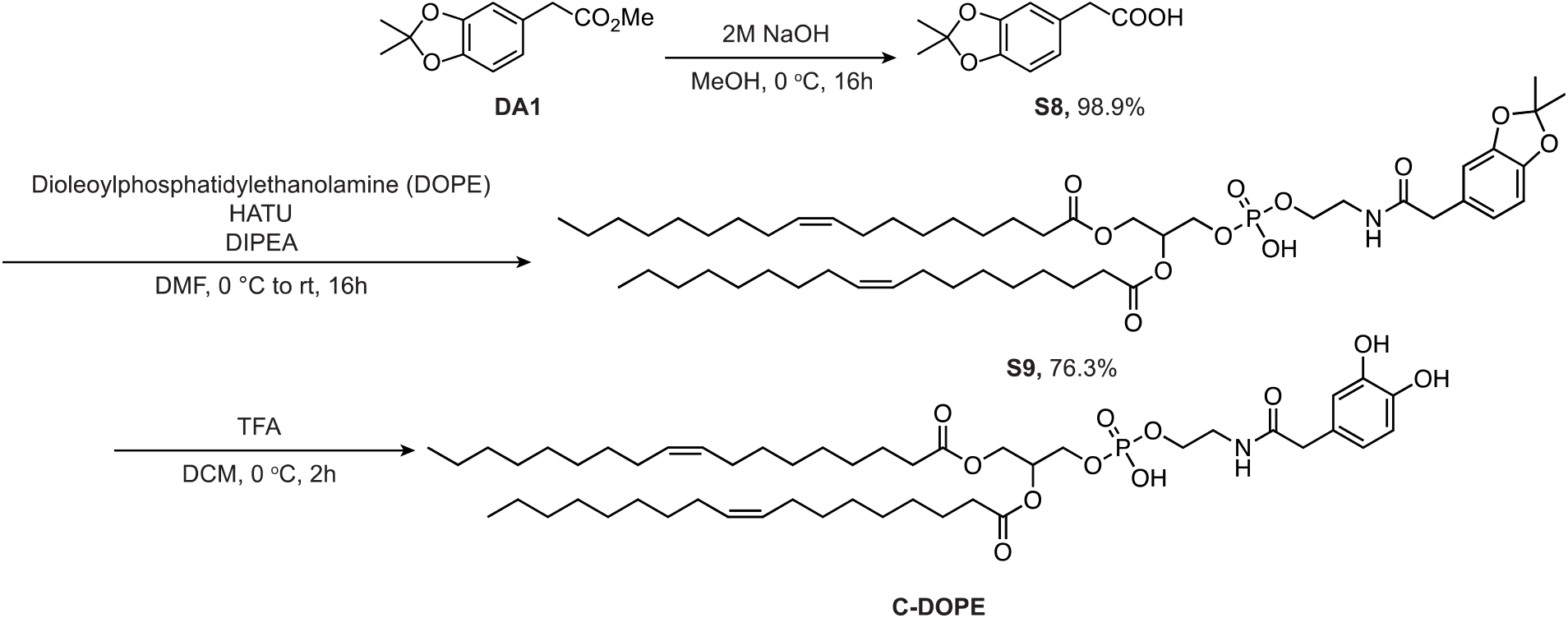
Methods for synthesis of compounds S8, S9 and C-DOPE.

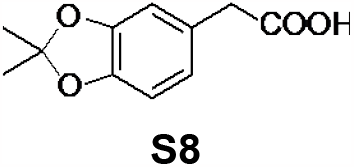

To an ice-cold solution of compound **DA1** (100 mg, 0.45 mmol) in MeOH (2 mL) at room temperature was added 2M aqueous NaOH (2 mL). The reaction was stirred for 16 hours, and the solvent was removed under a vacuum. The crude product is purified by flash chromatography over silica gel (50% EtOAc/hexanes (v/v)) to obtain **S8** (92.66 mg, 98.9%). ^1^H NMR (400 MHz, CDCl_3_) δ = 6.68 (s, 1H), 6.67 (s, 2H), 3.54 (s, 2H), 1.66 (6H). ^13^C NMR (176 MHz, CDCl_3_) δ = 178.16, 147.89, 146.99, 126.31, 122.15, 118.33, 109.81, 108.38, 40.91, 26.11. MS(ESI): Calcd C_11_H_13_O_4_+ [M+H]^+^: 208.1, found: 208.1.

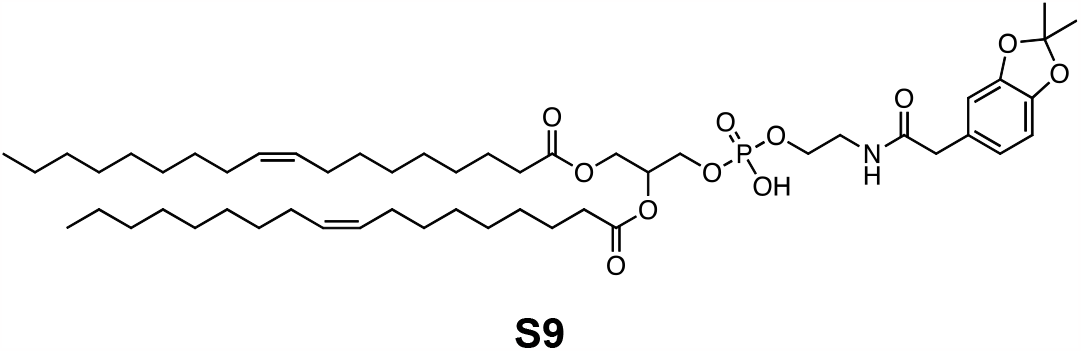

To an ice-cold suspension of compound **S8** (17 mg, 0.084 mmol, 1.2 eq) in anhydrous DCM (4 mL) was added DCC (14 mg, 0.07 mmol, 1 eq) and DIPEA (24 µL, 0.14 mmol, 2 eq) under N_2_. The mixture was stirred at 0 ^o^C for 10 min, and DOPE (50 mg, 0.07 mmol, 1 eq) in the anhydrous DCM (1 mL) was slowly added into the solution at 0 ^o^C. The reaction was stirred for an additional 16 hours, then the solvent was removed *in vacuo*. The crude product was purified by flash column chromatography on silica gel using 30% EtOAc/hexanes (v/v) to obtain compound **S6** (17 mg, 76.3%). ^1^H NMR (400 MHz, CDCl_3_) δ = 6.69–6.57 (m, 5H), 5.34–5.29 (m, 4H), 5.22 (s, 1H), 4.35–4.32 (m, 1H), 4.13 (t, *J* = 7 Hz, 1H), 3.94–3.90 (m, 2H), 3.67 (s, 2H), 3.41 (s, 2H), 2.25 (t, *J* = 14 Hz, 3H), 2.00–1.97 (m, 5H), 1.92–1.89 (m, 4H), 1.79–1.76 (m, 2H), 1.72–1.67 (m, 4H), 1.65 (s, 3H), 1.62–1.61 (m, 6H), 1.37–1.31 (m, 15H), 1.29–1.26 (m, 15H), 1.18–1.08 (m, 7H), 0.88 (t, *J* = 7 Hz, 4H). ^13^C NMR (176 MHz, CDCl_3_): δ = 173.84, 154.17, 147.94, 147.79, 147.68, 146.72, 146.64, 146.58, 130.18, 129.89, 127.69, 122.17, 122.05, 121.58, 118.24, 117.99, 109.90, 109.46, 108.39, 108.33, 108.17, 77.58, 77.46, 77.26, 76.94, 56.91, 53.65, 50.03, 49.47, 43.01, 34.40, 34.24, 34.04, 32.92, 32.13, 31.07, 30.02, 30.00, 29.76, 29.58, 29.56, 29.54, 29.49, 29.47, 29.43, 29.38, 27.45, 26.58, 26.06, 26.04, 25.80, 25.72, 25.53, 25.13, 25.05, 24.94, 22.90, 14.33. HRMS (ESI): Calcd C_52_H_88_NO_11_P [M+H]^+^: 934.6167, Found: 934.6153.

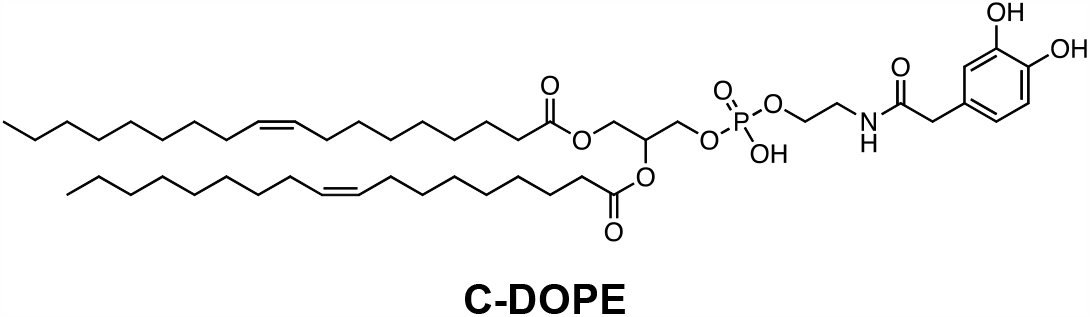

To an ice-cold suspension of compound **S9** (62 mg, 0.069 mmol, 1.2 eq) in anhydrous DCM (5 mL) was added TFA (5 mL). The mixture was stirred at 0 ^o^C for an hour. The solvent and TFA were removed *in vacuo* to obtain deprotected DOPE-linked catechol compound **C-DOPE**. The crude product was used without further purification. HRMS (ESI): Calcd C_49_H_85_NO_11_P+ [M+H]^+^: 894.5854, Found: 894.5856.

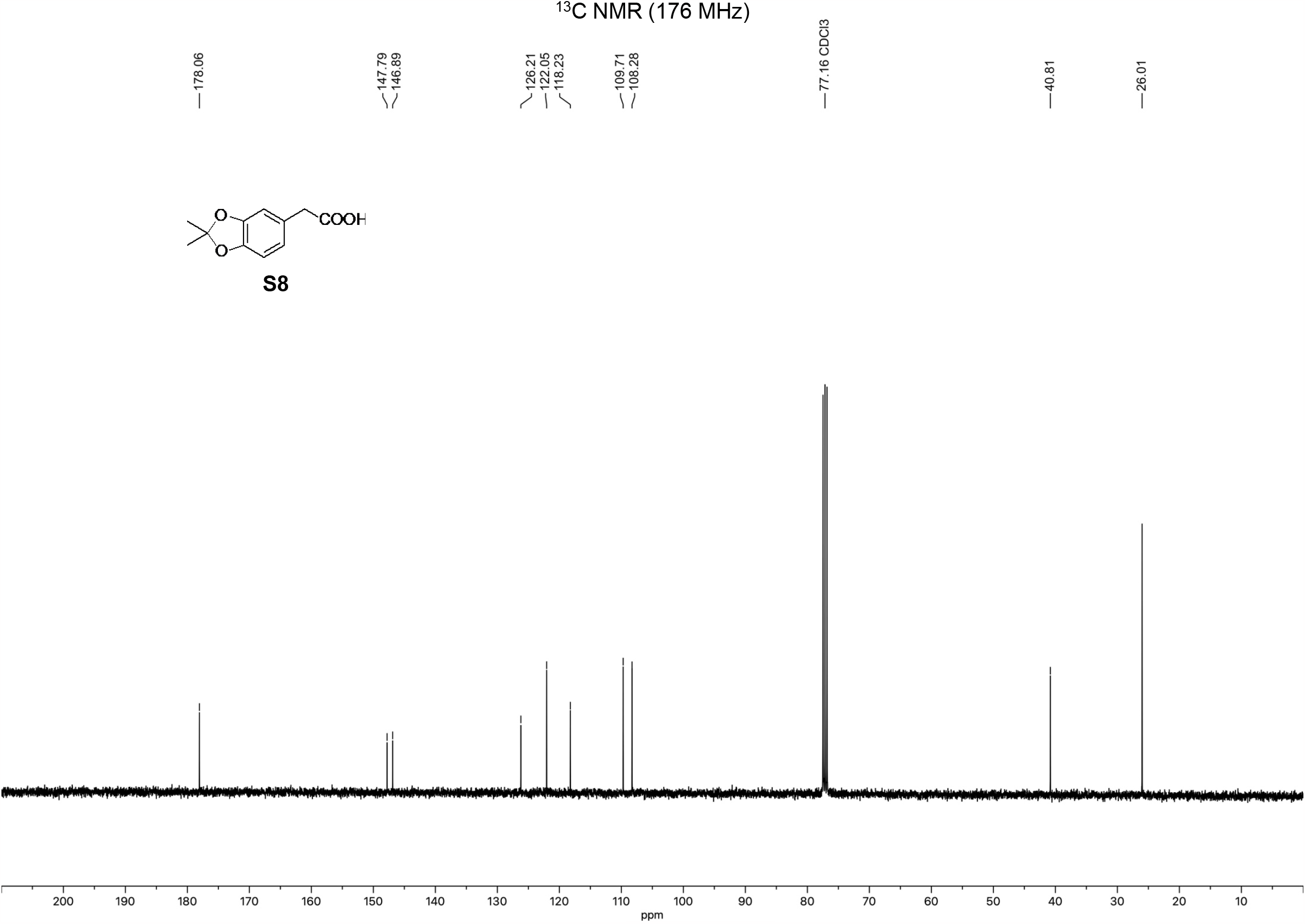

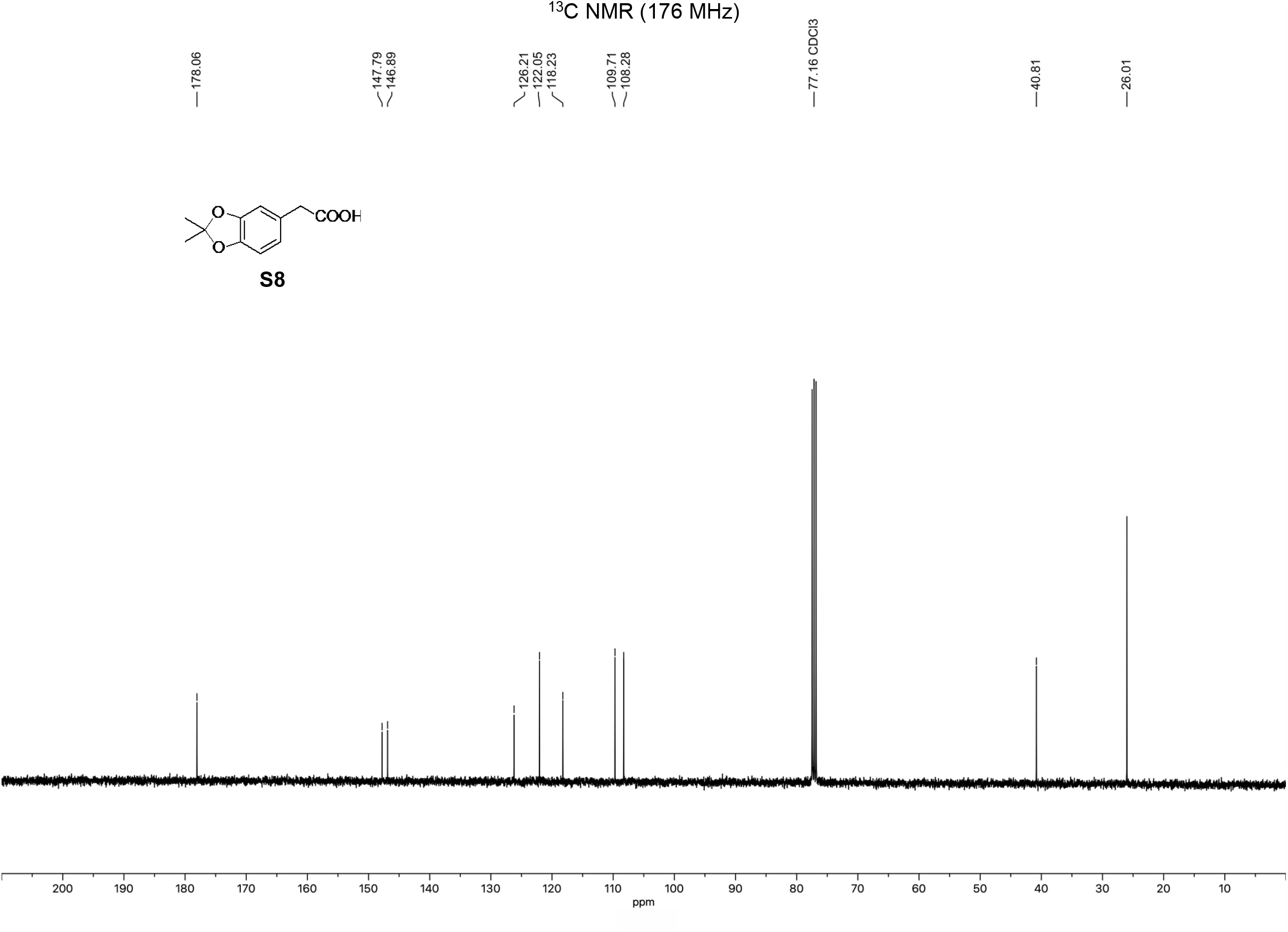

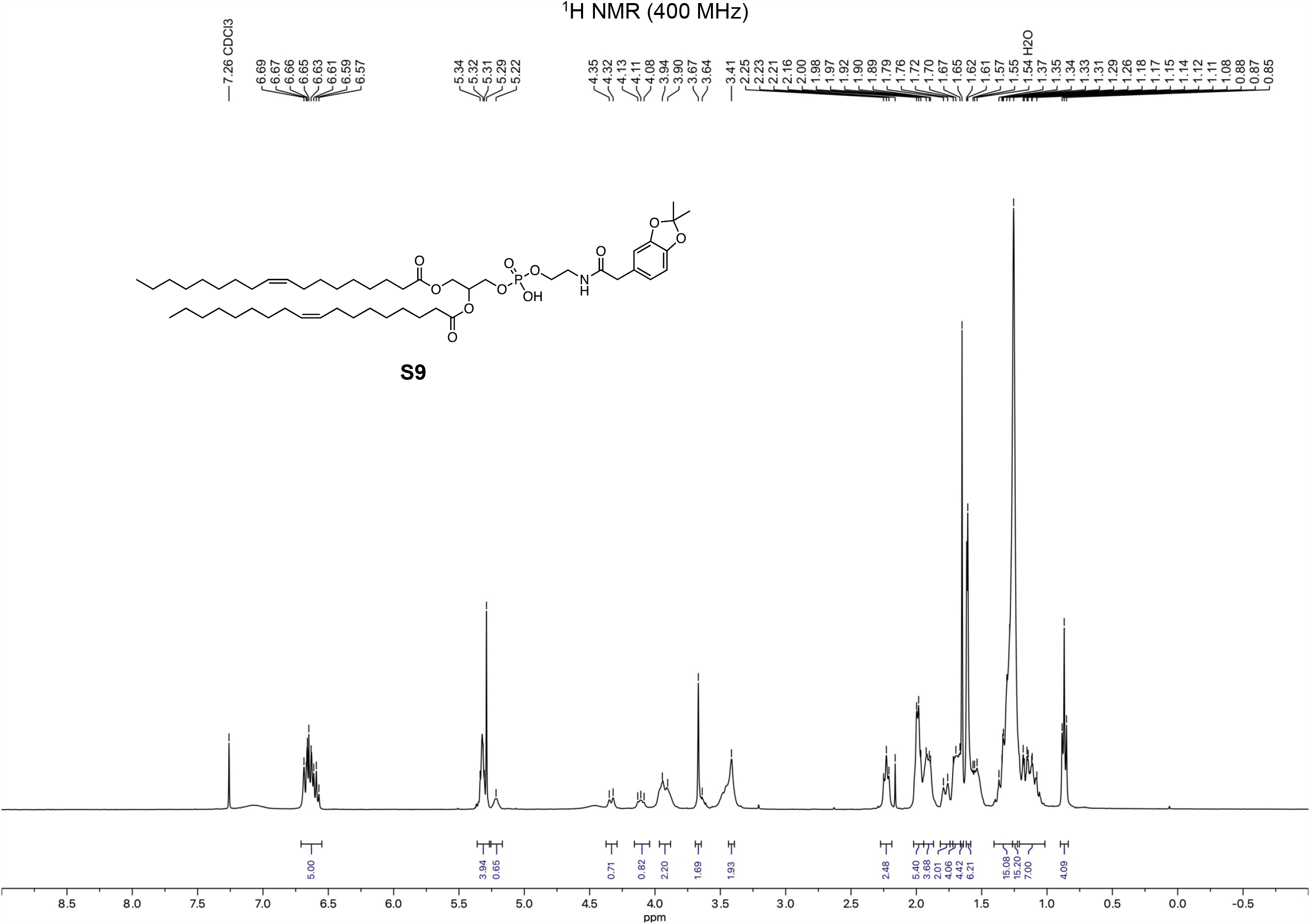

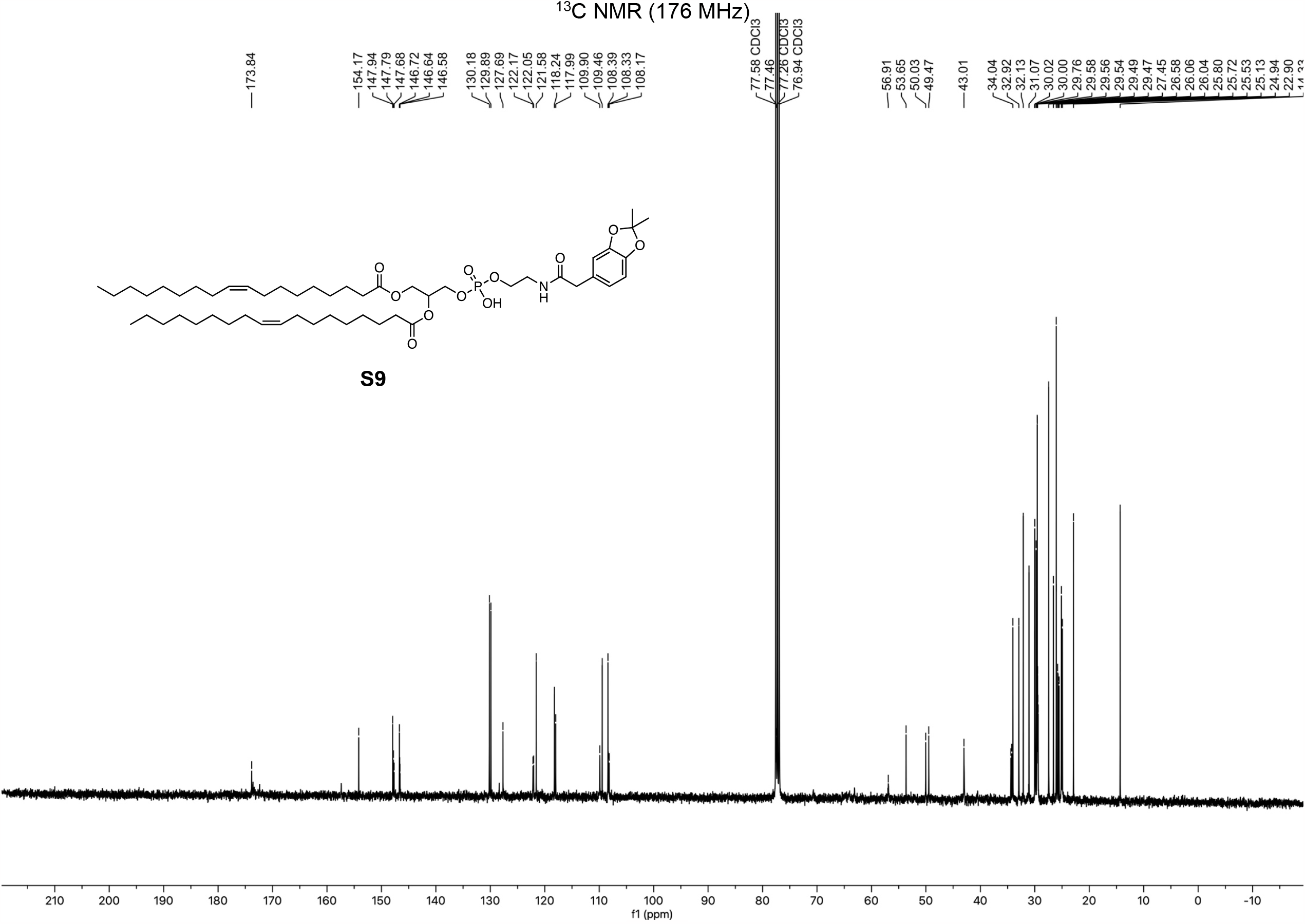

## Notes

### Competing Interest Statement

The authors have declared no competing interest.

